# Co-transcriptional production of programmable RNA condensates and synthetic organelles

**DOI:** 10.1101/2023.10.06.561174

**Authors:** Giacomo Fabrini, Nada Farag, Sabrina Pia Nuccio, Shiyi Li, Jaimie M. Stewart, Anli A. Tang, Reece McCoy, Róisín M. Owens, Paul W. K. Rothemund, Elisa Franco, Marco Di Antonio, Lorenzo Di Michele

**Affiliations:** Department of Chemistry, Imperial College London, 82 Wood Lane, London, W12 0BZ, UK; fabriCELL, Imperial College London, 82 Wood Lane, London, W12 0BZ, UK; Department of Chemical Engineering and Biotechnology, University of Cambridge, Philippa Fawcett Drive, Cambridge, CB3 0AS, UK; Department of Computing and Mathematical Sciences, California Institute of Technology, Pasadena, CA 91125, USA; Department of Bioengineering, University of California at Los Angeles, Los Angeles, CA 90024, USA; Department of Mechanical and Aerospace Engineering, University of California at Los Angeles, Los Angeles, CA 90024, USA

**Keywords:** Condensates, RNA, Nanotechnology, LLPS, Organelles, Synthetic Cells

## Abstract

Condensation of RNA and proteins is central to cellular functions, and the ability to program it would be valuable in synthetic biology and synthetic cell science. Here we introduce a modular platform for engineering synthetic RNA condensates from tailor-made, branched RNA nanostructures that fold and assemble co-transcriptionally. Up to three orthogonal condensates can form simultaneously and selectively accumulate guest molecules. The RNA condensates can be expressed within synthetic cells to produce membrane-less organelles with controlled number, size, morphology and composition, and that display the ability to selectively capture proteins. The *in situ* expression of programmable RNA condensates could underpin spatial organisation of functionalities in both biological and synthetic cells.

## 1 Introduction

Membrane-less compartmentalisation emerges from the condensation of proteins and RNA, and is recognised as a primary mechanism through which cells dynamically control biochemical processes [1–4]. By co-localising nucleic acids, enzymes and metabolites, membrane-less organelles (MLOs) such as nucleoli, Cajal bodies and stress granules are believed to regulate biogenesis, transcription, post-transcriptional modification and degradation, and generally improve cellular fitness [4–7]. The emergence of biomolecular condensates has also been linked to disease, particularly to neurodegeneration [8, 9].

The ability to express “designer condensates” with prescribed properties would be a valuable tool to program cellular behaviour [10, 11] and engineer synthetic cells [12]. Remarkable examples based on engineered peptides [10, 11] or condensate-forming RNA repeat sequences [9, 13, 14] and riboswitches [15, 16] have highlighted the feasibility of this concept. The generality of these strategies, however, is hampered by challenges linked to protein engineering and the limited programmability of natural RNA constructs.

Leveraging the algorithmic toolkit of nucleic acid nanotechnology [17–19], here we introduce a systematic method for expressing designer biomolecular condensates from synthetic RNA nanostructures. Our elementary motifs consist of star-shaped junctions, or nanostars, which fold co-transcriptionally and assemble thanks to programmable base-pairing interactions, forming liquid or gel-like condensates. Thanks to the selectivity of base pairing, up to three co-existing but fully distinct condensate types can be produced. Expressing the condensates in model synthetic cells generates MLOs with controlled size, number, morphology and composition. Finally, including RNA aptamers enables selective capture of small molecules and proteins, imitating the ability of natural MLOs to recruit clients.

Because the RNA nanostars are transcribed *in situ* from DNA templates, rather than assembled from pre-synthesised strands, our platform could be directly applied to engineering living cells, besides its immediate use for creating functional MLOs in synthetic cells. In this context, exploring the vast design space of RNA nanostars will allow for fine tuning of condensate properties and self-assembly behaviour, as shown by Stewart *et al*. [20].

## 2 Results

The four-armed RNA nanostars, shown in Fig. 1a, consist of a single RNA strand that folds co-transcriptionally into the intended shape. The star-shape was inspired by well characterised DNA nanostars [21–23]. However, the RNA motifs interact *via* self-complementary (palindromic) HIV-type Kissing-Loops (KLs) present at the end of each arm [24], rather than *via* the single-stranded (ss) overhangs or hydrophobic modifications adopted for DNA designs [21–23]. Similar KL interactions have been shown to facilitate condensation in bacterial riboswitches [15, 16]. We tested three RNA nanostar designs, labelled A, B and C, featuring mutually orthogonal KL sequences (see zoomed-in boxes in Fig. 1a). In designs A and B, one of the double-stranded RNA (dsRNA) arms includes a Fluorescent Light up Aptamer (FLAP): Malachite Green Aptamer (MGA) for A [25, 26] and Broccoli Aptamer (BrA) for B [27] (Figs 1a (i) and (ii)). FLAPs yield a fluorescent signal upon binding their cognate fluorophores (malachite green for MGA and DFHBI for BrA), enabling characterisation *via* fluorescence microscopy and fluorimetry. In design C, both MGA and BrA were included in non-adjacent arms (Fig. 1a (iii)). The fluorescence characteristics of all RNA constructs are reported in Fig. S1. The arms that do not host FLAPs were designed to be 25 base-pairs long, and separated by an unbound uracil residue at the junction to improve flexibility [28]. The motifs were designed to be transcribed with T7 RNA Polymerase (T7 RNAP) from fully double-stranded DNA (dsDNA) templates, labelled as A-T, B-T and C-T for designs A, B and C, respectively (see Fig. S2). Denaturing polyacrylamide gel electrophoresis confirms the expected electrophoretic mobility for most transcripts, with truncated and over-elongated products present in small amounts [29, 30] (Fig. S3). Native agarose gel electrophoresis suggest that transcripts retain the intended folded monomeric conformation rather than producing misfolded multimers connected by hybridisation of the arm domains, as reported for natural condensate-forming RNA constructs [15, 16] (Fig. S4). Details on sequence design and *in vitro* transcription protocols are reported in SI Methods, sections 1.1-1.3. All sequences are reported in Tables S1-S4.

**Fig. 1.**
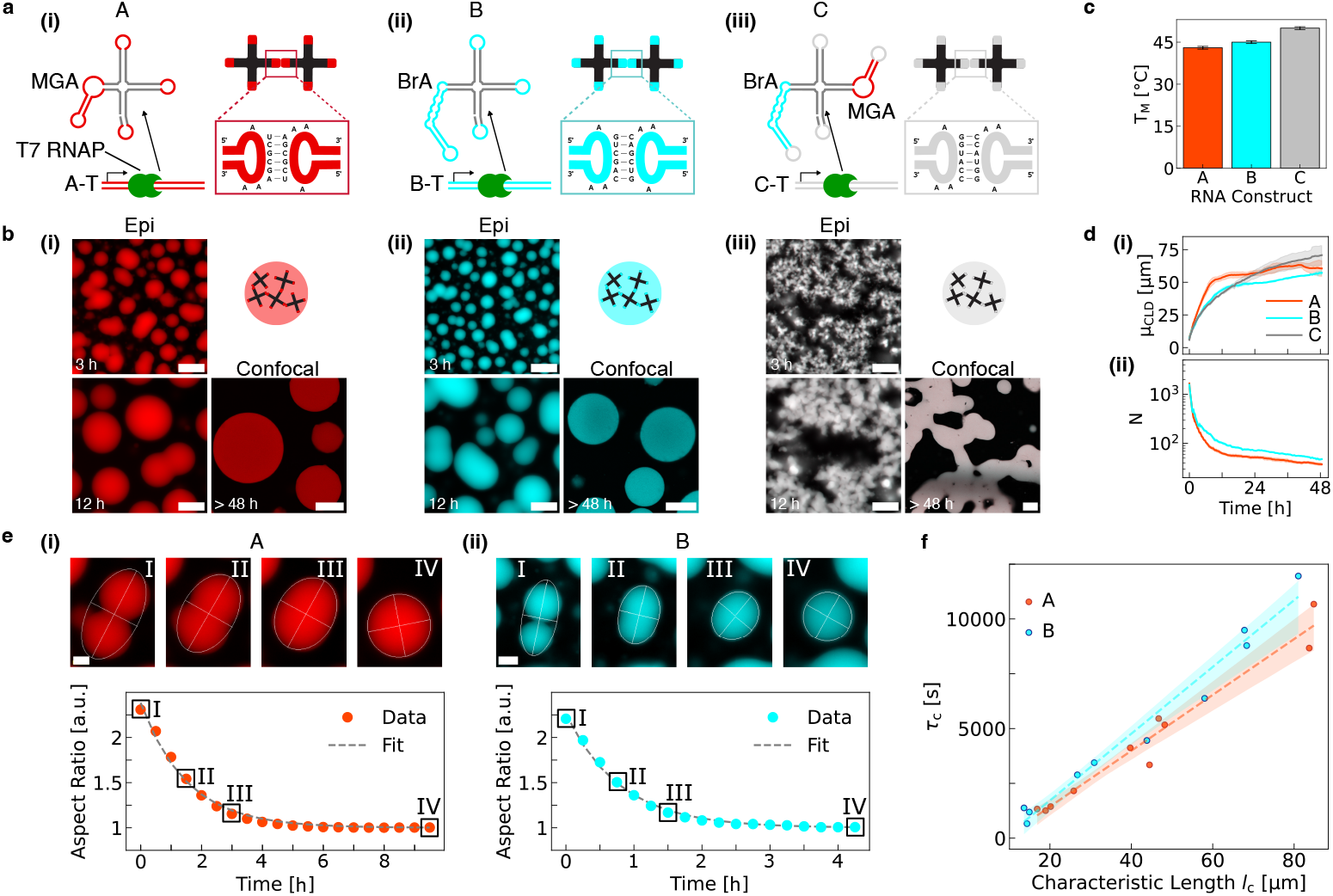
Condensation of co-transcriptionally folding RNA nanostars. (a) Structure of the RNA motifs. (i) A-type RNA nanostars include a Malachite Green Aptamer (MGA), (ii) B-type include a Broccoli Aptamer (BrA), and (iii) C-type include both aptamers. Variants feature mutually-orthogonal, self-complementary (palindromic) Kissing Loops (KLs), whose sequences are shown in the insets. RNA nanostars are transcribed from linear dsDNA templates by T7 RNA polymerase (T7 RNAP). (b) Epifluorescence and confocal micrographs showing condensate formation and coarsening for all three designs in (a) at different timepoints of an *in vitro* transcription reaction. Epifluorescence micrographs have been linearly re-scaled to enhance contrast (see SI Methods, section 1.4.2). Pristine micrographs are shown in Fig. S5. Scale bars are 50 *μ*m. Timestamps are reported with respect to the start of timelapse imaging (see SI Methods, section 1.4.1 and Table S5). (c) Melting temperatures of A-C condensates, determined as discussed in SI Methods, sections 1.4.1-1.4.2, and Figs S8 and S9. Errorbars (*±* 0.5 °C) stem from the discrete 1 °C step between consecutive measurements. (d) (i) Mean of the chord-length distribution, *μ*_CLD_, and (ii) average number of condensates per microscopy field-of-view, *N*, as a function of time. Full chord-length distributions, as extracted from image segmentation, are shown in Fig. S17 (see SI Methods, section 1.4.2). *N* is not computed for system C, which does not form discrete aggregates. Data are shown as mean (solid lines) *±* standard deviation (shaded regions) of three field-of-views within one sample. (e) Top: epifluorescence micrographs (contrast-enhanced) depicting coalescence events for (i) A and (ii) B condensates. Bottom: time-dependent aspect ratio of the condensates above, computed as the ratio between major and minor axes of the best-fit ellipse. The dashed line shows an exponential fit with decay constant *τ*_c_. (f) *τ*_c_ against the characteristic size (*l*_c_) of A and B condensates undergoing coalescence. Linear regression (dashed lines) yields inverse capillary velocities *μ/γ* = 127.4 s *μ*m^−1^ and 152.3 s *μ*m^−1^ for A and B condensates, respectively (SI Methods, section 1.4.2) [31].

As shown in Fig. 1b with representative epifluorescence and confocal snapshots, all three designs formed aggregates during *in vitro* transcription experiments. Additional examples are provided in Fig. S5, Fig. S6 (top) and video S1 (top). Variants A and B formed condensates that nucleated and grew, with frequent coalescence events, confirming that these designs yield liquid-like condensates. Lateral confocal projections reveal that the condensates were roughly spherical and did not significantly wet the glass substrate onto which they sedimented (Fig. S7), consistent with the evidence of Brownian motion seen in microscopy timelapses (video S1). Conversely, design C formed a gel-like percolating structure which failed to produce discrete condensates but still grew over time. The higher apparent viscosity of design C when compared to A and B correlates with the thermal stability of the materials. Indeed, C exhibited the highest melting temperature, followed by B and A (Figs 1c, S8 and S9, and video S2). Regardless of their morphology, all aggregates displayed the intended fluorescence output, namely in the MGA channel for A (red), BrA channel for B (cyan) and both channels for C (white).

Degradation of the condensates resulting in bubble formation was observed over time, ascribed to the action of environmental nucleases [32] or photo-degradation (Fig. S5). Bubbling was most prominent for A-type condensates, consistent with the lower melting temperature and consequent expectation that a lesser degree of RNA damage would be required to trigger disassembly. Sequence differences between the constructs may also result in different susceptibilities to enzymatic degradation.

The specificity of KL interactions was confirmed by non-sticky, control designs where KLs were replaced with scrambled sequences (Ā, 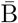). These designs did not yield condensates and only produced diffused fluorescence, as shown in Figs S6 (bottom) and S10, and video S1 (bottom).

The evidence that condensation requires KL complementarity suggests that the non-specific, cation-dependent, phase-separation mechanisms recently explored by Wadsworth *et al*. [33] are not dominant under the tested conditions. To further elucidate the role of the surrounding cationic makeup, we characterised the stability of A and B condensates upon buffer replacement, summarised in Fig. S11. Both condensate types remained largely stable after 24 hours in Tris-EDTA buffer supplemented with 5 mM or 10 mM MgCl_2_, but underwent disassembly when re-suspended in Phosphate-Buffered Saline, consistent with prior evidence that divalent cations stabilise KL interactions [34].

While not required in our system, crowding agents are often introduced to aid RNA condensation *in vitro* [35, 36]. We thus assessed the effect of crowding on cotranscriptional assembly of RNA nanostars by supplementing the reactions with 25% volume/volume poly-ethylene glycol (PEG) 200. While the formation of spherical condensates was still observed with both A and B systems, we noted a reduction in condensate size, consistent with prior observations that PEG 200 reduces the T7 RNAP transcription rate [37]. Increased viscosity induced by PEG may also slow down condensate growth and coalescence. Non-binding variants Ā and 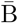 remained soluble in the presence of PEG, which is thus insufficient to trigger non-specific condensation.

Besides microscopy visualisation, aptamer fluorescence was also used to monitor the rate of synthesis of RNA building blocks with fluorimetry, as reported in Fig. S13. All designs displayed a rapid initial increase, whose rate scaled (nearly) linearly with template concentration (Figs S14 and S15). After an initial phase of rapid RNA production, synthesis slowed down significantly or plateaued, likely due to loss of polymerase activity and/or nucleotide depletion [38–40]. A peak was noted for condensate-forming motifs (A, B and C), which was not present in non-sticky designs (Ā and 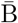), ascribed to sedimentation of the fluorescent aggregates below the measurement plane. Differences in plateauing behaviours could be due to variations in the kinetics of aptamer folding and/or complexation with the respective fluorophores [41– 43]. Fluorescence-intensity analysis of segmented epifluorescence timelapses reveals that, after an initial transient, the ratio between nanostar concentration in the bulk and within the condensates reached a plateau, indicative of a steady-state between the RNA concentration in the dilute and condensed phases (Fig. S16 and SI Methods 1.4.2).

We gained further insights on condensate growth and coarsening dynamics from the temporal evolution of the chord-length distribution (CLD), collated in Fig. S17 [44–46]. Chords were extracted from binarised epifluorescence images as the intersections between the condensates and straight lines (SI Methods, section 1.4.2). The CLD hence provides a snapshot of condensate length-scales in a way that is agnostic of their shape, and is thus equally applicable and meaningful for the branched structures formed by C-type nanostars and the compact A and B condensates.

Fig. 1d (i) shows the time evolution of the mean of the CLD – *μ*_*CLD*_ – useful as a proxy for the typical condensate size. For all designs, *μ*_*CLD*_ rapidly increased at early stages, likely sustained by the active transcription leading to monomer addition (Figs S13 and S16). For A and B designs, frequent coalescence events also contributed to the increase in *μ*_*CLD*_, and are reflected by a steep decrease in the number of A or B condensates *per* microscopy field-of-view (*N*, Fig. 1d (ii)). Coalescence appears to occur more readily in A compared to B, given the steeper decrease in *N* and increase in *μ*_*CLD*_ seen in the former. Video S1 suggests that early coalescence events may be driven by neighbouring condensates coming into contact as they grew through monomer addition, aided by Brownian motion. Consistently, both the size and number of A and B condensates plateaued when transcription slowed down (Fig. S13). In C aggregates, the increase of *μ*_*CLD*_ continued at later times, driven by the slow coarsening of the percolating RNA network (Fig. 1b (iii) and video S1).

The coalescence dynamics of A and B condensates can be further analysed to determine the inverse capillary velocity of the RNA phases, namely the ratio between their viscosity (*μ*) and surface tension (*γ*) [31, 47, 48] (see SI Methods, section 1.4.2). As summarised in Figs 1e and f we find *μ/γ* = 127.4 s *μ*m^−1^ and *μ/γ* = 152.4 s *μ*m^−1^ for A and B-type condensates, respectively. These values are significantly higher compared to those reported for DNA-nanostar condensates, ranging from 0.9 to 26.3 s *μ*m^−1^ [31, 47, 48]. For protein-based and biological condensates, reported inverse capillary velocities span several orders of magnitude, from ∼ 10^−2^ to ∼ 10^2^ s *μ*m^−1^ [49, 50], encompassing the values observed for RNA nanostar condensates. Fluorescence Recovery After Photobleaching (FRAP, SI Methods, section 1.4.4), performed using fluorescein dyes covalently linked to the RNA (SI Methods, section 1.3.5) revealed lack of recovery over *>* 500 s for both A and B designs (Fig. S18). Combined with the high *μ/γ* values, this evidence suggests that the RNA-nanostar condensates may be more viscous compared to their DNA counterparts [31, 47, 48]. When performing FRAP using the embedded FLAPs, both A and B condensates showed rapid fluorescence recovery, likely due to exchange of dyes with the bulk [43] (Fig. S18).

KL orthogonality enables the simultaneous transcription and assembly of designs A and B, that readily formed distinct, co-existing condensates, as shown in Fig. 2a (see also Figs S19, S20, and video S3). The absence of non-selective interactions between A and B condensates further confirms the negligible influence of base-pairing-independent condensation pathways [33]. Consistently, if one of the RNA motifs was rendered non-sticky, condensates of one species co-existed with dispersed RNA nanostars of the other, as shown in Figs 2b, S21, S22, and video S4.

**Fig. 2.**
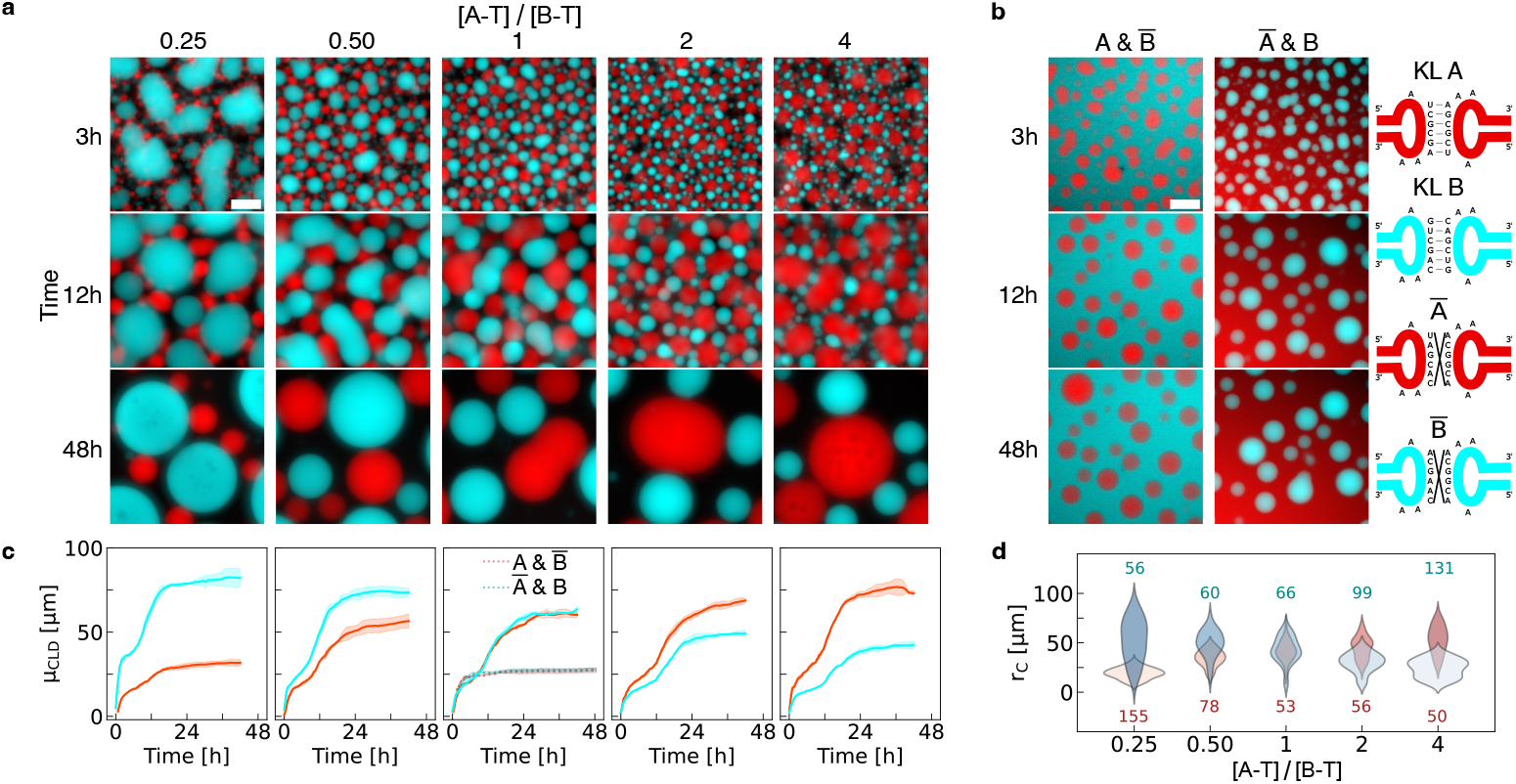
Co-transcribed orthogonal RNA nanostars form immiscible condensates of controlled size. (a) Epifluorescence micrographs of binary systems of A and B RNA nanostars (see Fig. 1) at various timepoints during the transcription transient. Different ratios between the concentrations of the two templates, A-T and B-T, are tested, while keeping [A-T]+[B-T] constant. (b) Epifluorescence snapshots analogous to panel (a), but where either A or B is replaced by its non-sticky variant, namely Ā or 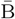. Sketches on the right hand side show examples of scrambled non-binding KL sequences. (c) Time evolution of the mean chord-length, *μ*_CLD_ (see SI Methods section 1.4.2), for samples in (a) and (b), the latter shown in the central panel as dashed lines. See Fig. S23 for full chord-length distributions. Red and cyan curves are relative to A and B condensates, respectively. Data are shown as mean (solid/dashed lines) *±* standard deviation (shaded regions) of three field-of-views within one sample. (d) Distribution of the radii, *r*_c_, of A (red) and B (cyan) condensates as a function of the template concentration ratio [A-T]/[B-T]. Epifluorescence micrographs in (a) and (b) have been linearly re-scaled to enhance contrast (see SI Methods, section 1.4.2). Pristine micrographs are shown in Figs S19 and S21. All scale bars are 50 *μ*m. Timestamps are reported with respect to the start of timelapse imaging (see SI Methods, section 1.4.1 and Table S5).

A simple strategy to control the relative size of A and B condensates is to tune the ratio between the concentrations of the corresponding DNA templates ([A-T] and [B-T], respectively), which determines the production rate of each nanostar, as demonstrated in Figs S14 and S15. Relative-size control is demonstrated visually in Fig. 2a (see also Figs S19 and S20), through time-dependent *μ*_*CLD*_ analysis in Fig. 2c (see Fig. S23 for full CLDs), and through the distribution of the final condensate radii, *r*_c_, shown in Fig. 2d (see SI Methods, section 1.4.2). We observed condensate number to anti-correlate with their size. For instance, many small A-type condensates were formed when [B-T]*>*[A-T] (see Fig. S24).

As shown in Fig. 2c, condensates were found to grow in two stages in all binary systems of sticky nanostars: after an initial increase, a brief intermediate plateau was reached, followed by another growth phase before saturation. This behaviour was not observed in single component systems (Fig. 1d), nor in binary systems where one nanostar population was rendered non-sticky, for which the temporal evolution of *μ*_*CLD*_ is included in the central panel of Fig. 2c. Inspection of the relevant microscopy videos (videos S3 and S4) reveals that the intermediate size plateau was reached when condensates became temporarily hindered in their ability to make contact with same-type assemblies, due to being “caged” by nearest-neighbours condensates of the opposite type. Coalescence events that still managed to occur, however, reduced lateral crowding given that merged condensates occupy less space in the horizontal plane, triggering a positive-feedback process that rapidly unjammed the system and accelerated coalescence. In fact, Fig. 2c shows that condensates reached a larger overall size in systems where both A and B were sticky, compared to Ā & B and A & 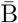 mixtures (Figs 2b and S23), indicating that steric encumbrance from non-binding condensates of the opposite phase ultimately facilitates coalescence. Two-step decreasing trends, consistent to the two-step growth seen in *μ*_*CLD*_, were observed in the number of condensates *per* field-of-view (Fig. S24).

As shown in Fig. S12, addition of 25% PEG induced non-specific affinity between A and B condensates, likely due to excluded-volume effects [51, 52]. As a result, the binary system formed extended networks of small alternating A- and B-type condensates, reminiscent of colloidal gelation [53].

Condensate co-assembly is also possible with all three RNA species A, B, and C, as shown in Figs S25, S26 and video S5. After allowing sufficient time for relaxation, all three species formed distinct spherical condensates, including C, which was unable to do so in single-component samples (Fig. 1b (iii)). The difference in morphology is likely due to A and B stars hindering the formation of a percolating C network in favour of smaller aggregates that relax more readily.

Addressable self-assembled RNA condensates could be extremely valuable to engineer compartmentalisation in synthetic or living cells, where they could operate as membrane-less organelles capable of recruiting target compounds and underpinning spatial separation of functionalities. To demonstrate this, we transcribed our condensates within basic synthetic cells constructed from water-in-oil emulsion droplets (Fig. 3a). All designs formed condensates that equilibrated into a single spherical organelle in each synthetic cell (Figs 3b and S27), including design C that generated only extended networks when produced in bulk. The different morphology is readily rationalised by noting that, when confined to the synthetic cell, C aggregates only need to relax over much smaller length-scales. Yet, shape relaxation was slower for C compared to A and B, as confirmed by the time evolution of *μ*_CLD_ in Fig. 3c (see also micrographs in Fig. S28 (top) and full CLDs in Fig. S29) and video S6 (top). Polydispersity in condensate size reflects the variability in the size of droplet-based synthetic cells produced *via* emulsion methods. We indeed observed that the final volume of the RNA condensates scales linearly with the volume of the surrounding emulsion droplet, as presented in Fig. 3d.

**Fig. 3.**
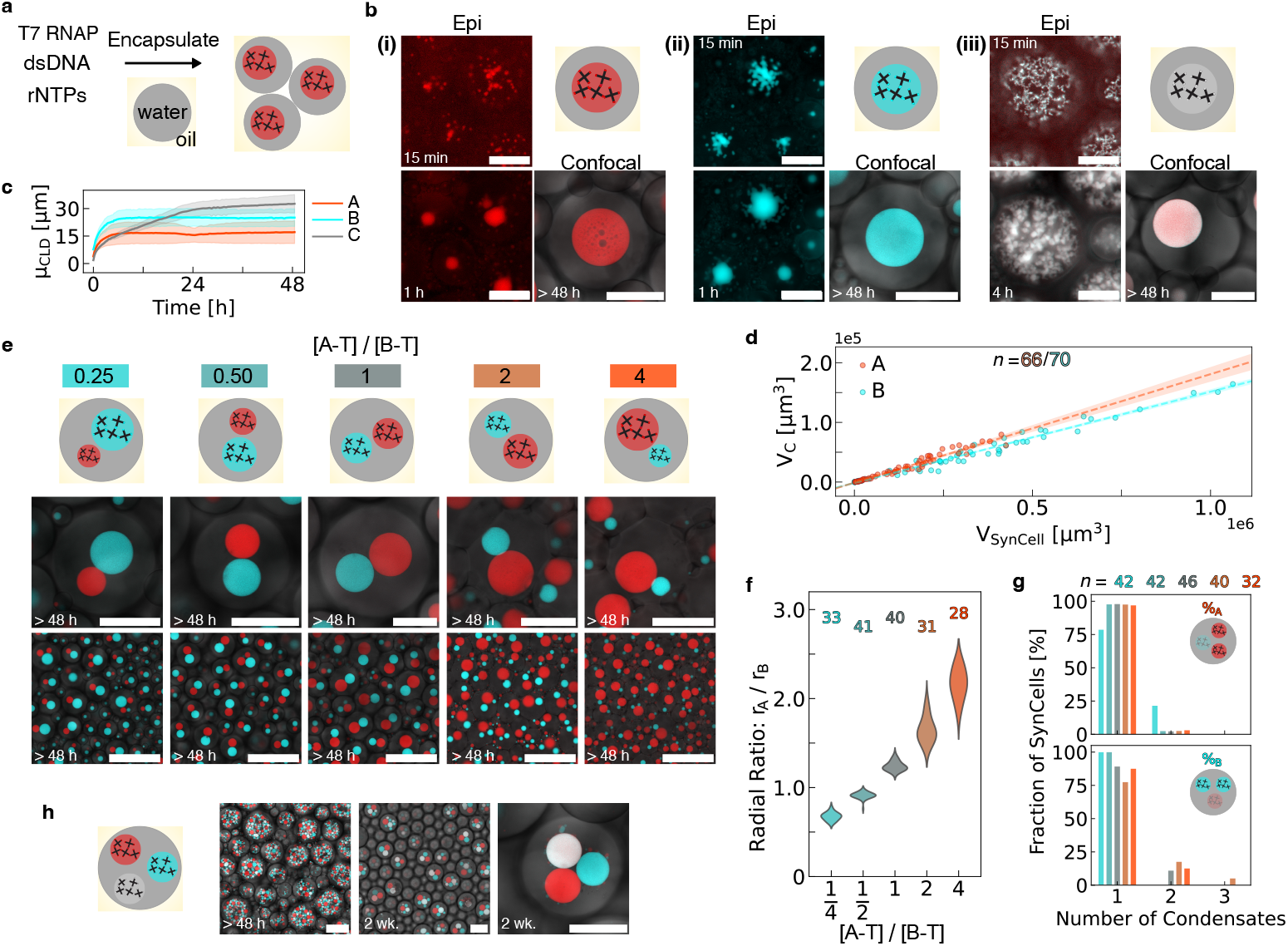
Membrane-less RNA organelles expressed in synthetic cells. (a) Diagram showing membrane-less organelles (MLOs) formed in synthetic cells consisting of water-in-oil emulsion droplets encapsulating transcription machinery and DNA templates. (b) Epifluorescence and confocal micrographs showing MLO formation over time in synthetic cells expressing (i) A-type, (ii) B-type, and (iii) C-type RNA nanostars (see Fig. 1). Epifluorescence micrographs have been linearly re-scaled to enhance contrast (see SI Methods, section 1.4.2). Pristine images are shown in Fig. S27, alongside images relative to additional timepoints. Timestamps are reported with respect to the start of timelapse imaging (see SI Methods, section 1.4.1 and Table S6). (c) Time-dependent mean of the chord-length distribution, *μ*_CLD_, computed as discussed in SI Methods section 1.4.2. Data are shown as mean (solid line) *±* standard deviation (shaded region) from three fields-of-view within one sample. (d) Scatter plot of condensate volume (*V*_C_) *vs* synthetic cell volume (*V*_SynCell_) for samples in (b)-(i) and (b)-(ii). Dashed lines indicate best fits to linear regression models, with 95% confidence intervals shown as shaded regions (SI Methods, section 1.4.2). MLOs occupy 18.2 *±* 0.5% and 15.3 *±* 0.3% of the volume of the synthetic cells for A and B systems, respectively. (e) Zoomed-in (top) and larger field-of-view (bottom) confocal micrographs depicting synthetic cells co-expressing A and B-type condensates, with different template concentration ratios [A-T]/[B-T] (compare Fig 2a). (f) Distribution of the ratio between the radii of A and B MLOs (*r*_A_*/r*_B_) as a function of [A-T]/[B-T] for samples in (e) (see SI Methods). (g) Percentage of synthetic cells containing a given number of A-type (top) or B-type (bottom) MLOs. The percentages of synthetic cells containing exactly one A and one B MLOs are 78.57%, 97.62%, 86.96%, 77.50%, 87.50% for [A-T]/[B-T] = 0.25, 0.50, 1, 2, 4, respectively. Colour-codes in panels (f) and (g) match those in (e). Numbers in (f) and (g) indicate sampled synthetic cells. (h) Confocal micrographs showing synthetic cells expressing three orthogonal MLO-forming RNA nanostars (A, B and C in Fig. 1) at different timepoints. Scale bars in panel (e), bottom, and panel (h), left and centre, are 150 *μ*m. All other scale bars are 50 *μ*m.

Control experiments with non-sticky designs revealed uniform fluorescence within the synthetic cells, confirming that the assembly specificity previously noted in bulk samples persists in confinement (Figs S28 (bottom), S30, and video S6 (bottom)). Like in bulk experiments, fluorimetry can be used to monitor RNA synthesis rates, as shown in Fig. S31, where the delayed growth in the MGA signal (A component) is due to initial accumulation of the malachite green fluorophore in the oil phase [54, 55], rather than to a slower growth of the condensates (compare with Fig. 3c). Also consistent with trends recorded in the bulk, initial transcription rates as determined *via* fluorimetry were found to scale (nearly) linearly with template concentration (Figs S32 and S33).

The condensate volume fraction determined in Fig. 3d can be used to calibrate fluorimetry measurements and estimate the time-dependent concentration of B-type RNA nanostars during the transcription transient, as outlined in the SI Methods, section 1.5. Data in Fig. S34 show that the nanostar concentration exceeded 10 *μ*M (or ∼ 1 g L^−1^) within 1.5 hours of the start of transcription, corresponding to the 15-minute timestamped micrograph in Fig. 3b (ii) (Table S6). The observation of aggregation at these early stages is consistent with reports on DNA nanostar constructs, displaying phase separation at concentrations as low as 0.25 *μ*M [41] or ≤ 0.1 g L^−1^ [56].

We obtained synthetic cells with two distinct synthetic organelles by encapsulating both A-T and B-T templates, as shown in Figs 3e, S35-S39, and videos S7-S8 with microscopy, and Fig. S40 with fluorimetry. Like in the bulk experiments, we could control the relative size of the organelles by changing the template ratio, as quantified in Fig. 3f for synthetic cells containing exactly one A and one B condensate. In the vast majority of cases, in fact, each synthetic cell contained one condensate of each type (Fig. 3g). When including component C, we obtained 3 distinct phases, as demonstrated by microscopy in Fig. 3h (see also Figs S41, S42 and, video S9), and fluorimetry in Fig. S43, exemplifying the possibility for scaling up the number of addressable organelles. Consistent with bulk experiments (Figs S25 and S26), relaxation timescales were significantly slower for the ternary system compared to binary ones (Figs 3e and S41). It is also evident that the synthetic cells often failed to produce exactly three distinct MLOs, likely due to steric effects and the intrinsic slow relaxation of phase C. When replacing component C with 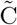, which features identical KLs but lacks any FLAPs, synthetic cells with exactly three MLOs were more common (Fig. S42), suggesting that the presence of the aptamers has an effect on the coarsening kinetics of the material.

We further expanded the possible organelle architectures in the synthetic cells by introducing linker RNA nanostars, dubbed L, that could control the mixing between the A and B designs upon their simultaneous transcription. Similarly to the strategy demonstrated by Jeon *et al*. with DNA constructs [57], nanostar L is “chimeric”, as it features two A-type and two B-type KLs (Fig. 4a). As shown in Fig. 4b, low fractions of the linker template ([A-T]:[L-T]:[B-T]=10:1:10, or Linker Template Fraction (LTF) = 1/21) produced grape-like clusters by inducing adhesion between condensates rich in A and B, while blocking their relaxation into two large condensates (see also Figs S44-S46). Arrested coarsening is arguably due to inter-phase adhesion limiting the ability of the condensates to slide past each other, similarly to what we observed for linker-free A-B systems in the presence of crowding agents (Fig. S12). At higher LTFs, the two opposite-type droplets coarsened into bigger domains, with Janus-like morphologies emerging at [A-T]:[L-T]:[B-T] = 5:1:5 (LTF = 1/11) (see also Fig. S46 and videos S10, S11). Here, we note the occasional formation of a cavity in the contact region between A-rich and B-rich phases, hinting at an uneven linker distribution. For [A-T]:[L-T]:[B-T] = 3:1:3 (LTF = 1/7) we observe hollow, capsule-like organelles for most of the larger synthetic cells, as shown in videos S11 and S12 (see also Fig. S46). Composition [A-T]:[L-T]:[B-T] = 2:1:2 (LTF = 1/5) produced Russian-doll morphologies with the A-rich phase forming the outer shell, while single phase spherical condensates were observed for LTFs of 1/3 and above.

**Fig. 4.**
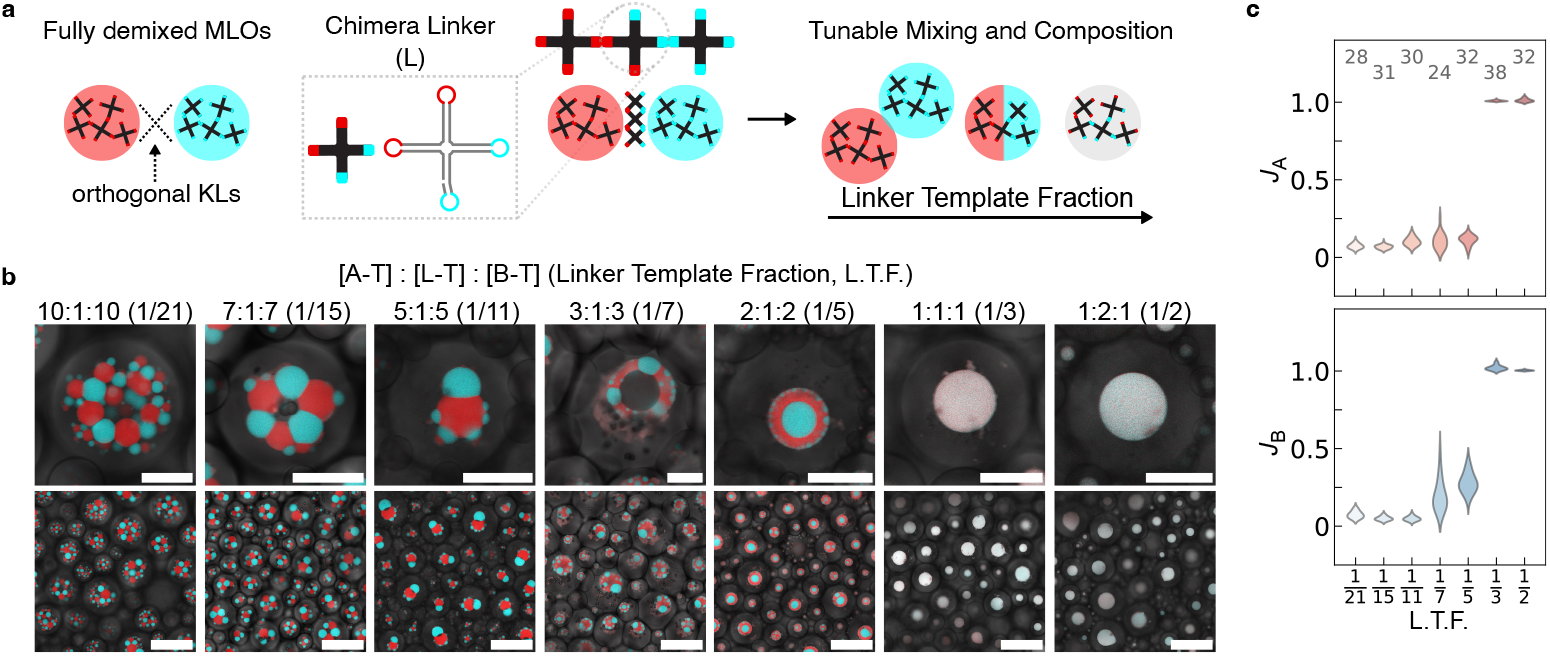
Controlling morphology and composition of membrane-less organelles with linker RNA nanostars. (a) Chimeric RNA linker nanostars (L), with two A and two B KLs, enable control over mixing in systems of A and B nanostars by varying the relative concentration of the linker template L-T. (b) Zoomed-in (top) and larger field-of-view (bottom) confocal micrographs, acquired after more than 48 h, depicting synthetic cells producing A, B and L nanostars with different ratios between DNA templates ([A-T]:[L-T]:[B-T]). The Linker Template Fraction (LTF), shown in brackets, is computed as [L-T]/([A-T]+[L-T]+[B-T]). For LTF = 1/3 and 1/2, slight changes in condensate colour occur away from the confocal imaging plane, likely due to differences in the extinction coefficients of Malachite Green and DFHBI. All scale bars are 50 *μ*m. (c) Distributions of mixing indices *J*_A_ and *J*_B_ of the MLOs computed as discussed in the SI Methods (section 1.4.3) and shown as a function of LTF for samples in (b). Low *J*_A_ and *J*_B_ are indicative of purer A-rich and B-rich phases, while *J*_A_, *J*_B_ ≈ 1 indicate complete mixing of the two RNA species. Numbers indicate examined synthetic cells.

The trends observed in condensate morphology are broadly consistent with a decrease in the interfacial tension between A- and B-rich phases (*γ*_AB_) with increasing LTF, as observed by Jeon *et al*. for equilibrium assembly of DNA nanostars [57]. Adhering de-mixed droplets (LTFs = 1*/*21 to 1*/*11), are indeed expected at equilibrium when *γ*_AB_ ∼ *γ*_A_ ∼ *γ*_B_, where *γ*_A_ (*γ*_B_) is the interfacial tension between the A-rich (B-rich) phase and the surrounding buffer [2]. The Russian-doll morphology (LTF = 1/5) should emerge for *γ*_A_ *< γ*_AB_ *< γ*_B_, with the evidence that *γ*_A_ *< γ*_B_ being consistent with trends seen in melting temperatures (Fig. 1c), while full mixing (LTF ≥ 1/3) should occur when *γ*_AB_ ∼ 0 [57]. However, we note that some morphological features, including the cavities seen for LTF = 1/11 and 1/7 and the outer layer of small B-rich domains seen for LTF = 1/5, are not expected at equilibrium, hinting that these may constitute metastable states emerging from isothermal cotranscriptional assembly.

Lateral confocal projections reveal that the loosely packed clusters formed at LTF = 1/21 had a curved morphology, ascribed to sedimentation within the spherical droplet container (Fig. S46). In all other conditions, the more compact or spherical condensates appeared unaffected by substrate curvature.

The abundance of linker nanostars also influences the degree of mixing of the two phases, assessed by measuring indices *J*_A_ and *J*_B_, computed as the ratios between intensity of the fluorescent signal from the minority RNA component (*e*.*g*. signal from component A in B-rich condensates) over that recorded for the intensity of the majority phase (*e*.*g*. signal from A in A-rich condensates), as outlined in the SI Methods (section 1.4.3). The data, collated in Fig. 4c, show limited mixing (*J*_A_, *J*_B_ ≪ 1) for low LTF, followed by moderate increase in mixing when more linker was produced, and by an abrupt jump to *J*_A_, *J*_B_ ≈ 1 upon reaching the threshold for complete mixing. A similarly sharp mixing transition was noted for annealed DNA nanostars by Gong *et al*., remarkably occurring at similar linker fractions when tetravalent linkers were considered [58].

While the data in Figs 1-4 demonstrate that RNA condensates can selectively sequester small molecules, namely the fluorophores associated to MGA and BrA, imitating the functions of natural MLOs requires capturing larger and more functionally relevant macromolecules, particularly proteins. To this end, we modified designs A and B to include a 5^*′*^ overhang, to which a protein-binding RNA aptamer can connect *via* base pairing (Fig. 5a (i) and (ii)). In particular, the new nanostructures A_YFP_ and B_STV_ were designed to connect to a YFP-binding aptamer (YFP_apt_) [59], and a streptavidin (STV)-binding aptamer (STV_apt_) [60], respectively (see also Fig. S47). For both designs, when co-expressing the modified nanostars and their partner aptamers from distinct templates in synthetic cells (Fig. S48), the target proteins (EYFP and Alexa405-STV) readily partitioned within the formed MLOs (Figs 5b (i) and (ii), S49, S50, and video S13). If the aptamer template was omitted, the target proteins remained in the lumen of the droplet-based synthetic cell. Protein partitioning was quantified through the parameter *ξ*, computed from image segmentation as the ratio between the fluorescence intensity of the protein recorded within or outside the MLOs for individual synthetic cells (SI Methods, section 1.4.3). When protein-binding aptamers were present, the median *ξ* was ∼ 3.5 and ∼ 2 for YFP and STV, respectively, while it fell below 0.5 when no aptamers were present (Fig. 5c (i) and (ii)). The significant anti-partitioning noted in the absence of aptamers is likely a consequence of excluded volume interactions between the RNA duplexes and the proteins. Indeed, the condensate mesh size, estimated as twice the RNA nanostar arm length (∼ 15.7 nm, see SI Methods, section 1.5), is comparable with the hydrodynamic diameters of the both STV (6.4 nm) and EYFP (5 nm), as estimated with the HullRad software tool [61]. Both STV and EYFP are reported to have mildly acidic to neutral isoelectric points [62, 63], hence Coulomb repulsion towards the RNA backbones may also enhance anti-partitioning.

**Fig. 5.**
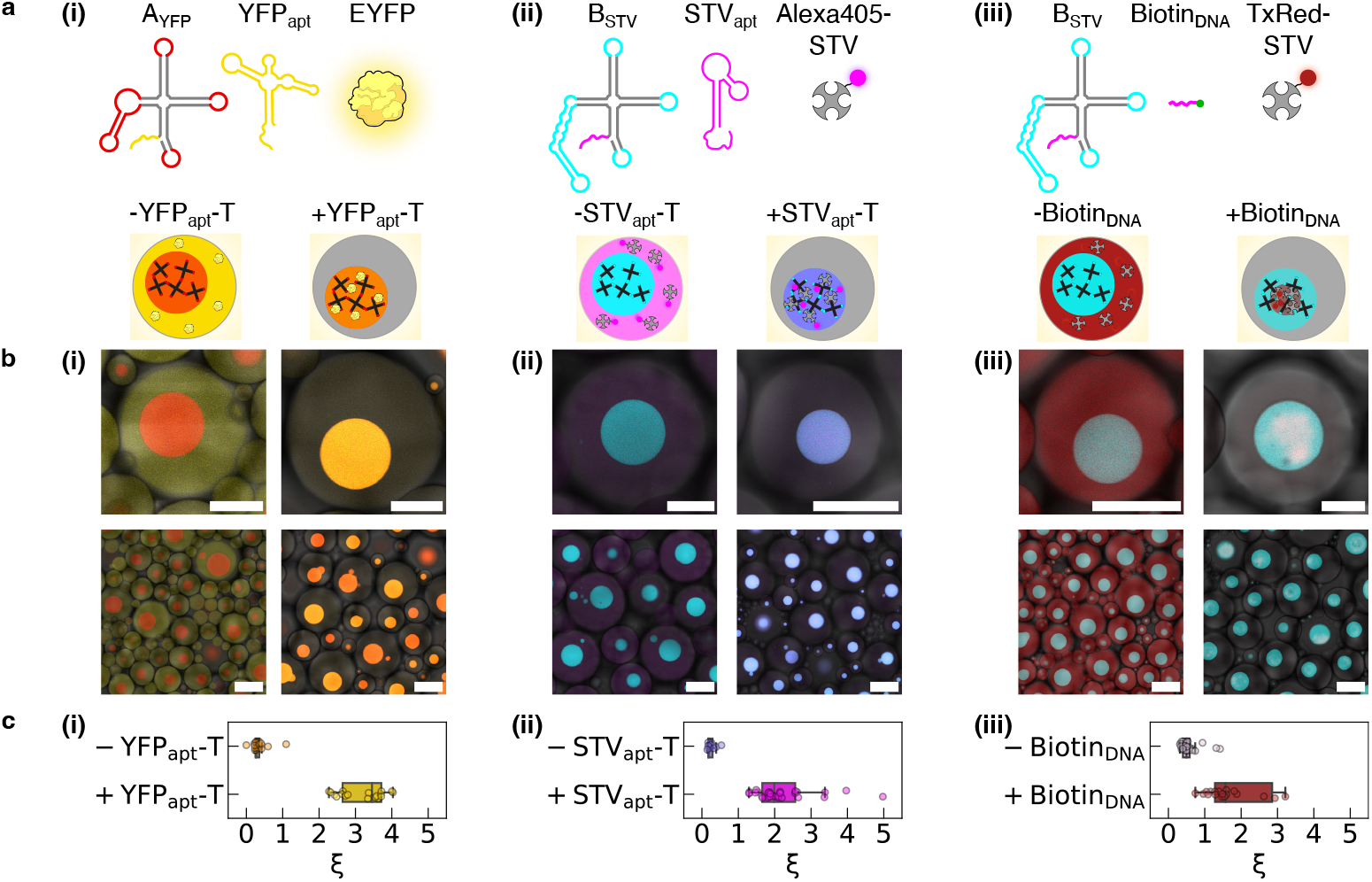
Selective protein capture in designer RNA MLOs. (a) RNA nanostar designs are modified to include a single-stranded 5^*′*^ overhang that can connect to a protein-binding moiety. Design A_YFP_ is identical to nanostar A (Fig. 1) but can connect to YFP-binding aptamer (YFP_apt_) (i). Design B_STV_ is identical to B but can connect to either a streptavidin (STV)-binding aptamer (STV_apt_) (ii) or a biotinylated DNA oligonucleotide (Biotin_DNA_) (iii). The templates of protein-binding aptamers (YFP_apt_-T and STV_apt_-T) are transcribed in synthetic cells alongside the associated RNA nanostar templates (A_YFP_-T and B_STV_-T, respectively), while pre-synthesised Biotin_DNA_ is encapsulated alongside template B_STV_-T. (b) Diagrams (top) and confocal micrographs (bottom) of synthetic cells expressing RNA MLOs in the absence (left) or in the presence (right) of protein-binding moieties for the systems presented in (a). When protein-binding aptamers are expressed or the biotinylated DNA oligonucleotide is included, target proteins are recruited in the MLOs. Scale bars are 50 *μ*m (top) and 150 *μ*m (bottom). (c) Protein partitioning parameter, *ξ*, computed from confocal micrographs in (b) as the ratio between the fluorescence signal of the target proteins recorded within and outside the MLOs for individual synthetic cells (see SI Methods, section 1.4.3). Data are shown for the three systems in (a) and (b), with and without protein-binding moieties. In the box plots, the central line marks the median, the box marks the Q1-Q3 interquartile range (IRQ) and the whiskers enclose datapoints within Q1−1.5×IQR and Q3+1.5×IQR. All data-points are shown except for four outliers with *ξ >* 5 in (i) and (iii), omitted for ease of visualisation.

As an alternative strategy for STV capture, we replaced STV_apt_ with a biotinylated DNA oligonucleotide complementary to the overhang in nanostar B_STV_ (Figs 5a (iii), S49, S50 and video S13). This approach still enables B-type MLOs to capture STV (Figs 5b (iii) and c (iii)), with the difference that the protein was distributed non-uniformly within the RNA condensates. Indeed, we observed irregularly-shaped clusters with solid-like appearance, reminiscent of multi-phase condensates found in cells [64–66]. The non-uniform protein distribution likely results from the finite amount of the biotinylated DNA anchor available, which is all sequestered at early transcription stages and thus accumulates at the centre of the condensates. The solid-like look of the protein-rich material may be a consequence of the tetravalent STV, which can irreversibly crosslink up to four RNA nanostars and thus make the material more viscous.

## 3 Discussion

Our platform enables the expression of synthetic condensates and membrane-less organelles with prescribed size, number, morphology, composition, and the ability to capture small guest molecules and proteins. The elementary building blocks, RNA nanostars, were designed from first principles utilising the rule-based approaches of nucleic acid nanotechnology, which provide extensive opportunities for further design variations aimed at programming arbitrary characteristics of the designer MLOs. Among the many design features that can be straightforwardly controlled are nanostar valency, flexibility and arm-length, all known to predictably influence self-assembly in analogous DNA systems [21, 23, 28, 41, 67]. Control over condensate-size could be achieved by co-transcribing monovalent, surface passivating RNA constructs [68, 69]. Further RNA aptamers might be embedded to recruit molecular guests, including enzymes and metabolites, while ribozymes [70] could confer catalytic properties to the synthetic MLOs. Because the condensates can be expressed from DNA templates, their formation could be controlled through standard transcription regulation pathways in both synthetic cells and living cells. Owing to their open-ended programmability, we expect that the RNA-nanostar condensates will constitute a valuable new solution for the toolkit of synthetic biology.

## Supplementary information

Supplementary information available: Methods, Figures, Tables and Videos.

## Data availability

Raw data underpinning these results are available from the corresponding author during the review process. Upon possible publication the data will be shared on a freely accessible permanent repository, for which the DOI link will be provided.

## Code availability

All data analysis code is available from the corresponding author.

## Acknowledgments

LDM and NF acknowledge support from the European Research Council (ERC) under the Horizon 2020 Research and Innovation Programme (ERC-STG No 851667 – NANOCELL) and a Royal Society University Research Fellowship (UF160152, URF*\*R*\*221009). GF acknowledges funding from the Department of Chemistry at Imperial College London. MDA acknowledges support from a Biotechnology and Biological Sciences Research Council (BBSRC) David Phillips Fellowship (BB/R011605/1) and a Lister Institute Research Prize. SPN acknowledges support from the Engineering and Physical Sciences Research Council (EPSRC) (EP/S023518/1). JMS is a Merck Awardee of the Life Sciences Research Foundation. EF acknowledges support from the US NSF through CAREER award 1938194 and FMRG: Bio award 2134772, and from the Sloan Foundation through award G-2021-16831. RM acknowledges funding from the EPSRC Centre for Doctoral Training in Nanoscience and Nanotechnology (NanoCDT, EP/S022953/1). GF, LDM and MDA acknowledge the Facility for Imaging by Light Microscopy (FILM) at Imperial College London and thank Stephen Rothery for his invaluable assistance and the publicly released FIJI macros he developed.

