## Supplementary Information for "Co-transcriptional production of programmable RNA condensates and synthetic organelles"

### 1 Materials and Methods

#### 1.1 Sequence design

Four-armed RNA nanostars with 25 base pair (bp)-long arms separated by one unpaired uracil residue were designed starting from the sequences of previously reported four-way RNA junctions [1] and fluorogenic Malachite Green [2, 3] and Broccoli [4] RNA aptamers.

Three main nanostar variants, namely A, B and C, were designed to bind identical motifs, while being non-interacting towards each other. Intra-population interactions were guaranteed by palindromic Kissing Loops (KLs). KL A (5'-AUCGCGAAA-3') was adapted from the KL domain in the bottom right arm of the T4 tetrahedron in Ref. [1], by making it palindromic. Asymmetrical flanking bases (5'-A. . . AA-3') were introduced due to their presence, albeit in reverse order, in the original KL domains found in the Lai variant of the Human Immunodeficiency Virus-1 Dimerisation Initiation Sequence (HIV-1 DIS) [5, 6]. KLs B and C were designed to match the interaction strength of KL A, while maximising orthogonality towards it and each other. This was achieved by computationally generating, using Python3, all possible palindromic, 6 nucleotide (nt)-long sequences with the same GC content as KL A. The resulting set was filtered to exclude all sequences with more than two overlapping nucleotides with KL A. Among these, KL C was selected as 5'-AGGUACCAA-3'. To determine KL B, the constraint was relaxed to allow a maximum of 3 overlapping nucleotides with both KL A and KL C. This produced several candidates, among which KL B was chosen as 5'-AGUCGACAA-3'. We note that this sequence is similar to the Mal variant of the HIV-1 DIS KL (5'-GUGCAC-3') [5].

Minimum free energy configuration of all designs was evaluated using NUPACK (default Serra and Turner, 1995 parameters and 1 M $Na^{+}$) [7]. Kinefold [8] was used to test co-transcriptional folding. In particular, all designs were tested *via* batch jobs, and further considered only if the helix tracing graph showed correct folding order and reasonable stability of each helix. All sequences are provided in Table S1. Sequences for the coding/non-template DNA strands were obtained by adding a prefix comprising the T7 promoter (5'-TTCTAATACGACTCACTATA-3', with the 17 nt consensus T7 promoter underlined) to the equivalent DNA sequence of the designed RNA nanostructures. This 'extended' promoter was taken from Ref. [1]. DNA templates are labelled with the name of the construct they code for followed by '-T', *e.g.* A-T codes for RNA nanostar A.

From the non-template and template (reverse complement) strands of the blunt end DNA templates, DNA primers for PCR amplification were designed aiming for 40-60% GC content and length between 18 and 26 nt. DNA primers were verified using NUPACK, as well as NEB melting temperature calculator (https://tmcalculator.neb.com/#!/main) for use with Q5 High-Fidelity DNA Polymerase under standard 500 nM primer concentration.

#### 1.2 RNA transcription

##### 1.2.1 Materials

DNA primers were purchased from, and purified by Integrated DNA Technologies (IDT) *via* standard desalting. Unless otherwise specified, double-stranded DNA templates were purchased from Integrated DNA Technologies (IDT) as gBlocks. Only $STV_{apt}$-T was purchased as a double-stranded Ultramer due to its shorter sequence. All DNA strands were received lyophilized and reconstituted at 100 $\mu$M. DNA primers were reconstituted in nuclease-free water (UltraPure DNase/RNase-Free Distilled Water, Invitrogen), while gBlock DNA templates were reconstituted in syringe-filtered TE buffer (10 mM Tris, 1 mM EDTA, pH 8.0), obtained by diluting Tris-EDTA 100× (Sigma-Aldrich) in nuclease-free water. DNA primer concentration was determined by measuring absorbance at 260 nm (average of five repeated measurements) using a Thermo Scientific Nanodrop One Microvolume UV-Vis spectrophotometer using extinction coefficients provided by the supplier. Primers were then diluted to 10 $\mu$M in nuclease-free water as *per* PCR kit instructions. Malachite Green chloride and DFHBI (3,5-difluoro-4-hydroxybenzylidene imidazolinone) were purchased from Sigma-Aldrich. The as-received powders were dissolved in nuclease-free water and DMSO to produce 1 mM and 10 mM stock solutions, respectively. The 10 mM DFHBI solution was then diluted to 1 mM in nuclease-free water. The resulting 1 mM Malachite Green and DFHBI solutions were stored at 4°C and −25°C, respectively.
Recombinant Enhanced Yellow Fluorescent Protein (EYFP) was purchased from RayBiotech, resuspended at 0.67 $mg\,mL^{-1}$ in nuclease-free water as *per* manufacturer's instructions, and stored at -80 °C. EYFP was found to aggregate over time, hence it was used within 2 days from reconstitution. TexasRed Streptavidin conjugate was purchased from Sigma Aldrich (CalBioChem), already resuspended at 1 $mg\,mL^{-1}$ in 50 mM bicarbonate-borate buffer, 0.9% NaCl, 5 $mg\,ml^{-1}$ BSA, pH 8.1, and stored at 4 °C according to specifications. Alexa Fluor 405 Streptavidin conjugate was purchased from Invitrogen, resuspended at 1 $mg\,mL^{-1}$ in Phosphate Buffered Saline (PBS), pH 7.2 (Gibco) with addition of 5 mM sodium azide (0.1 M solution, Sigma Aldrich), as *per* manufacturer's instructions, and stored at -20 °C.

##### 1.2.2 PCR amplification of DNA templates

Amplification of gBlock DNA templates was carried out using Q5 High-Fidelity DNA Polymerase (New England Biolabs). PCR mixtures were prepared on ice following manufacturer instructions, with 10 ng DNA template *per* mixture. PCR mixtures were annealed in a Bio-Rad C1000 Touch Thermal Cycler according to the following protocol from the manufacturer: pre-heating at 98°C; initial denaturation at 98°C for 30 s; 30 amplification cycles (98°C for 10 s, 64-65°C for 20 s depending on primer melting temperature calculated using NEB melting temperature calculator, 72°C for 7 s); final extension at 72°C for 2 min; hold at 4°C. PCR mixtures were stored at 4°C and gel-purified within a week.

##### 1.2.3 Gel electrophoresis and gel image analysis

**Agarose Gel Electrophoresis (AGE).** Agarose gels were prepared at 2% weight/volume agarose (Sigma-Aldrich) in Tris-Borate-EDTA (TBE) 1× buffer (from TBE 10×, Thermo Scientific), with the addition of GelRed nucleic acid gel stain (3×, Biotium). Gels were run at 120 V for 90 min (DNA templates) or 45 min (RNA transcripts).

**Polyacrylamide Gel Electrophoresis (PAGE).** 3.7 mL 30% weight/volume Acrylamide/Bis-Acrylamide Partitioned Solution (29:1, Sigma Aldrich), 4.0 mL 10× TBE (Tris-borate EDTA Buffer, ThermoFisher), and 6.3 g urea (Sigma-Aldrich) were combined to achieve a mixture with 7 M final urea concentration. To obtain a 1.5 mm 8% polyacrylamide gel, this volume was adjusted to 14 mL using UltraPure RNase-free water in a 50 mL Falcon tube. Following the addition of 75 $\mu$L of 10% weight/volume APS (Ammonium Persulfate, Sigma-Aldrich) and 15 $\mu$L TEMED (N, N, N', N'-tetramethylethylenediamine, Sigma-Aldrich), the tube was inverted several times. The final mixture was allowed to polymerise for 20 min in an assembled gel electrophoresis cassette (Bio-Rad). Gels were run at 150 V for 60 min, followed by post-staining with 1× SYBR Gold Nucleic Acid Gel Stain (diluted from 10,000× concentrate in DMSO, ThermoFisher) in 1× TBE buffer for 15 min.

**Imaging.** Imaging was performed using a Syngene G:BOX Chemi XRQ gel documentation system.

**Ladders and loading dyes.** DNA ladders were purchased from Thermo Scientific (molecular biology-grade GeneRuler Ultra Low Range DNA Ladder). Single-stranded RNA ladder (RiboRuler Low Range ssRNA Ladder) and 2× RNA Gel Loading Dye were purchased from ThermoFisher.

**Image analysis.** Lane profiling was performed *via* a semi-automated Python3 script requiring users to specify cropping - in the form of (x, y) coordinates of the upper and lower corners - and number of lanes to profile. Lanes are identified by assuming constant lane-width in the same gel, and profiles are computed as the average intensity across a given lane-width along the height of the gel. The script is publicly available on GitHub https://github.com/GiacomoFabrini/Gel_Image_Analysis_1.0. Images with bright bands on dark background produce easily interpretable lane profiles, where each band corresponds to a peak. For consistent result presentation, images with dark bands on bright background were look-up table inverted prior to analysis to produce similar profiles. In case of clear gel image tilting, images were straightened using rotation-correction in FIJI [9]. Gel images are reported with no contrast enhancement, dark bands on white background for ease of visualisation (obtained, if needed, after look-up table inversion), with overlayed lanes as found by lane profiling.

##### 1.2.4 DNA template purification and quantification

When gel-purifying PCR amplified DNA templates, large combs were adopted to load one PCR reaction (50 $\mu$L + 10 $\mu$L TriTrack DNA Loading Dye 6×, Thermo Scientific) *per* well. Gel bands were cut using a scalpel under the UV illumination of the gel imaging system, using a UV-safe visor. Gel bands were loaded in pairs in 2 mL Eppendorf tubes, treated by adding 4 $\mu$L of Monarch Gel Dissolving Buffer (New England Biolabs) *per* mg of gel, and incubated at 50°C until complete dissolution. The obtained mixtures were purified using the Monarch PCR & DNA Cleanup Kit from New England Biolabs, following manufacturer instructions. Elution was performed with 12-14 $\mu$L *per* gel band pair.
Purified DNA template concentration was determined by measuring absorbance at 260 nm (average of three repeated measurements) using a Nanodrop One with double-stranded DNA settings. Concentrations in ng $\mu$L$^{-1}$ were then converted to micromolar units *via* the Biomath Calculators by Promega, accessible at https://www.promega.co.uk/resources/tools/biomath/.

##### 1.2.5 RNA transcription: kit and protocol

Transcription was carried out using the CellScript T7-FlashScribe Transcription Kit, following manufacturer protocols. The final reaction mixture contained T7 RNA polymerase in the proprietary transcription buffer, complemented by 9 mM of each ribonucleotide triphosphate (rNTP), as well as 0.05 units $\mu$L$^{-1}$ of RNase inhibitor and 10 mM dithiothreitol (DTT). DNA templates were added for a final overall concentration of 40 nM, apart from experiments investigating the effect of template concentration on transcription rate with fluorimetry, for which template concentrations of 10 and 20 nM were also used.
For fluorimetry and imaging, malachite green and DFHBI dyes were added to transcription mixtures in proportions equal to 1 $\mu$L 1 mM dye : 20 $\mu$L mixture, yielding a final concentration of approximately 45.45 $\mu$M *per* dye.

#### 1.3 Sample preparation

##### 1.3.1 Gel electrophoresis on RNA transcripts

RNA transcripts were prepared as described in section 1.2.5 (10 $\mu$L *per* sample). No malachite green or DFHBI dyes were added to the final reaction mixtures. Samples were incubated in a Bio-Rad C1000 Touch Thermal Cycler at 30°C (heated lid at 35°C) for 16-18 h, and then treated with 0.5 $\mu$L of DNase I provided as part of the CellScript T7-FlashScribe Transcription Kit at a concentration of 1 unit $\mu$L$^{-1}$ for 30 minutes at 37°C (heated lid at 40°C). After incubation, samples were diluted 3× with RNase-free water and mixed with 3 $\mu$L of 60% glycerol (used instead of loading dyes). 6 $\mu$L of each diluted sample were loaded into the wells of 2% weight/volume agarose (see section 1.2.3 above). 10 $\mu$L RiboRuler Low Range ssRNA Ladder were loaded in the leftmost lane. For the control DNA nanostars (sequences provided in Table S4), 10 $\mu$L of 0.7 $\mu$M DNA sample (annealed in the same Bio-Rad C1000 Touch

Thermal Cycler) were loaded in the rightmost lane.

**Denaturing PAGE.** RNA transcripts were prepared and treated with DNase I as for native AGE, then diluted 50× with RNase-free water. Sample (4 $\mu$L) and RiboRuler Low Range ssRNA Ladder (2.5 $\mu$L) aliquots were each mixed with equal volumes of 2× RNA Gel Loading Dye, prior to incubation in a Bio-Rad C1000 Touch Thermal Cycler at 70°C (heated lid at 80°C) for 10 min. All samples were promptly transferred on ice before loading onto the gel. 8 $\mu$L of each sample and 5 $\mu$L of RiboRuler Low Range ssRNA Ladder were loaded into the wells of a 7 M urea, 8% polyacrylamide gel.

##### 1.3.2 Imaging - bulk samples in pristine *In Vitro* Transcription Mixture (IVTM)

Bulk samples were prepared as described in section 1.2.5, at 40 nM DNA template concentration, 1× 22 $\mu$L transcription mixture (including malachite green and DFHBI dyes at 45.45 $\mu$M each) *per* sample, unless otherwise specified. See sections 1.4 and 1.5 for further details.

##### 1.3.3 Imaging – effect of buffer exchange on condensate stability.

To inspect the stability of A and B RNA condensates in different ionic conditions, *In Vitro Transcription* reactions were carried out as described in section 1.2.5 (20 $\mu$L *per* sample) but using 384 well plates (black, Greiner Bio-One) rather than glass capillaries to enable facile buffer exchange. DNA template concentration was reduced to 2 nM to account for the different geometry and reduced bottom-surface to volume ratio of the microplate wells compared to the capillaries, thus avoiding the formation of extremely large condensates upon sedimentation. Microplates were sealed with an adhesive aluminium film. Samples were incubated using a custom-made microplate heated stage with temperature set at 30 °C for the plate chamber and at 35 °C for the lid (to prevent evaporation), and then imaged after 24 h to check condensate formation. Samples underwent buffer exchange with either Phosphate-Buffered Saline (PBS) 1× pH 7.4 diluted from (10× PBS, Invitrogen), Tris-EDTA (TE) 1× diluted from (100× TE, ThermoFisher) supplied with 5 mM $MgCl_2$ (Sigma-Aldrich) pH 8.0 or TE 1× supplied with 10 mM $MgCl_2$ pH 8.0 *via* four consecutive washes, separated by 30 min intervals (during which the samples were kept at 30 °C thanks to the heated microscope stage). In particular, for the first wash, 70 $\mu$L of the desired buffer were added to the sample well. For remaining washes, 70 $\mu$L supernatant were removed prior to adding an equal volume of fresh buffer. Samples were imaged after the final wash, as well as after an additional 24 h incubation at 30 °C.

##### 1.3.4 Imaging – effect of crowding agents on bulk assembly.

Samples were prepared as described in section 1.2.5, at 40 nM DNA template concentration, 1× 20 $\mu$L transcription mixture (including malachite green and DFHBI dyes at 45.45 $\mu$M each) *per* sample. To reach the indicated final volume, samples were supplemented with either RNase-free water (for control samples, "No PEG") or poly(ethylene glycol) 200 (PEG 200, Sigma-Aldrich) at a final concentration of 25% volume/volume.

##### 1.3.5 Imaging – FRAP

For FRAP experiments conducted on the Fluorescence Light-Up Aptamers (FLAP) embedded within the condensates, samples were prepared as described in section 1.2.5, at 40 nM DNA template concentration, 1× 20 $\mu$L transcription mixture (including malachite green and DFHBI dyes at 45.45 $\mu$M each) *per* sample. For FRAP conducted using the covalently linked Fluorescein-12-UTPs, samples were prepared as described in section 1.2.5, at 40 nM DNA template concentration, 1× 20 $\mu$L transcription mixture (no Malachite Green and DFHBI dyes were added) *per* sample. 100 mM UTP in the CellScript T7-FlashScribe Transcription Kit was diluted to 50 mM in RNase-free water provided with the kit to obtain 8.95 mM UTPs in each sample. 50 $\mu$M of Fluorescein-12-UTP (Sigma-Aldrich) was added to the final mixtures to label the formed RNA condensates, corresponding to $\sim 0.55\%$ of the total UTP content. All samples were loaded in hollow, rectangular, flat glass capillaries (see section 1.4.1) and incubated in a Bio-Rad C1000 Touch Thermal Cycler at 30°C (heated lid at 35°C) for 24 h before imaging.

##### 1.3.6 Imaging - samples in water-in-oil droplets (synthetic cells)

*In vitro* transcription mixtures were prepared as discussed in section 1.2.5. For protein capturing experiments, unless otherwise specified, the volume of transcription mixtures was increased to 23 $\mu$L to accommodate Enhanced Yellow Fluorescent Protein (EYFP), TexasRed-conjugated Streptavidin (TexasRed-STV) or Alexa405-conjugated Streptavidin (Alexa405-STV), each at a final concentration of 1.25$\mu$M. For control experiments in the absence of target proteins in Fig. S48, transcription mixture volume was kept equal to 22 $\mu$L instead.
In assays relying on protein-binding aptamers ($\mathrm{YFP_{apt}}$, $\mathrm{STV_{apt}}$), the total DNA template concentration was kept equal to 40 nM, and the composition ratio was chosen to be [nanostar DNA template]/[aptamer DNA template] = 3. For TexasRed-STV capture assays *via* $\mathrm{Biotin_{DNA}}$, [$\mathrm{B_{STV}}$-T] was similarly kept at 30 nM, while [$\mathrm{Biotin_{DNA}}$] was chosen to be 10 $\mu$M, yielding an $\approx 8\times$ excess of biotin compared to streptavidin.

**Encapsulation *via* emulsion method.** Encapsulation of *in vitro* transcription mixtures within water-in-oil droplets (synthetic cells) was achieved *via* the emulsion method, similarly to what described in [10]. Briefly, 22-23 $\mu$L of transcription mixture were added on top of 90 $\mu$L of 2% (weight/weight) Pico-Surf (Sphere Fluidics Ltd.), a biocompatible surfactant, in Novec$^{\mathrm{TM}}$ 7500, a fluorinated oil, within an Eppendorf tube. The resulting mixture was vortexed at 2500 rpm for 30 s and then left to equilibrate for 1-2 min before extracting the top layer containing the synthetic cells.

#### 1.4 Imaging

##### 1.4.1 Epifluorescence imaging

Timelapse imaging was performed on a Nikon Eclipse Ti2-E inverted microscope with Perfect Focus System (PFS), equipped with Plan Apochromat $\lambda$ 10× (NA 0.45, WD 4000 $\mu$m) and $\lambda$ 20× (NA 0.75, WD 1000 $\mu$m) objective lenses, a Hamamatsu Orca-Flash 4.0 v3 camera, and a Lumencor SPECTRA X LED engine. The following SPECTRA X LEDs were used to excite the corresponding fluorophores or fluorescent molecule/aptamer complexes: 395 nm for Alexa Fluor 405 Streptavidin conjugate (Alexa405-STV), 470 nm for DFHBI/Broccoli aptamer (BrA), 550 nm for EYFP, 575 nm for TexasRed Streptavidin conjugate (TexasRed-STV), and 640 nm for malachite green (MG)/Malachite Green aptamer (MGA). Final concentration for MG and DFHBI dyes was 45.45 $\mu$M each, with both present in solution for every sample (even samples lacking the corresponding aptamer). MG was omitted only in assays including TexasRed-STV due to fluorescence emission overlap. The overall DNA template concentration was kept constant and equal to 40 nM.
Samples were imaged in hollow, rectangular, flat glass capillaries (either 0.20 mm × 4.00 mm × 50 mm or 0.40 mm × 4.00 mm × 50 mm, VitroCom) sealed and glued on top of glass coverslips (24 mm × 60 mm, Menzel Gläser) *via* a 2-component epoxy (Araldite Rapid). In the case of bulk samples, to avoid the glue coming in contact with the water phase, the sides of the capillary were capped with mineral oil. Glue was allowed to set for 30 min, during which samples were kept in a dark environment at room temperature. The glass coverslips were taped to a peltier-controlled copper temperature stage (Temikra Ltd), and held upside down, allowing only for fluorescence imaging. Temperature was controlled *via* custom XML scripts on a RaspberryPi3 running RaspbianOS.
Imaging was automated *via* ND Acquisition, with PFS enabled to ensure constant z-height during the timelapse. Three non-overlapping fields-of-view (FOVs) *per* capillary (sample) were imaged. Epifluorescence z-stacks (120-160 $\mu$m z-height, distributed from $-20/-60$ to + 100 $\mu$m around the PFS plane, with a 3.5-4 $\mu$m step depending on the run) were captured at each timepoint.
For condensate formation timelapses, both in bulk and in synthetic cells, temperature was set to 30°C and monitored *via* a thermocouple. Samples were imaged every 15 min for 10 h, and every 30 min for further 38 h (with the exception of binary and ternary systems in bulk, where only 42 h overall were collected due to software issues leading to early termination of the run. Data at 48 h were collected manually).
For melting of condensates in bulk, temperature was set to 25°C and increased by 1°C every 15 min up to 75°C. Samples were imaged after 10 min of hold at each temperature.
Due to the time required for sample preparation and setup, imaging started 1-2 h after mixing the DNA templates with the rest of the transcription mixtures (see Tables S5-S6 for a detailed per-figure breakdown). Micrographs and videos have been labelled according to imaging time, with time 0 referring to the start of the imaging run, rather than to the start of transcription. When comparing samples from different runs in the same figures, timepoints have been aligned to reflect any delays in the

run starting times (as in Fig. 3b of the manuscript and Figs S27, S44-S45, S49-S50). Conversely, videos (see below) have all been labelled independently, *i.e.* relatively to the start of their specific run.

##### 1.4.2 Epifluorescence - image extraction and analysis

**Extraction pipeline for micrographs.** Because the file size of each of the time-lapse datasets exceeds the size of the workstation's RAM, we used a custom pipeline written in Python3 to extract the optimal focal planes for each timepoint *per* sample FOV. As metric to define the plane of best focus we opted for Normalised Variance, as described in Refs. [11, 12], which associates optimal focusing with the z-plane displaying the highest ratio between variance and mean intensity along the z-axis.
Briefly, pristine Nikon ND2 files were handled *via* ND2Reader.nd2reader, which creates a pointer (reader) to the file. By accessing the reader metadata, the script looped hierarchically through XY positions (FOVs of samples), channels, timepoints and finally z-levels. The plane of best focus across z-levels for each timepoint in a given FOV and channel was found according to the Normalised Variance metric, and appended to a list. After looping through all timepoints of a given FOV and channel, the sequence of optimal planes was extracted, converted into an array and saved as a TIFF file *via* tiffile.imwrite, choosing the minimum but most efficient compression (zlib) and specifying the axes order as 'TYX' (time, spatial orientation) for later compatibility with FIJI (ImageJ).
When determining planes of best focus for each time-point, three scenarios might occur: condensation occurs in one channel only, in both or in none (as for negative controls). In the case of a single condensating component, the planes of best focus were determined in the associated channel (*via* numpy.argmax), saved and re-used to extract the corresponding planes in the other channel, which were then saved as a separate TIFF file. In the case of two condensing components, or of individual condensates displaying fluorescence in both channels, the normalised variances of separate channels in the same FOV were calculated, summed and the position of the maximum of the resulting profile along the z-axis was found. The timelapses for each channel were then saved sequentially in TIFF format. In the case of no condensating components, planes of best focus were visually inspected and manually extracted by specifying the desired z-range. It is worth noting that, even with samples forming condensates, Normalised Variance can yield incorrect results, usually in very high z planes, in the first few timepoints where no clear condensates can be observed. For this reason, in case the found planes of best focus exceeded the 80% of the overall z-range - indicating the calculated plane was well above the PFS one - and then stabilised to a lower z-level, these initial values were discarded and replaced with the first occurring lower z-level. Epifluorescence micrographs are reported either in their pristine form (no adjustments, composite of all available channels) or after linear rescaling (0.25% pixels (px) saturated, normalisation enabled, using only the channels that show fluorescence) to enhance contrast. All epifluorescence micrographs in the main text (Figures 1-3) are contrast-enhanced, as stated in the respective captions, with pristine counterparts included in Supplementary Figures. Contrast-enhanced

micrographs in Supplementary Figures are labelled with a half-shaded circle for ease of distinction. videos S1-S10 and S12 were generated in FIJI from extracted timelapses in TIFF format *via* the following steps: contrast enhancement (0.25% pixels saturated according to the stack histogram, using only the channels that show fluorescence), calibration *via* px/$\mu$m from ND2 metadata, binning (from 2044 px × 2048 px down to 1022 px × 1024 px, to reduce file size), labelling with a scale bar, time-stamping (or temperature-stamping in the case of the melting of condensates in bulk), and finally export in AVI, each frame compressed in PNG format, at a playback speed of 10 frames *per* second (fps). The resulting AVI files were then processed with a FIJI macro to maintain a constant relative playback speed with respect to experimental time: every frame from 10 h to 48 h was duplicated so as to maintain the same playback speed as the first 10 h. Related videos were collated and labelled in single AVI and converted to MP4 *via* Permute3 for ease of access.

**Extraction pipeline for coarsening analysis.** When extracting data for coarsening analysis by segmentation and extracting the Chord-Length Distribution (see below) as in Figs 1-3, S8, S17, S23 and S29, the pipeline above was modified in the following ways: a) after determining the best focal plane at each timepoint *per* FOV and channel, a z-volume spanning 2 focal planes on either side of it (5 planes or ± 7-8 $\mu$m around the mid plane) was extracted and its maximum intensity z-projection was calculated and saved instead of the single plane described above; b) optimal focal planes for each channel of every FOV were extracted independently, thus discarding the coupled-channels approach described above for both condensating components. Coarsening analysis was performed, in fact, on each channel individually, as differently sized condensates (*e.g.* binary systems with extreme DNA-template ratios) might be located on very different focal planes. In the case of constructs that are fluorescent in both channels, this still yielded the same result as the two fluorescent signals stem from the same condensates and thus the same volume.

**Segmentation and particle analysis.** Segmentation and Particle Analysis for object counting were carried out in FIJI (ImageJ) [9] *via* a custom macro implementing the following pipeline: linear rescaling (0.25% pixels saturated, normalisation enabled), pixel calibration (from ND2 metadata), denoising *via* Gaussian Blur (sigma = 1), two steps of sliding paraboloid background subtraction (smoothing disabled, radius = 1 px), conversion to binary mask *via* automatic thresholding (Li for bulk, Otsu for synthetic cells), and finally particle analysis excluding objects smaller than 10 px$^2$ in area. Segmentation masks and analysis results were saved in TIFF and CSV formats, respectively. The macro operated on split fluorescence channels, thus producing two masks *per* sample FOV. Normalisation was enabled in the first contrast enhancement step to better resolve smaller condensates in the first few timepoints, as it causes each frame to be contrast-enhanced separately without considering the stack histogram. For segmentation, timelapses of maximum intensity z-projected volumes (see above) were used to better capture the size distribution of objects that might be lying on slightly different focal planes. Despite this, different automatic thresholding algorithms had to be used for bulk (Li) and synthetic cells

(Otsu) due to the tendency of Li's algorithm to over-segment (particularly close-by objects in the confined droplet environments) and its sensitivity to incorrectly threshold signal from out-of-focus synthetic cells.

**Chord-Length Distributions from binary masks.** In order to quantify the time-dependent evolution of the typical system length-scales for both droplet-forming A and B and network-forming C designs, we applied Chord Length Distribution (CLD) analysis, previously adopted to characterise phase separation kinetics [13, 14]. CLDs were extracted from segmented binary masks *via* a Python3 script based on PoreSpy, particularly porespy.filters.apply_chords, which in turn relies on scipy and skimage. The selected options allow for adjacent chords (spacing = 1) and include objects touching the edges of the FOV (trim_edges = False). Inputs were binary masks, each representing one channel of a single sample FOV. Channels were only analyzed if condensates were observed in them. For each binary mask, the script looped through timepoints and calculated chords along x and y directions *via* the chord_counts function from PoreSpy. The resulting multidimensional arrays (time, CLD) were arranged in dictionaries (key: sample, subkey1: channel, subkey2: FOV) and were then saved in NPY format *via* numpy. For each FOV and channel of every sample, the resulting chord lengths at every timepoint, both along x and y, were pooled together, converted to $\mu$m *via* the pixel scaling within ND2 metadata, and saved in lists, from which the mean of the CLD, $\mu_{\mathrm{CLD}}$, at each timepoint was computed and saved in new arrays. This resulted in one array of $\mu_{\mathrm{CLD}}$ versus time *per* FOV for each sample-channel combination. We thus obtained as many replicates of the time evolution of the mean as the number of independent FOVs at every timepoint, and proceeded to estimate the mean and the standard deviation among those (proxy for the standard error, as it is the standard deviation of the distribution of mean replicates). CLD data in Figs 1d(i), 2c and 3c are thus represented as mean ± standard deviation among replicates of the mean (*i.e.* across FOVs). Full CLDs are also presented in Figs S17 (single RNA species in bulk), S23 (binary systems in bulk) and S29 (single RNA species in water-in-oil droplets) in ridge plots obtained by vertically stacking kernel density estimation plots (sns.kdeplot) corresponding to representative timepoints. Data on the number of condensates versus time in Fig. 1d(ii) and Fig. S24 is instead represented as the mean number of objects across independent FOVs ± its standard deviation. These data are only presented for A and B condensates and their binary combinations due to the mesh-like nature of C assemblies (which causes Particle analysis to fail).
Note that the mean-chord-length of a circle of radius $r$ is given by $\frac{2}{r}\int_0^r \sqrt{r^2 - x^2}\,dx = \frac{\pi}{2}r$. Hence, for images comprising non-overlapping, monodisperse circular features, one would find $\mu_{\mathrm{CLD}} = \frac{\pi}{2}r$. Therefore, $\mu_{\mathrm{CLD}}$ represents a good proxy for the mean condensate radii for A and B systems at late assembly stages, when condensates appear spherical and have moderate polydispersity. The similarity of these two observables is notable in Fig. 2c and d, when comparing the (plateau values of) $\mu_{\mathrm{CLD}}$ and condensate radii distributions.

**Melting temperature calculation from microscopy thermal ramps.** Resulting timelapses (every frame corresponding to an increase by 1°C starting from 25°C) were segmented as described above for bulk self-assembly, and the binary masks underwent chord-length distribution extraction. Mean (solid line) and standard error on the mean (shaded region) stemming from three non-overlapping FOVs were calculated and plotted for the three constructs A, B and C. The melting temperature of interactions allowing for phase separation into liquid condensates was defined as the temperature value at which the condensate size reached 50% of the original value at 25°C (highlighted by a horizontal dotted line), indicated by a vertical dotted line and by the adjacent text.

**Sintering analysis and inverse capillary velocity computation.** Ten fusion/-coalescence events *per* sample were manually found and cropped from the discussed epifluorescence timelapses, exported as TIFFs and imported in Python3 for the following analysis. For each fusion event, condensates were segmented using Otsu unsupervised thresholding as implemented in skimage.filters.threshold_otsu. The binarised image was then labelled using skimage.measure.label, which associates a different label to each segmented object. As the most frequent label is the background, the fusing droplets were identified by the second and third most frequent labels in the pre-fusion image (two separate objects) or the second most frequent label in the post-fusion image (single object), respectively. Associated geometrical properties, including centroid position, major and minor axis lengths and orientation, were extracted *via* skimage.measure.regionprops_table. First, the pre-fusion characteristic droplet size, $l_c$, was computed as the average of the two droplet radii (in turn determined as half of the average best-fit ellipse axis length) in the timepoint preceding droplet contact. For timepoints following contact, the Aspect Ratio was computed as the ratio between the major and the minor axis length of the best-fit ellipse, and appended to a list. Time constants, $\tau_c$, were obtained by fitting the Aspect Ratio *vs* time profiles with a single exponential decay with the formula $1 + a \cdot e^{\left(-\frac{t}{\tau_c}\right)}$ [15, 16] using scipy.optimize.curve_fit, where $a$ and $\tau_c$ are fitting parameters. Finally, computed time constants were linearly fitted against the pre-fusion characteristic droplet size (Characteristic Length, $l_c$) using scipy.stats.linregress, yielding the inverse capillary velocity as the fit slope. Linear regression parameters (mean ± standard error), $R^2$ and p-value with respect to the null hypothesis (H0) of null slope are as follow: A – slope = 127.40 ± 9.16, intercept: -1108.83 ± 448.69, $R^2$: 0.98, p-value = $6.92 \times 10^{-7}$; B – slope = 152.30 ± 9.06, intercept: -1327.22 ± 438.56, $R^2$: 0.99, p-value = $1.59 \times 10^{-7}$. Micrographs reported in Fig. 1e were contrast-enhanced similarly to what described above, using skimage.exposure.rescale_intensity, with 0.2% of pixels saturated.

**Dilute phase buffering in bulk RNA condensates.** Segmentation masks were obtained in FIJI (Li unsupervised thresholding) and exported as TIFFs as described above. The following analysis was performed in Python3. Time-dependent fluorescence intensity profiles within the condensed and dilute phases were computed by intersecting raw timelapse images with the corresponding binary masks (simply related by inversion, *i.e.* the ∼ operator) using OpenCV (cv2.bitwise_AND). Raw intensity profiles were filtered to lessen the effect of segmentation artifacts by: a) removing dilute phase intensity values above the 90% percentile (stemming from bright halos surrounding segmented condensates, or out-of-focus condensates which segmentation failed to capture), and b) removing condensed phase intensity values below the 10% percentile (resulting from imprecise segmentation borders). The ratio between mean dilute phase intensity and mean condensed phase intensity was computed for each FOV. Data are presented in Fig. S16 as sample mean (solid line) ± standard deviation (shaded region) for both A and B RNA nanostar motifs.

**Circle Hough Transform (CHT) for radial distribution of binary systems.** A semi-automated Python3 script was written to extract the radii of orthogonal condensates in binary systems observed after 48.5 h. Input data were maximum intensity z-projected epifluorescence micrographs (see above) acquired with 20× lens, with three non-overlapping FOVs *per* sample. The script implemented the following pipeline: open z-projected micrograph in TIFF format *via* OpenCV (cv2.imread), convert to 8 bit, denoise *via* Median Blur (cv2.medianBlur, radius = 3 px), grayscale to colour conversion (necessary for following step) and Hough Circle Transform (cv2.HoughCircles, image to accumulator resolution = 5, minDistance = 20 px, minRadius = 20 px, maxRadius = 500 px, method = HOUGH_GRADIENT_ALT). The two threshold parameters were manually tuned. Overlapping circles found by the Hough Circle Transform were removed, leaving only the largest one. After the described quality checks, the following numbers of condensates (condensate A/condensate B) were kept for the respective sample composition: 155/56 (1:4), 78/60 (1:2), 53/66 (1:1), 56/99 (2:1), 50/131 (4:1). Radii in px were converted to $\mu$m *via* px/$\mu$m calibration for 20× lens from ND2 metadata. Data were arranged in two Pandas DataFrame objects, both having sample composition ratio as x, and the radius of condensate A / B as y. Data are presented in Fig. 2d as violin plots produced using seaborn (sns), with condensate A / B radii in red/blue, respectively. As for kernel density estimation for the violin plots, the width of the distributions (along the x axis) was chosen to be proportional to the number of observations (scale = 'count') and the kernel width (bw) was set to 0.45.

##### 1.4.3 Confocal imaging

Apart from Fluorescence Recovery After Photobleaching (FRAP) experiments (see below), Laser Scanning Confocal imaging was performed on a Leica Stellaris 8 (DMi8 CS Premium) inverted microscope. The microscope was equipped with a solid state 405 nm laser as well as a white light laser (WLL, 440-790 nm). The following objective lenses were used: HC PL APO CS2 10× DRY (NA 0.40) and HC PL APO CS2 20× (NA 0.75) DRY. The microscope had four internal detection channels (3 Power HyDS and 1 HyDX) plus a transmitted light detector (Trans PMT) for bright-field imaging. For Alexa Fluor 405 Streptavidin conjugate (Alexa405-STV), the 405 nm solid state laser was used, and emission was recorded around 421 nm. For DFHBI bound to the Broccoli aptamer (BrA), the WLL excitation was tuned to 447 nm and emission was recorded around 501 nm. For EYFP, the WLL excitation was tuned to 514 nm, with emission recorded around 527 nm. For TexasRed Streptavidin conjugate (TexasRed-STV), the WLL excitation was tuned to 595 nm, and emission was recorded around

615 nm. Finally, for malachite green (MG) bound to the Malachite Green aptamer (MGA), the WLL excitation was tuned to 628 nm and emission was recorded around 650-660 nm. A line-sequential illumination mode was adopted, *e.g.* Trans PMT (bright-field), 405 nm (Alexa405-STV) and 628 nm (MG/MGA) in sequence 1, 447 nm (DFHBI/BrA) in sequence 2, 514 nm (EYFP) in sequence 3. Pinhole was set to 1 Airy Unit (AU, automatically calculated). Line-averaging was enabled and set to 2-3. Scan mode along the x direction was selected to be bi-directional after phase calibration. Stage speed was set to 400 Hz for high resolution stills (4096 px × 4096 px or 8192 px × 8192 px) captured with the 10× or 20× lens, and 1000 Hz for zoomed-in z-stacks acquired with the 20× lens. For the latter, the zoom factor was tuned to select a single condensate or droplet, and top/bottom planes were manually tuned, with a Nyquist optimised z-step. Unless otherwise stated, all reported confocal micrographs are pristine. 2D orthogonal cross-sections (XY, XZ, YZ) and volume 3D renderings presented in Figs S38-S40 were generated from z-stacks using 'Sections' in the 3D module of Leica Application Suite (LAS) X. Similarly, clipping of 3D renderings to illustrate the internal microstructure in video S11 were produced *via* the 'Movie Editor' in the 3D module of LAS X, converted to AVI using ffmpeg, collated in FIJI and finally re-exported in MP4 for ease of access using Permute3.

**Image presentation: z-spacing calibration and orthogonal cross-sectional slice extraction for z-stacks.** The z-spacing of all presented confocal lateral projections (Figs S7 and S46) was experimentally calibrated. Calibration was conducted using the uniformly-fluorescent water-in-oil (w/o) droplets (synthetic cells) transcribing non-sticky (*i.e.* soluble) $\overline{\text{A}}$ and $\overline{\text{B}}$ RNA nanostars. Because of the high water/oil interfacial tension, these droplets are expected to be near-perfect spheres, *i.e.* their measured height in z should be equal to the diameter of their equatorial cross-section. The diameter-to-height ratio of these droplets was determined using image segmentation of confocal z-stacks, and used to compute the rescaling factor to calibrate lateral projections. The height of the w/o droplets was determined using the full-width-at-half-maximum of the z-projected fluorescence intensity.
Orthogonal cross-sectional slices in Figs. S7 and S46 were obtained using the FIJI plugin "Ortho-slice Extractor" developed by Stephen Rothery for the Facility of Imaging by Light Microscopy (FILM). More information can be found at the following link: https://www.imperial.ac.uk/media/imperial-college/medicine/facilities/film/macro-description.pdf.

**Image analysis: Scaling of A/B condensate volume with emulsion droplet volume.** Confocal micrographs of pure A and pure B RNA condensates self-assembled within emulsion droplets, acquired with 20× lens, were manually annotated in FIJI, separately labelling the emulsion droplet containers as well as the RNA compartments with circular masks. In particular, 66/70 emulsion droplet-RNA condensate pairs were annotated for A/B, respectively. Area of annotated regions was obtained *via* FIJI measurements, saved in CSV format and analyzed in Python3. Emulsion droplet and condensate volumes were computed from the radii extracted from area values, assuming perfect spherical shape in both cases. The scaling relationship among RNA condensate volume and surrounding emulsion droplet volume was fit to a linear regression model using scipy.stats.linregress (using ordinary least squares) and plotted with seaborn.regplot. Results are presented in Fig. 3d. Linear regression parameters (mean $\pm$ standard error), $R^2$ and p-value with respect to the null hypothesis (H0) of null slope are as follows: A – slope = 0.182 $\pm$ 0.005, intercept: -974.321 $\pm$ 861.614, $R^2$: 0.98, p-value = $2.20 \times 10^{-45}$; B – slope = 0.153 $\pm$ 0.003, intercept: -1462.931 $\pm$ 993.888, $R^2$: 0.99, p-value = $2.72 \times 10^{-59}$.

**Image analysis: Radial size ratio distribution in binary A/B systems in emulsion droplets.** Radial ratios of orthogonal condensates in synthetic cells were calculated by implementing a semi-automated custom pipeline in Python3 similar to the one for bulk condensates. However, as ratios had to be computed for condensates within the same droplet, an ROI (region of interest) selection step was included. Input data were single plane confocal micrographs acquired with 20× lens, one FOV *per* sample. The pipeline executed the following steps: a composite image of bright-field (contrast-enhanced by contrast stretching with skimage.exposure.rescale_intensity, 0.2% of saturated pixels), MG and DFHBI channels was generated and displayed for ROI selection *via* OpenCV (cv2.merge and cv2.selectROIs, respectively), each ROI was cropped from both fluorescent channels, converted to 8 bit (using skimage.util.img_as_ubyte), denoised by Median Blur (cv2.medianBlur, radius = 5 px), converted from grayscale to colour, and finally circles fitting condensates were found *via* Hough Circle Transform (cv2.HoughCircles, image to accumulator resolution = 1, minDistance = 20 px, minRadius = 10 px, maxRadius = 150 px, method = HOUGH_GRADIENT_ALT, parameter1 = 150, parameter2 = 0.6). Only ROIs where at least one circle was found in both channels were kept. All found circles were semi-automatically inspected with a quality control pipeline, with false positives (due to synthetic cells rather than condensates being sometimes detected) removed upon visual inspection. Overlapping circles were removed, with the largest being kept. Finally, only the synthetic cells containing one condensate of each type were considered to compute the radial ratio, as when multiple condensates of a given type are present in the same droplet they tend to be considerably smaller and, as such, inevitably skew the size ratio distribution. After the described quality checks, the following numbers of synthetic cells were kept for the respective sample composition: 33 (1:4), 41 (1:2), 40 (1:1), 31 (2:1), 28 (4:1). Data were arranged in a single Pandas DataFrame, with x being sample composition (single ROI per row), y1 and y2 the radii (px) of condensate A and B, respectively. Radial ratios were computed and are presented in Fig. 3f as violin plots using seaborn (scale = 'count', bw = 0.70). The number of condensates of each type (A, B) *per* synthetic cell with varying DNA template ratio was extracted from the same dataset without the constraint of synthetic cells containing a single condensate of each type, resulting in the following numbers of of examined synthetic cells (composition ratio): 42 (1:4), 42 (1:2), 46 (1:1), 40 (2:1), 32 (4:1)). The fractions of synthetic cells presenting a certain number of A or B-type condensates with respect to the overall number of examined droplets for the same template ratio are presented in two bar charts (A top, B bottom) in Fig. 3g.

**Evaluation of mixing *via* linker nanostars: Segmentation.** The fluorescence channels (MG, DFHBI) of confocal micrographs acquired with the 10× lens (single equatorial planes, already pixel calibrated) were segmented in FIJI with a macro detailing the following pipeline: denoising *via* Gaussian Blur (sigma = 3), sliding paraboloid background subtraction (smoothing disabled, radius = 10 px), and finally conversion to binary mask *via* automatic thresholding with both Li and Otsu algorithms. Segmentation masks were saved separately in TIFF format.

**Evaluation of mixing *via* linker nanostars: Detection and mixing index calculation.** An object detection pipeline similar to the one above was implemented in Python3 to evaluate mixing of orthogonal RNA nanostars upon introduction of a linker RNA construct. We shall define Linker Fraction as the ratio [L-T]/([A-T] + [L-T] + [B-T]). Input data were single plane confocal micrographs acquired with 10× lens, one FOV *per* sample (two in the case of 5:1:5 and 2:1:2 compositions, or equivalently 1/11 and 1/5 Linker Fractions), and the corresponding binary masks from the segmentation step above. The pipeline proceeded as follows. First, a composite image of MG and DFHBI channels was generated and displayed for ROI selection *via* OpenCV (cv2.merge and cv2.selectROIs, respectively). Patches corresponding to each user-selected ROI were cropped from both fluorescent channels as well as from the corresponding binary masks, and converted to 8 bit (using skimage.util.img_as_ubyte). The quantities $I_A^A$, $I_A^B$, $I_B^B$ and $I_B^A$, where $I_X^Y$ is the average intensity of channel X within the binary mask of channel Y, were then computed by bitwise AND (cv2.bitwise_and) followed by extraction of mean value (np.mean). When calculating same-phase terms $I_A^A$ and $I_B^B$, the bitwise AND was computed directly between the patches and the corresponding binary masks, as in $I_A^A$ = mean(bitwiseAND(croppedROI_A,mask_A)). When calculating mixed-terms $I_A^B$ and $I_B^A$ for partially mixed systems (Linker Fraction $< 1/3$, *i.e.* below stoichiometry), the bitwise AND was computed between the patch of one channel and the non-overlapping binary mask from the opposite one (*i.e.* the mask minus the intersection among opposite-type masks), as in $I_A^B$ =mean(bitwiseAND(croppedROI_A,mas_B - (mask_A ∩ mask_B))), to prevent signal from either phase from leaking into the mixed terms. Conversely, in the case of fully mixed systems (Linker Fraction $\geq 1/3$), mixed-terms were computed by bitwise AND among the patches and the union of the masks from both channels, as in $I_A^B$ =mean(bitwiseAND(croppedROI_A,mask_A ∪ mask_B)), as the systems were visibly fully mixed but segmentation can perform differently on channels with different intensities. Finally, the normalised mixed-term ratios $J_A = I_A^B/I_A^A$ and $J_B = I_B^A/I_B^B$, which we refer to as mixing indices, were computed from the quantities described above. The pipeline first calculated the discussed quantities using binary masks obtained *via* Li thresholding, then saved the ROIs for a given sample FOV and repeated the calculation on the same synthetic cells using binary masks according to Otsu thresholding. All synthetic cell ROIs were visually inspected, with those presenting incorrect or extremely small binary masks in either channel being discarded. Bounding boxes of the selected synthetic cells were checked against overlap to ensure independent data points. The resulting intensity ratios for Li and Otsu thresholding were averaged due to the opposite tendencies of the two thresholding algorithms to undersegment (Otsu) and oversegment (Li). Such ratios were arranged in a Pandas DataFrame, with x being the composition ratio and y either of the mixing indices $J_A$ and $J_B$, which can be used as proxy to determine the mixing between the two fluorescent channels, and thus between the *a priori* orthogonal complexes bearing the corresponding fluorogenic aptamers. Checks were implemented to remove NaN values, and data points either below $Q_1 - 1.5\,IQR$ or above $Q_3 + 1.5\,IQR$ were discarded as outliers, where $Q_1$, $Q_3$ and $IQR$ are the first and third quartiles and the interquartile range, respectively. Consequently, the following numbers of synthetic cells *per* sample composition (A-T:L-T:B-T) were kept: 28 (10:1:10), 31 (7:1:7), 30 (5:1:5), 24 (3:1:3), 32 (2:1:2), 38 (1:1:1), 32 (1:2:1). Data are presented in Fig. 4c as violin plots using seaborn (scale = ‘count’, bw = 0.7).

**Evaluation of protein partitioning.** Confocal micrographs acquired with the 20× lens (single equatorial planes) were exported in TIFF format (same micrographs as in Fig. 5b(i)-(iii), bottom). In-focus emulsion droplets were manually annotated in FIJI using the circular selection tool. Binary masks were created from the annotated ROIs using the “Masks from ROIs” FIJI plugin (https://github.com/LauLauThom/MaskFromRois-Fiji) and saved in TIFF format (one mask comprising all annotated droplets). Micrographs and emulsion droplet masks were analysed through a Python3 script performing the following pipeline. First, each mask was thresholded to identify the droplet contours, and masks were generated for the individual droplets. For each droplet within the binary mask, the channel corresponding to the nanostar-binding dye (MG/DFHBI) was denoised *via* Gaussian Blur (cv2.GaussianBlur, sigma = (3, 3)), intersected with the droplet annotation mask, and the RNA organelles were segmented using unsupervised Li thresholding (skimage.filters.threshold_li). This resulted in an inner mask comprising the RNA organelles (above threshold). The outer mask was obtained by inverting the inner mask (*i.e.* selecting below threshold), intersecting the result with the emulsion droplet annotation mask, and applying morphological opening (skimage.morphology.opening) to remove thin bright halos around the previously segmented organelles. Results were manually inspected to remove droplets where organelles were segmented incorrectly, leading to the following number of datapoints being kept *per* condition: 22 (-$\mathrm{YFP_{apt}}$-T), 20 (+$\mathrm{YFP_{apt}}$-T), 11 (-$\mathrm{STV_{apt}}$-T), 28 (+$\mathrm{STV_{apt}}$-T), 32 (-$\mathrm{Biotin_{DNA}}$), 18 (+$\mathrm{Biotin_{DNA}}$).
The channel corresponding to the protein of interest (EYFP/Alexa405-STV/TexasRed-STV) was then intersected with, *i.e.* multiplied by, both inner and outer masks in the previous step, and the mean was computed in each case. This produced the average protein fluorescence intensity within the RNA organelles and outside them, *i.e.* in the droplet lumen. The ratios between inner and outer mean protein fluorescence intensity (one measurement per droplet), termed protein partitioning parameters $\xi$, both in the absence and presence of the corresponding protein-binding construct are presented in the form of one boxplot with superimposed scatter plot (with jitter to minimise overlapping of points) *per* investigated system in Fig. 5c (i)-(iii).

##### 1.4.4 FRAP

**Experiments.** FRAP experiments were performed using a Leica TCS SP5 laser scanning confocal microscope, enclosed in a Ludin environmental chamber with temperature controller. The microscope was equipped with an HCX PL Apo 40× DRY (NA 0.85) objective lens, an Ar-ion multi-line laser (458 nm, 5 mW; 476 nm, 5 mW; 488 nm, 20 mW; 496 nm, 5 mW; 514 nm, 20 mW) and two HeNe single-line lasers (594 nm, 2 mW and 633 nm, 10 mW). The Leica LAS AF FRAP wizard was used for managing FRAP experiments.

For each run, FOVs were imaged pre-bleaching for 2 or 3 frames at 1/6 frames *per* second. Bleaching was induced on multiple point-like ROIs (diffraction-limited spots), positioned on condensates within the FOV. Bleaching of Fluorescence Light-Up Aptamers (FLAPs), was induced using the HeNe 633 nm laser line for A-type condensates (Malachite Green) and Ar-ion laser lines at 476 nm, 488 nm and 496 nm for B-type condensates (DFHBI). For covalently-linked fluorescein, we used Ar-ion laser lines at 488 nm and 496 nm for both A-type and B-type condensates. All laser lines were set to 100% of the output intensity for bleaching. The Ar-ion laser was operated at 68% power. The bleaching time was set to 2 seconds *per* spot for FLAPs and 10 *per* spot seconds for Fluorescein-UTP.

Post-bleaching, samples were imaged at the same framerate for 480-600 seconds (80-100 frames). For both pre- and post-bleaching imaging, excitation was provided using the 488 nm laser line (Broccoli and Fluorescein) or the 633 nm laser line (Malachite Green), operated at 4-6% intensity, depending on the sample.

Throughout the experiments, pinhole size was set to 5 Airy units. For FLAP experiments, data were collected in 3 FOVs for each (A and B) sample, with a total of 25 bleached spots for A and 27 for B. For fluorescein experiments, data were collected in a single FOV per sample, with 7 bleaching spots. Some condensates were left unbleached in each FOV to serve as reference for image analysis.

A standard photomultiplier tube was used as detector, with acquisition window tailored to the emission spectra of the relevant dyes. Image format was 1024 px × 1024 px (387.88 $\mu$m × 387.88 $\mu$m), 8 bit. All experiments were performed at 30°C.

**Image segmentation and data analysis.** Image segmentation and data analysis for FRAP were performed using Matlab tailor-made scripts. Native .lif files were exported to TIFF format using FIJI, and imported in Matlab in raw format.

For FRAP experiments conducted on FLAPs, the centres of the bleached spots were initially identified manually on the first post-bleach frame. The positions of the bleached spots were then refined as follows: 1) images (first post-bleach frames) were cropped using a square ROI centered around the manually-selected bleached spots position, with size smaller than the condensate; 2) a Gaussian filter was applied to the cropped images of the bleached spots to suppress high-spatial-frequency noise; 3) the location of the minimum of the filtered (cropped) image was computed within each circular ROI, and used as a refined location of the bleached spots. Fluorescence intensity $F.I.(t)$ was then sampled over both the pre- and post-bleach phases within circular ROIs (radius 5.7 $\mu$m) centered around the refined positions of the bleached spots. Two or three reference circular ROIs, with radius 18.9 $\mu$m, were selected on non-bleached condensates in each FOV, and used for correcting for photobleaching occurring in the recovery phase. Individual traces collected for each bleached spot were corrected by dividing by the time-dependent average intensity recorded in the reference ROI (both pre- and post-bleach). The corrected intensities $F.I._{\mathrm{C}}(t)$ were then further normalised between 0 and 1 as follows

$$F.I._{\mathrm{N}}(t) = \frac{F.I._{\mathrm{C}}(t) - F.I._{\mathrm{C}}(t=0)}{\langle F.I._{\mathrm{C}} \rangle_{\mathrm{pre-bleach}} - F.I._{\mathrm{C}}(t=0)}, \tag{1}$$

where $t = 0$ marks the first post-bleach frame and $\langle \ldots \rangle_{\mathrm{pre-bleach}}$ indicates the average over the pre-bleach frames. Normalised, corrected curves were then averaged over all bleached spots for each sample, and fitted with an exponential function $f(t) = A \times [1 - \exp(-t/\tau_f)]$, as shown in Fig. S18 (i).

For experiments conducted on fluorescein, circular ROIs with radius of $15.2\,\mu$m were manually selected for each bleached spot on the first post-bleach frame, and used to sample fluorescence intensities pre- and post-bleach. Reference ROIs of the same size were manually selected on non-bleached condensates. For each bleached-spot and reference ROI, we defined a cognate background ROI in a region close to the relevant condensate, but outside of it. The time-dependent intensity of each background ROI was subtracted from that of its cognate bleached-spot/reference ROI. This step was required to remove the strong, position- and time-dependent background signal arising from the fluorescein-UTPs not incorporated in the condensates, which were observed to bleach during the bleaching phase and quickly recover in the initial frames of the post-bleach phase due to free diffusion. As done for experiments conducted on FLAPs, corrected intensity traces of the bleached spots, $F.I._{\mathrm{C}}(t)$, were computed by dividing the (background subtracted) traces of each individual bleached-spot ROI by the average (background subtracted) signal sampled in the reference ROIs. Final normalisation was conducted as

$$F.I._{\mathrm{N}}(t) = \frac{F.I._{\mathrm{C}}(t)}{\langle F.I._{\mathrm{C}} \rangle_{\mathrm{pre-bleach}}}, \tag{2}$$

producing the data shown in Fig. S18 (ii). Note that, despite background subtraction, a small, quick recovery is seen at early times, due to free diffusion of unbound fluorescein-UTPs. Similarly, despite correcting for recovery-phase bleaching, a small downward trend remains.

In all cases, fluorescence intensities were sampled on raw (non-filtered) images.

#### 1.5 Fluorimetry

Fluorescence excitation/emission scans reported in Fig. S1 and fluorescence kinetics assays presented in Figs S13-S15, S31-S33, S40 and S43 were performed using a BMG CLARIOstar Plus plate reader, using transparent UV-Star 384 well-plates (Greiner Bio-One).

**Fluorescence Intensity Kinetics.** For DFHBI bound to the Broccoli aptamer (BrA), excitation and emission were set to $447 \pm 5$ nm and $501 \pm 5$ nm, respectively (dichroic: 474 nm). For malachite green (MG) bound to the Malachite Green aptamer

(MGA), excitation and emission were set to 620 ± 5 nm and 655 ± 5 nm, respectively (dichroic: 637.5 nm). Well multichromatics were enabled to measure both channels in each well for every cycle. Final sample volumes, *i.e.* after adding the DNA templates if necessary, were kept constant across the plates and equal to 22 $\mu$L. Dyes in nuclease-free water ("Water") and dyes in transcription mixtures ("*In Vitro* Transcription Mixture (IVTM)"), where DNA templates were replaced with nuclease-free water, were used as negative controls. In these cases, the nuclease-free water volumes replacing DNA templates were added prior to starting the measurement run. At least 30 min before running an assay, the thermal stage of the instrument was set to 30 °C. "Plate Kinetics" was selected as measurement mode, with the entire plate being scanned at each timepoint. No shaking was selected. Bottom optics readout was chosen due to the plate being covered by an adhesive film to reduce evaporation (see below). Number of flashes was set to 20 *per* well *per* cycle. The instrument was set in "Enhanced Dynamic Range" to avoid signal saturation. Three kinetics windows (*i.e.* measurement intervals) were selected: denser at first (every 60 s for 500 cycles) to better sample the initial transcription burst and assembly transient, and sparser later (every 180 s for 250 cycles, then every 400 s for the remaining 250 cycles). Focal height for each assay was optimised by preparing sacrificial samples 48 h in advance and measuring optimal focal height with the same sample volume in the same plate. For bulk measurements, runs were paused after 11 measurement cycles (around 16 min) to inject the DNA templates and trigger transcription. Injection was performed by manual pipetting and care was taken to ensure proper mixing of the DNA templates within the wells. Immediately after the injection, the plate was covered with an adhesive film (MicroAmp optical adhesive films, Applied Biosystems, ThermoFisher Scientific) to reduce evaporation over the experiment duration (48 h). For measurements in synthetic cells, due to the need to encapsulate the entire transcription mixture, including the DNA template/templates, the plate was sealed prior to the start of the measurement run, with no subsequent injection nor interruption. Similarly to microscopy experiments, final concentration for malachite green and DFHBI dyes was 45.45 $\mu$M each. Unless otherwise specified, the final overall DNA template concentration was 40 nM, *i.e.* when studying binary systems in a 1:1 ratio 20 nM of each DNA template were added. When studying the influence of [DNA template] on transcription rates, different DNA template concentrations (Figs S13-S15 and S31-S33), namely 10, 20 and 40 nM, were reached by reducing the injected (bulk)/added (synthetic cell mimics) volumes while adding the missing amounts in nuclease-free water.

**Excitation/Emission scans.** For Excitation scans, emission was set to 507 ± 8 nm / 650 ± 8 nm and excitation was measured in the range 400-486 nm / 520-630 nm with a 1 nm step for DFHBI+BrA / MG+MGA respectively. For Emission scans, excitation was set to 447 ± 8 nm / 607 ± 8 nm and emission was measured in the range 470-620 nm / 630-720 nm with a 1 nm step for DFHBI+BrA / MG+MGA respectively. All settings from the bulk kinetics assay were re-used, with manual gain adjustment performed on the entire plate before acquisition ("Enhanced Dynamic Range" is unavailable for spectral scans). For RNA nanostar C, only Emission scans were acquired.

**Data Analysis.** Data were acquired in triplicates for all samples, including negative controls. Data processing, analysis and visualisation were carried out in Python3. All sample profiles were first min-max normalised (subtract minimum and divide by (maximum - minimum) range). For each sample/condition, mean and standard deviation were calculated from these normalised triplicates, as well as from raw data (non-normalised). Unless otherwise specified, for each group of samples to be visualised in the same panel/subplot (*e.g.* all Br or all MG channels), ratios between the raw average maximum of each sample and the highest raw average maximum in the corresponding batch were calculated. Finally, normalised profiles (mean, standard deviation) were rescaled by multiplying by the calculated ratios to preserve relative intensities. For binary systems in synthetic cells (Fig. S40), normalised curves were instead rescaled by multiplying them by the ratio between their raw average maximum and the maximum of the 1:1 composition ratio (sticky nanostars, [A-T] = [B-T]) to better highlight the trends in intensity. To evaluate the influence of the DNA template concentration on the initial transcription rate, 30 data points from mean normalised profiles within the first, linear-like transient after injection were linearly fitted using scipy.optimize.curve_fit. The resulting slopes from the fit were plotted versus increasing DNA template concentration in Figs S15 (bulk) and S33 (synthetic cells), with errorbars displaying standard deviations calculated from the covariance matrix of the fitting parameters.

**Estimation of time-dependent RNA nanostar synthesis yield *via* calibration of fluorescence intensity curves.** Micrographs in Fig. 3b show the formation of RNA organelles within droplet-based synthetic cells after just 1 h and 30 min from the start of transcription (*i.e.* 15 min after the start of the imaging run, see Table S6) In order to estimate the time-dependent RNA nanostar synthesis yield, we calibrated the time-resolved fluorescence intensity profiles of B RNA nanostars within synthetic cells with estimates of their concentration obtained from confocal micrographs.

We used the same confocal micrographs employed for the scaling analysis of organelle volume with synthetic cell volume reported in Fig. 3d and outlined in SI Methods section 1.4.3. First, while these micrographs were acquired more than 48 h after the start of transcription, fluorescence intensity profiles only monitor the system for 48 h, thus failing to provide a common reference timeframe. However, fluorimetry measurements for RNA nanostar B plateau around 12 h, well before the end of the investigated time window. Therefore, given that the amount of synthesised RNA nanostars is proportional to the corresponding fluorescence intensity, we can safely map the concentration estimates from confocal micrographs to the endpoint value (48 h) from fluorimetry data.

Specifically, we can roughly gauge the amount of synthesised RNA by modelling each RNA nanostar as a sphere with radius equal to the nanostar arm length $l_{\mathrm{ns}} = n_{\mathrm{bp}}\, d_{\mathrm{bp}}$, where $n_{\mathrm{bp}}$ is the number of base pairs in each arm, and $d_{\mathrm{bp}} = 0.28\,\mathrm{nm}$ [17] is the average distance between consecutive double-stranded RNA base pairs. The number of base pairs in each arm, $n_{\mathrm{bp}}$, was set to 28 bp to take into account the 25 bp-long double-stranded arm and half of the 6 bp-long coaxially stacked KLs upon interaction with neighbouring nanostars. In this simple model, each condensate of volume $V_{\mathrm{C}}$

contains a number of nanostars, $N_{\rm ns}$, equal to:

$$N_{\rm ns} = \frac{V_{\rm C}}{V_{\rm ns}} = \frac{V_{\rm C}}{\frac{4}{3}\,\pi\,l_{\rm ns}^3}. \tag{3}$$

We can thus evaluate the total RNA mass, $m_{\rm ns}$, in such condensate as:

$$m_{\rm ns} = {\rm MW}_{\rm ns}\,N_{\rm ns} \tag{4}$$

where ${\rm MW}_{\rm ns}$ is the molecular weight of a single B nanostar (approximately 83.24 kDa according to the AAT Bioquest RNA molecular weight calculator, found at https://www.aatbio.com/tools/calculate-RNA-molecular-weight-mw). By assuming the fraction of RNA nanostars in the dilute phase to be negligible compared to the one in the condensed phase (as confirmed by Fig. S16, dashed horizontal lines), we can further estimate the overall mass concentration of RNA nanostars within the droplets, $\rho_{\rm ns}$, as:

$$\rho_{\rm ns} = \frac{m_{\rm ns}}{V_{\rm droplet}}. \tag{5}$$

Under the same assumptions, the number concentration of the nanostars in molar units, $n_{\rm ns}$, is:

$$n_{\rm ns} = \frac{\frac{N_{\rm ns}}{N_{\rm A}}}{V_{\rm droplet}}. \tag{6}$$

where $N_{\rm A}$ is the Avogadro constant.
RNA concentration was computed for all B organelle/droplet pairs examined in the above-mentioned scaling analysis. The resulting values (mean ± standard deviation) were used to calibrate the corresponding fluorescence intensity profiles, reported in Fig. S31, by simply scaling the fluorescence intensity value range by the computed endpoint concentration values. The resulting data are presented in Fig. S34.

### 2 Supplementary Tables

**Table S1** Sequences of all RNA nanostars, reported according to 5'-3' convention.

| **Design** | **Sequence** |
|---|---|
| $\overline{\text{A}}$ | GCA CAG UGC UAU GAG UGU GCA CGG GAU CCC GAC UGG CGC<br>CAA CGA ACA AGG UGC CAG GUA ACG AAU GGA UCC UGU GCU<br>GCA CAU UAG AGU CGC UGU AUG ACC CAU AGC ACA AGG GUC GUA<br>CAG CGG CUC UAG UGU GCU CGC GUG CCU CAG AGG ACC UGU CAC<br>CAU GGC GAA AGG UGA UAG GUC CUU UGA GGU ACG CGU CAC<br>UCG UAG CAU UGU GCC UGU CUC CAA CGG CAA AGG AGA UAG |
| A | GCA CAG UGC UAU GAG UGU GCA CGG GAU CCC GAC UGG CCG<br>CAU CGC GAA AGU GGC CAG GUA ACG AAU GGA UCC UGU GCU<br>GCA CAU UAG AGU CGC UGU AUG ACC CAU CGC GAA AGG GUC<br>GUA CAG CGG CUC UAG UGU GCU CGC GUG CCU CAG AGG ACC<br>UGU CAC CAU CGC GAA AGG UGA UAG GUC CUU UGA GGU ACG<br>CGU CAC UCG UAG CAU UGU GCC UGU CUC CAU CGC GAA AGG<br>AGA UAG |
| $\overline{\text{B}}$ | GCA CAG UGC UAU GAG UGU CGC GAC GGA GAC GGU CGG GUC<br>CAG AUA GGC CAU GGC GAA AGG UCU AUC UGU CGA GUA GAG<br>UGU GGG CUC CGU CGC GUG CAC AUU AGA GUC GCU GUA UGA<br>CCC AUA GCA CAA GGG UCG UAC AGC GGC UCU AGU GUG CUG<br>CAC AGU GUC UGU GCG ACU GCA CCC AAC GAA CAA GGG UGU<br>AGU CGC AUA GAC AUU GUG CUC ACU CGU AGC AUU GUG CCU<br>GUC UCC AAC GGC AAA GGA GAU AG |
| B | GCA CAG UGC UAU GAG UGU CGC GAC GGA GAC GGU CGG GUC<br>CAG AUA GGC CAG UCG ACA AGG UCU AUC UGU CGA GUA GAG<br>UGU GGG CUC CGU CGC GUG CAC AUU AGA GUC GCU GUA UGC<br>CAC AGU CGA CAA GUG GCG UAC AGC GGC UCU AGU GUG CUG<br>CAC AGU GUC UGU GCG ACU GCA CCC AGU CGA CAA GGG UGU<br>AGU CGC AUA GAC AUU GUG CUC ACU CGU AGC AUU GUG CCU<br>GUC UCC AGU CGA CAA GGA GAU AG |
| C | GCA CAG UGC UAU GAG UGU CGC GAC GGA GAC GGU CGG GUC<br>CAG AUA GGC CAG GUA CCA AGG UCU AUC UGU CGA GUA GAG<br>UGU GGG CUC CGU CGC GUG CAC AUU AGA GUC GCU GUA UGC<br>CAC AGG UAC CAA GUG GCG UAC AGC GGC UCU AGU GUG CUG<br>CAC GGG AUC CCG ACU GGC CGC AGG UAC CAA GUG GCC AGG<br>UAA CGA AUG GAU CCU GUG CUC ACU CGU AGC AUU GUG CCU<br>GUC UCC AGG UAC CAA GGA GAU AG |
| | **Continues on next page** |

Table S1 – continued from previous page.

| Design | Sequence |
|---|---|
| $\tilde{C}$ | GCA CAG UGC UAU GAG UGU GCA CAG UGU CUG UGC GAC UGC ACG CAG GUA CCA AGC GUG UAG UCG CAU AGA CAU UGU GCU GCA CAU UAG AGU CGC UGU AUG ACC CAG GUA CCA AGG GUC GUA CAG CGG CUC UAG UGU GCU CGC GUG CCU CAG AGG ACC UGU CAG CAG GUA CCA AGC UGA UAG GUC CUU UGA GGU ACG CGU CAC UCG UAG CAU UGU GCC UGU CUC CAG GUA CCA AGG AGA UAG |
| L | GCA CAG UGC UAU GAG UGU GCA CAG UGU CUG UGC GAC UGC ACC CAU CGC GAA AGG GUG UAG UCG CAU AGA CAU UGU GCU GCA CAU UAG AGU CGC UGU AUG AGG GAU CGC GAA ACC CUC GUA CAG CGG CUC UAG UGU GCU CGC GUG CCU CAG AGG ACC UGU CAC CAG UCG ACA AGG UGA UAG GUC CUU UGA GGU ACG CGU CAC UCG UAG CAU UGU GCC UGU CUC CAG UCG ACA AGG AGA UAG |
| $A_{YFP}$ | GGA UCC GGU UGC AAA GCA CAG UGC UAU GAG UGU GCA CGG GAU CCC GAC UGG CCG CAU CGC GAA AGU GGC CAG GUA ACG AAU GGA UCC UGU GCU GCA CAU UAG AGU CGC UGU AUG ACC CAU CGC GAA AGG GUC GUA CAG CGG CUC UAG UGU GCU CGC GUG CCU CAG AGG ACC UGU CAC CAU CGC GAA AGG UGA UAG GUC CUU UGA GGU ACG CGU CAC UCG UAG CAU UGU GCC UGU CUC CAU CGC GAA AGG AGA UAG |
| $YFP_{apt}$ | GGC AAC CGG AUC CAA AGC AGC UUC UGG ACU GCG AUG GGA GCA CGA AAC GUC GUG GCG CAA UUG GGU GGG GAA AGU CCU UAA AAG AGG GCC ACC ACA GAA GCU GCA |
| $B_{STV}$ | GCA CUA GAC CAC AAA GCA CAG UGC UAU GAG UGU CGC GAC GGA GAC GGU CGG GUC CAG AUA GGC CAG UCG ACA AGG UCU AUC UGU CGA GUA GAG UGU GGG CUC CGU CGC GUG CAC AUU AGA GUC GCU GUA UGC CAC AGU CGA CAA GUG GCG UAC AGC GGC UCU AGU GUG CUG CAC AGU GUC UGU GCG ACU GCA CCC AGU CGA CAA GGG UGU AGU CGC AUA GAC AUU GUG CUC ACU CGU AGC AUU GUG CCU GUC UCC AGU CGA CAA GGA GAU AG |
| $STV_{apt}$ | GUG GUC UAG UGC UUU AUG CGG CCG CCG ACC AGA AUC AUG CAA GUG CGU AAG AUA GUC GCG GGU CGG CGG CCG CAU |

**Table S2** Sequences of all nontemplate strands of dsDNA templates coding for RNA nanostars, reported according to 5'-3' convention.

| Design | Sequence |
|---|---|
| $\overline{\text{A}}$-T | TTC TAA TAC GAC TCA CTA TAG CAC AGT GCT ATG AGT GTG CAC GGG ATC CCG ACT GGC GCC AAC GAA CAA GGT GCC AGG TAA CGA ATG GAT CCT GTG CTG CAC ATT AGA GTC GCT GTA TGA CCC ATA GCA CAA GGG TCG TAC AGC GGC TCT AGT GTG CTC GCG TGC CTC AGA GGA CCT GTC ACC ATG GCG AAA GGT GAT AGG TCC TTT GAG GTA CGC GTC ACT CGT AGC ATT GTG CCT GTC TCC AAC GGC AAA GGA GAT AG |
| A-T | TTC TAA TAC GAC TCA CTA TAG CAC AGT GCT ATG AGT GTG CAC GGG ATC CCG ACT GGC CGC ATC GCG AAA GTG GCC AGG TAA CGA ATG GAT CCT GTG CTG CAC ATT AGA GTC GCT GTA TGA CCC ATC GCG AAA GGG TCG TAC AGC GGC TCT AGT GTG CTC GCG TGC CTC AGA GGA CCT GTC ACC ATC GCG AAA GGT GAT AGG TCC TTT GAG GTA CGC GTC ACT CGT AGC ATT GTG CCT GTC TCC ATC GCG AAA GGA GAT AG |
| $\overline{\text{B}}$-T | TTC TAA TAC GAC TCA CTA TAG CAC AGT GCT ATG AGT GTC GCG ACG GAG ACG GTC GGG TCC AGA TAG GCC ATG GCG AAA GGT CTA TCT GTC GAG TAG AGT GTG GGC TCC GTC GCG TGC ACA TTA GAG TCG CTG TAT GAC CCA TAG CAC AAG GGT CGT ACA GCG GCT CTA GTG TGC TGC ACA GTG TCT GTG CGA CTG CAC CCA ACG AAC AAG GGT GTA GTC GCA TAG ACA TTG TGC TCA CTC GTA GCA TTG TGC CTG TCT CCA ACG GCA AAG GAG ATA G |
| B-T | TTC TAA TAC GAC TCA CTA TAG CAC AGT GCT ATG AGT GTC GCG ACG GAG ACG GTC GGG TCC AGA TAG GCC AGT CGA CAA GGT CTA TCT GTC GAG TAG AGT GTG GGC TCC GTC GCG TGC ACA TTA GAG TCG CTG TAT GCC ACA GTC GAC AAG TGG CGT ACA GCG GCT CTA GTG TGC TGC ACA GTG TCT GTG CGA CTG CAC CCA GTC GAC AAG GGT GTA GTC GCA TAG ACA TTG TGC TCA CTC GTA GCA TTG TGC CTG TCT CCA GTC GAC AAG GAG ATA G |
| C-T | TTC TAA TAC GAC TCA CTA TAG CAC AGT GCT ATG AGT GTC GCG ACG GAG ACG GTC GGG TCC AGA TAG GCC AGG TAC CAA GGT CTA TCT GTC GAG TAG AGT GTG GGC TCC GTC GCG TGC ACA TTA GAG TCG CTG TAT GCC ACA GGT ACC AAG TGG CGT ACA GCG GCT CTA GTG TGC TGC ACG GGA TCC CGA CTG GCC GCA GGT ACC AAG TGG CCA GGT AAC GAA TGG ATC CTG TGC TCA CTC GTA GCA TTG TGC CTG TCT CCA GGT ACC AAG GAG ATA G |

**Continues on next page**

Table S2 – continued from previous page.

| Design | Sequence |
|---|---|
| $\tilde{C}$-T | TTC TAA TAC GAC TCA CTA TAG CAC AGT GCT ATG AGT GTG CAC AGT GTC TGT GCG ACT GCA CGC AGG TAC CAA GCG TGT AGT CGC ATA GAC ATT GTG CTG CAC ATT AGA GTC GCT GTA TGA CCC AGG TAC CAA GGG TCG TAC AGC GGC TCT AGT GTG CTC GCG TGC CTC AGA GGA CCT GTC AGC AGG TAC CAA GCT GAT AGG TCC TTT GAG GTA CGC GTC ACT CGT AGC ATT GTG CCT GTC TCC AGG TAC CAA GGA GAT AG |
| L-T | TTC TAA TAC GAC TCA CTA TAG CAC AGT GCT ATG AGT GTG CAC AGT GTC TGT GCG ACT GCA CCC ATC GCG AAA GGG TGT AGT CGC ATA GAC ATT GTG CTG CAC ATT AGA GTC GCT GTA TGA GGG ATC GCG AAA CCC TCG TAC AGC GGC TCT AGT GTG CTC GCG TGC CTC AGA GGA CCT GTC ACC AGT CGA CAA GGT GAT AGG TCC TTT GAG GTA CGC GTC ACT CGT AGC ATT GTG CCT GTC TCC AGT CGA CAA GGA GAT AG |
| $A_{YFP}$-T | TTC TAA TAC GAC TCA CTA TAG GAT CCG GTT GCA AAG CAC AGT GCT ATG AGT GTG CAC GGG ATC CCG ACT GGC CGC ATC GCG AAA GTG GCC AGG TAA CGA ATG GAT CCT GTG CTG CAC ATT AGA GTC GCT GTA TGA CCC ATC GCG AAA GGG TCG TAC AGC GGC TCT AGT GTG CTC GCG TGC CTC AGA GGA CCT GTC ACC ATC GCG AAA GGT GAT AGG TCC TTT GAG GTA CGC GTC ACT CGT AGC ATT GTG CCT GTC TCC ATC GCG AAA GGA GAT AG |
| $YFP_{apt}$-T | TTC TAA TAC GAC TCA CTA TAG GCA ACC GGA TCC AAA GCA GCT TCT GGA CTG CGA TGG GAG CAC GAA ACG TCG TGG CGC AAT TGG GTG GGG AAA GTC CTT AAA AGA GGG CCA CCA CAG AAG CTG CA |
| $B_{STV}$-T | TTC TAA TAC GAC TCA CTA TAG CAC TAG ACC ACA AAG CAC AGT GCT ATG AGT GTC GCG ACG GAG ACG GTC GGG TCC AGA TAG GCC AGT CGA CAA GGT CTA TCT GTC GAG TAG AGT GTG GGC TCC GTC GCG TGC ACA TTA GAG TCG CTG TAT GCC ACA GTC GAC AAG TGG CGT ACA GCG GCT CTA GTG TGC TGC ACA GTG TCT GTG CGA CTG CAC CCA GTC GAC AAG GGT GTA GTC GCA TAG ACA TTG TGC TCA CTC GTA GCA TTG TGC CTG TCT CCA GTC GAC AAG GAG ATA G |
| $STV_{apt}$-T | TTC TAA TAC GAC TCA CTA TAG TGG TCT AGT GCT TTA TGC GGC CGC CGA CCA GAA TCA TGC AAG TGC GTA AGA TAG TCG CGG GTC GGC GGC CGC AT |

**Table S3** PCR primers, their sequences reported according to 5'-3' convention, as well as the DNA templates they amplify. The sequences of the corresponding DNA templates can be found in Table S2.

| **Strand** | **Templates** | **Sequence** |
|---|---|---|
| Primer f (forward) | $\overline{\text{A}}$-T, A-T, $\overline{\text{B}}$-T, B-T, C-T, $\tilde{\text{C}}$-T, L-T | TTC TAA TAC GAC TCA CTA TAG CAC AG |
| Primer r$\alpha$ (reverse) | $\overline{\text{A}}$-T, $\overline{\text{B}}$-T | CTA TCT CCT TTG CCG TTG GA |
| Primer r$\beta$ (reverse) | A-T | CTA TCT CCT TTC GCG ATG GA |
| Primer r$\gamma$ (reverse) | B-T, L-T | CTA TCT CCT TGT CGA CTG GAG |
| Primer r$\delta$ (reverse) | C-T, $\tilde{\text{C}}$-T | CTA TCT CCT TGG TAC CTG GAG |
| Primer f2 (forward) | $\text{A}_{\text{YFP}}$-T | TTC TAA TAC GAC TCA CTA TAG GAT CCG |
| Primer r2 (reverse) | $\text{A}_{\text{YFP}}$-T | CTA TCT CCT TTC GCG ATG GA |
| Primer f3 (forward) | $\text{B}_{\text{STV}}$-T | TTC TAA TAC GAC TCA CTA TAG CAC TAG A |
| Primer r3 (reverse) | $\text{B}_{\text{STV}}$-T | CTA TCT CCT TGT CGA CTG GAG A |
| Primer f4 (forward) | $\text{YFP}_{\text{apt}}$-T | TTC TAA TAC GAC TCA CTA TAG GCA |
| Primer r4 (reverse) | $\text{YFP}_{\text{apt}}$-T | TGC AGC TTC TGT GGT GG |
| $\text{Biotin}_{\text{DNA}}$ | not applicable | GTG GTC TAG TGC TTT /3Bio/ |

**Table S4** Sequences of oligonucleotide strand components for DNA nanostars used as controls in native agarose gels on RNA transcripts (Fig. S4). Sequences are reported according to 5'-3' convention.

| **Strand** | **Sequence** |
|---|---|
| Core 1 | GAT CGC CGC CGC AAT CAC GCG CGT GCT CGG CGC CAG CAG TCC TGG CG |
| Core 2 | GAT CGC CGC CAG GAC TGC TGG CGC CGT CGC TTC TCT TCA TAA CAA CG |
| Core 3 | GAT CGC CGT TGT TAT GAA GAG AAG CGT CGC TCT GGC ACA GGT GTA CG |
| Core 4 | GAT CGC CGT ACA CCT GTG CCA GAG CGT GCA CGC GCG TGA TTG CGG CG |

**Table S5** Time delay ([hh:mm] for hours:minutes) between start of transcription (*i.e.* sample mixing) and start of timelapse run for imaging in bulk.

| Samples | Figures | Time delay [hh:mm] |
|---|---|---|
| A, Ā, B, B̄, C | 1b, S5, S10 | 01:25 |
| Binary sticky mixtures with varying [A-T]/[B-T] | 2a, S19 | 01:24 |
| Binary non-sticky mixtures with [A-T]/[B-T] = 1 | 2b, S21 | 00:55 |
| Ternary sticky mixtures | S25 | 01:24 |

**Table S6** Time delay ([hh:mm] for hours:minutes) between start of transcription (*i.e.* sample mixing) and start of timelapse run for imaging within droplet-based synthetic cells.

| Samples | Figures | Time delay [hh:mm] |
|---|---|---|
| A, B | 3b, S27 | 01:17 |
| C | 3b, S27 | 01:00 |
| Ā, B̄ | S30 | 01:00 |
| Binary sticky mixtures with [A-T]/[B-T] = 1 | S35, S36 | 01:15 |
| Binary non-sticky mixtures with [A-T]/[B-T] = 1 | S38 | 01:17 |
| Ternary sticky mixtures | S41 | 01:00 |
| Linker-mixed systems with Linker Fraction $\neq \frac{1}{7}, \frac{1}{15}$ | S44, S45 | 01:30 |
| Linker-mixed systems with Linker Fraction $= \frac{1}{7}, \frac{1}{15}$ | S44, S45 | 01:59 |
| $A_{YFP}$ capturing EYFP *via* $YFP_{apt}$ | S49, S50 | 01:21 |
| $B_{STV}$ capturing Alexa405-STV *via* $STV_{apt}$ | S49, S50 | 01:02 |
| $B_{STV}$ capturing TexasRed-STV *via* $Biotin_{DNA}$ | S49, S50 | 01:04 |

### 3 Supplementary Figures

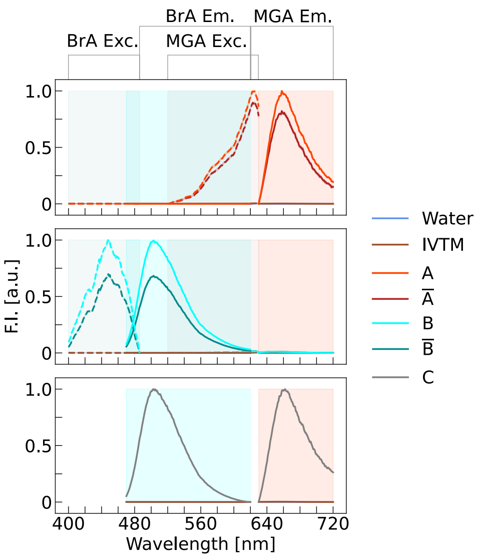


**Fig. S1 Fluorescence excitation and emission scans for RNA nanostars in bulk.** Emission (Em., solid lines) scans are presented for all motifs, while excitation (Exc., dashed lines) scans are included only for negative controls (dyes in water and transcription mixtures) and constructs bearing a single aptamer (top and middle panels). Exc./Em. Scans were performed both in the DFHBI + Broccoli aptamer (BrA, cyan) and in the malachite green (MG) + MG aptamer (MGA, red) bands (see Methods), as indicated by color-coding and text label. (Top) Constructs A and $\overline{A}$, hosting MGA, display the characteristic excitation/emission signature of MG bound to MGA, with peak excitation/emission around 616-630/650-660 nm, respectively, and no significant excitation or emission in the DFHBI+BrA band. (Middle) Constructs B and $\overline{B}$, hosting BrA, display instead the characteristic excitation/emission signature of DFHBI bound to BrA, with peak excitation/emission around 447/500-510 nm, respectively, with no significant excitation or emission in the MG+MGA band. (Bottom) Construct C features both BrA and MGA aptamers, and, as such, displays both emission signatures in the corresponding bands. Data are shown as normalised (see Methods) mean (solid or dashed line) ± standard deviation (shaded region) from triplicates.

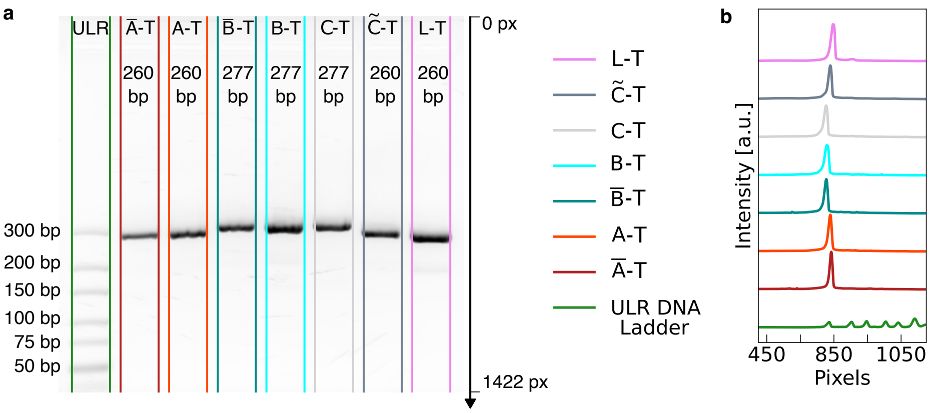


**Fig. S2 Size of PCR amplified DNA templates investigated *via* agarose gel electrophoresis.** (a) Agarose gel (see SI Methods, section 1.2.3) depicting the electrophoretic mobilities of PCR amplified DNA templates $\overline{A}$-T, A-T, $\overline{B}$-T, B-T, C-T, $\tilde{C}$-T (same Kissing Loop domain as construct C but with no fluorogenic aptamers embedded) and L-T. Ultra-Low-Range (ULR) DNA ladder is used as reference (green). Length in base pairs for ladder markings is reported as text close to the corresponding bands. One 50 $\mu$L PCR (10 ng gBlock *per* PCR) was used *per* lane. For the ULR ladder, 1.5 $\mu$g DNA were loaded. (b) Lane intensity profiles extracted from gel in (a) *via* gel lane profiling script (see SI Methods, section 1.2.3). Visual inspection of the gel in (a) and the lane profiling in (b) confirms the expected sizes of the DNA templates coding for the designed constructs (expected length in base pairs (bp) is reported on top). While all PCR amplified DNA templates show a single main band, some display shorter and faster migrating secondary bands (see L-T). The bands corresponding to the expected sizes were extracted and eluted to obtain the purified products for *in vitro* transcription (see SI Methods, section 1.2.4).

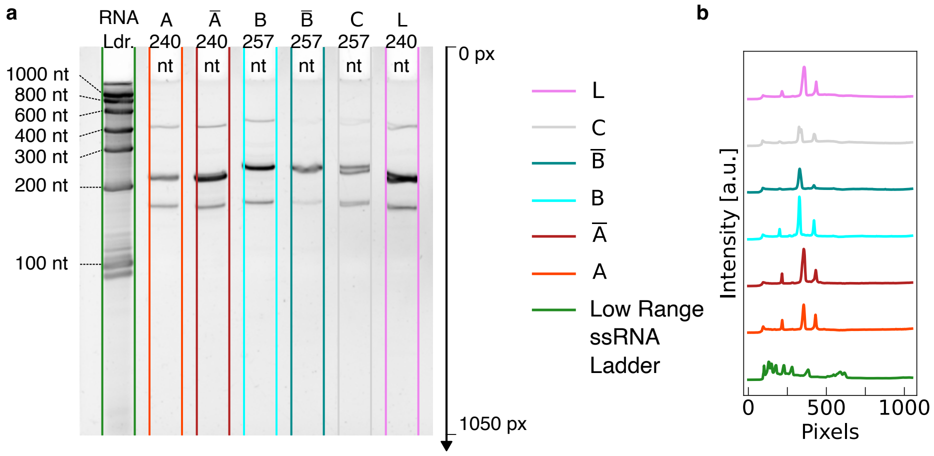


**Fig. S3 Denaturing polyacrylamide gel investigating the electrophoretic mobilities of the transcribed RNA nanostars.** (a) Denaturing polyacrylamide (8%, 7 M urea, see SI Methods, sections 1.2.3 and 1.3.1) depicting the electrophoretic mobilities of transcribed RNA nanostars A, $\overline{A}$, B, $\overline{B}$, C, and L. RiboRuler Low Range single-stranded RNA Ladder ("RNA Ldr") was used as a reference (green), with nucleotide (nt) lengths for markings shown on the left. (b) Lane intensity profiles extracted from gel in (a) *via* gel lane profiling script (see SI Methods, section 1.2.3). Visual inspection of the gel in (a) and the lane profiling in (b) confirms the expected sizes of the transcribed RNA constructs (compare the main band of each sample/lane with the expected length in nt reported on top), and the presence of small populations of truncated and extended products. Truncated products may occur due to early transcription termination caused by the strong secondary structure or specific sequence of one of the RNA nanostar arms [18]. Given the size of the truncated products, termination is likely to occur when transcribing the fourth RNA arm in order of transcription ("opposite" to the aptamer-bearing arms). Elongated products may be due to self-templated addition [19]. The slight band splitting in $\overline{B}$, C and L lanes is likely an artefact of gel loading.

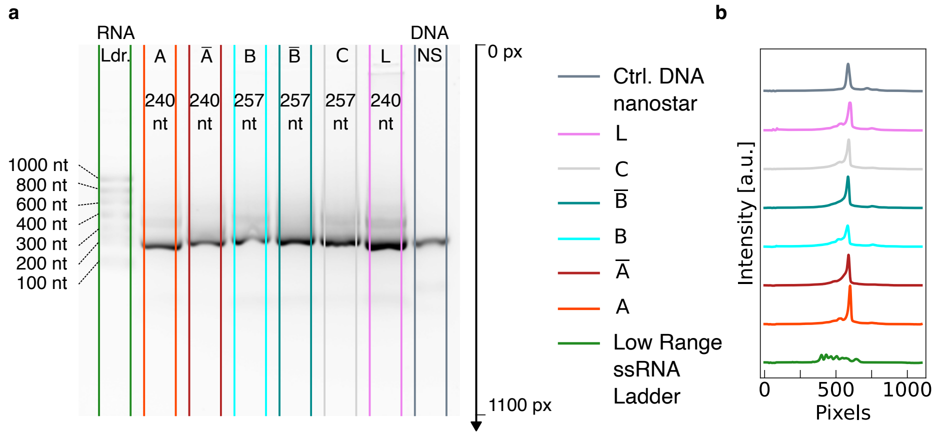


**Fig. S4 Native agarose gel investigating the electrophoretic mobilities of the transcribed RNA nanostars.** (a) Agarose gel (see SI Methods, sections 1.2.3 and 1.3.1) of RNA transcripts A, $\overline{\text{A}}$, B, $\overline{\text{B}}$, C, L, and a control DNA nanostar (NS). The latter has similar size (20 bp dsDNA arms, approximately equal in length to 25 bp dsRNA) and topology (four arms) to the RNA nanostars. The DNA oligonucleotide sequences used for the nanostars are reported in Table S4. RiboRuler Low Range single-stranded RNA Ladder was used as a reference, with nucleotide (nt) lengths for markings reported on the left. (b) Lane intensity profiles extracted from the gel in (a) *via* gel lane profiling script (see SI Methods, section 1.2.3). Visual inspection of the gel in (a) and the lane profiling in (b) confirms that RNA nanostars co-transcriptionally self-assemble into the intended nanostar topology, exhibiting migration behaviour in excellent agreement with that of annealed DNA nanostars with similar size and topology.

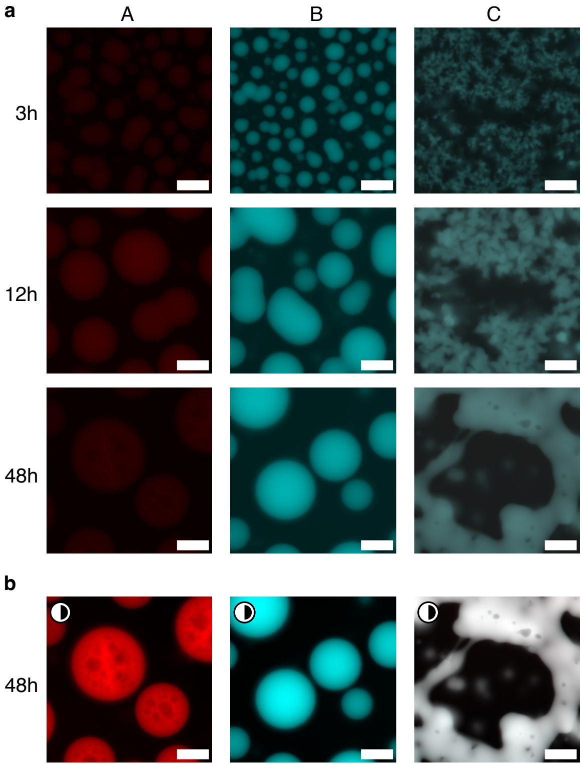


**Fig. S5 Epifluorescence micrographs of RNA condensates in bulk.** (a) Pristine (*i.e.* non contrast-enhanced) epifluorescence micrographs of representative timepoints (3, 12, 48 h, along vertical axis) depicting the bulk self-assembly of condensates from constructs A, B and C, respectively. Micrographs for timepoints at 3 h and 12 h correspond to those in Fig. 1b. (b) Contrast-enhanced (as highlighted by the half-shaded circle) epifluorescence micrograph at 48 h corresponding to the ones in (a) (bottom row). All timestamps are reported with respect to the start of timelapse imaging (see SI Methods, section 1.4.1 and Table S5). Scale bars are 50 $\mu$m. Representative contrast-enhanced videos are provided in video S1 (top).

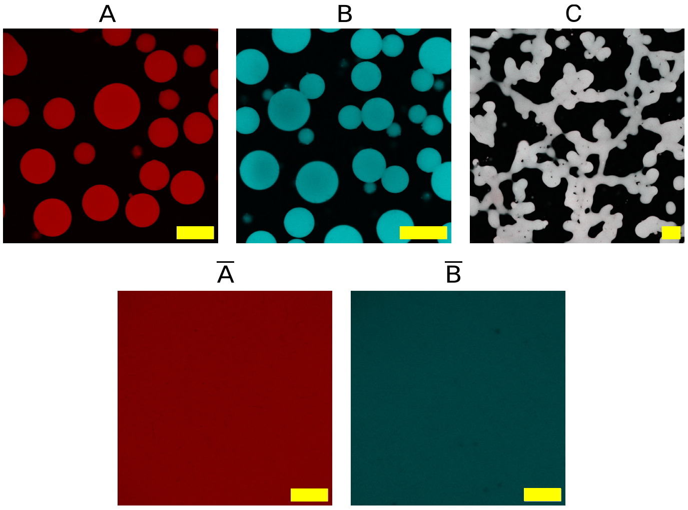


**Fig. S6 Confocal micrographs depicting condensate formation, or lack thereof, in samples of sticky and non-sticky RNA nanostars.** Samples were imaged after more than 48 h. Micrographs are pristine composites of MG and DFHBI channels, and have been binned down to the desired size (1024 px × 1024 px). Scale bars are 100 $\mu$m.

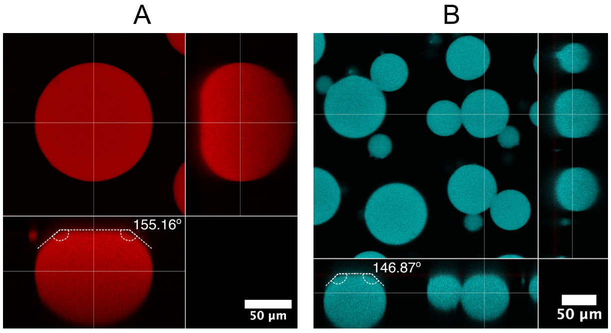


**Fig. S7 Orthogonal confocal cross-sections depicting the shape of A and B RNA condensates transcribed in bulk.** Samples were imaged after more than 48 h. Micrographs are pristine composites of MG and DFHBI channels. Larger RNA condensates deviate from the ideal spherical shape and appear oblate, with a flat contact region with the glass substrate. Z-spacing calibration and procedure to obtain orthogonal cross-sections are described in SI Methods, section 1.4.3. Contact angles, as measured *via* manual annotation using the Angle tool in FIJI [9], are reported for representative condensates (A: 155.16°, B: 146.87°). Scale bars are 50 $\mu$m.

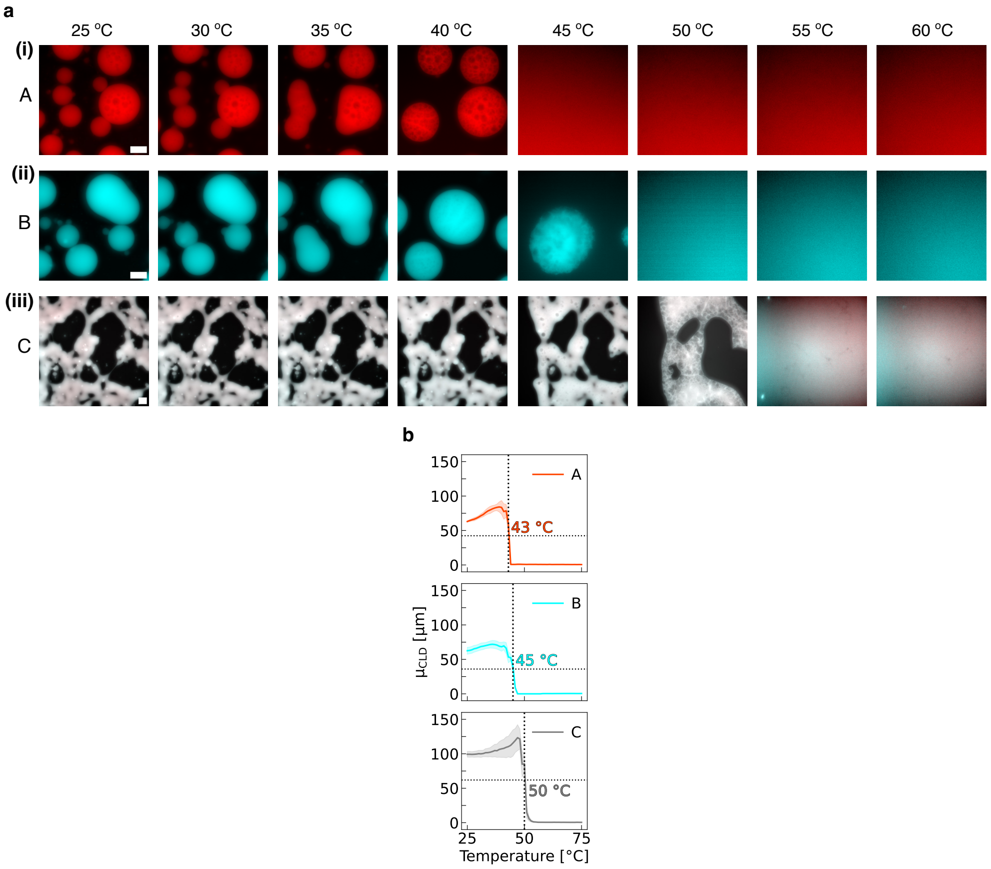


**Fig. S8 Melting of RNA condensates in bulk.** (a) Contrast-enhanced epifluorescence micrographs depicting representative temperatures during melting of A (i), B (ii) and C (iii) condensates, respectively. RNA condensates were allowed to self-assemble at 30 °C for 48 h, stored for 12 h at room temperature in a dark environment. Finally, they were imaged while slowly increasing the temperature from 25 °C up to 75 °C at +1 °C every 15 min. Condensates first appear to swell, as indicated by the formation of holes/bubbles, and then disassemble. All micrographs in each series show the same FOV area, thus sharing the same scale bar. Scale bars are 50 $\mu$m. (b) Change in condensate size with increasing temperature examined *via* the mean of the chord-length distribution ($\mu_{\mathrm{CLD}}$, see SI Methods, section 1.4.2 Melting temperatures ($T_{\mathrm{M}}$) of KL interactions (vertical dotted lines and adjacent text), defined as the temperature at which condensates reach 50% of their $\mu_{\mathrm{CLD}}$ at 25 °C (horizontal dotted line), support trends in interaction strength hypothesized from condensate morphologies at 30 °C. Data are presented as mean (solid line) ± standard deviation among mean replicates (shaded region) from three non-overlapping fields-of-view within the same sample. Representative contrast-enhanced videos are provided in video S2.

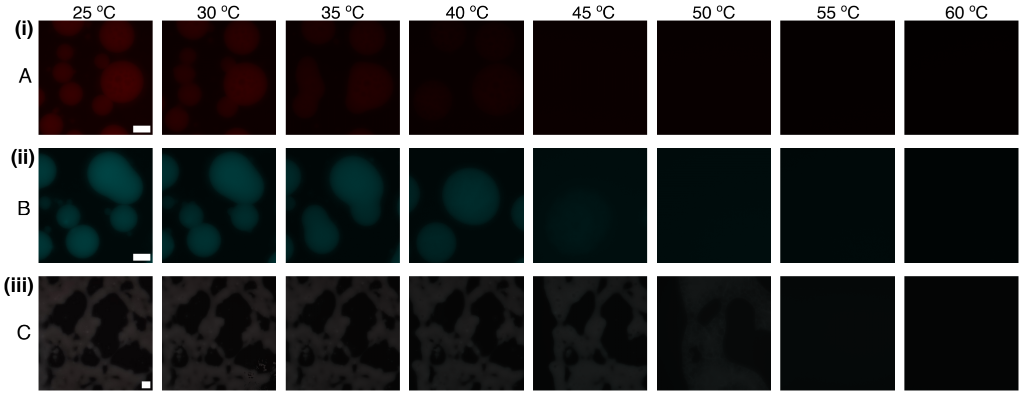


**Fig. S9 Pristine epifluorescence micrographs depicting melting of RNA condensates in bulk.** These micrographs correspond to contrast-enhanced ones in Fig. S8. All micrographs in each series (i, ii, iii) show the same FOV area and, as such, share the same scale bar as the first micrograph in the series. Scale bars are 50 $\mu$m. Representative contrast-enhanced videos are provided in video S2.

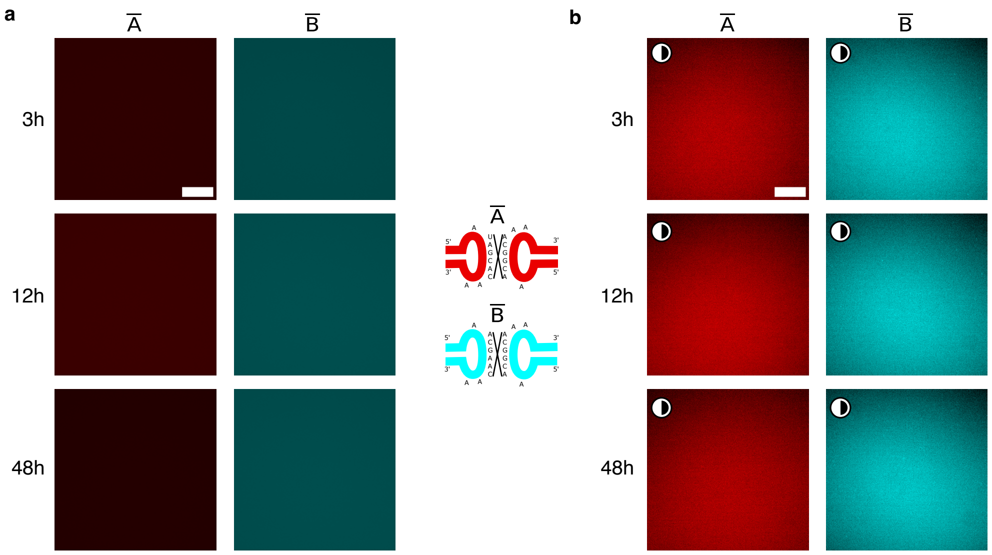


**Fig. S10 Non-sticky nanostars do not yield condensates.** Pristine (a) and contrast-enhanced (b, as highlighted by the half-shaded circle) epifluorescence micrographs of representative timepoints (3, 12, 48 h, along vertical axis) depicting the bulk transcription of RNA nanostars for control designs incapable of condensation ($\overline{\text{A}}$ and $\overline{\text{B}}$) due to scrambling of palindromic KL domains into non-binding sequences (loop sequences reported in the adjacent sketch). All frames in (a) depict the same FOV area, and, as such, share the same scale bar as the top left one. Same applies to (b). All timestamps are reported with respect to the start of timelapse imaging (see SI Methods, section 1.4.1 and Table S5). Scale bars are 50 $\mu$m. Representative contrast-enhanced videos are provided in video S1 (bottom).

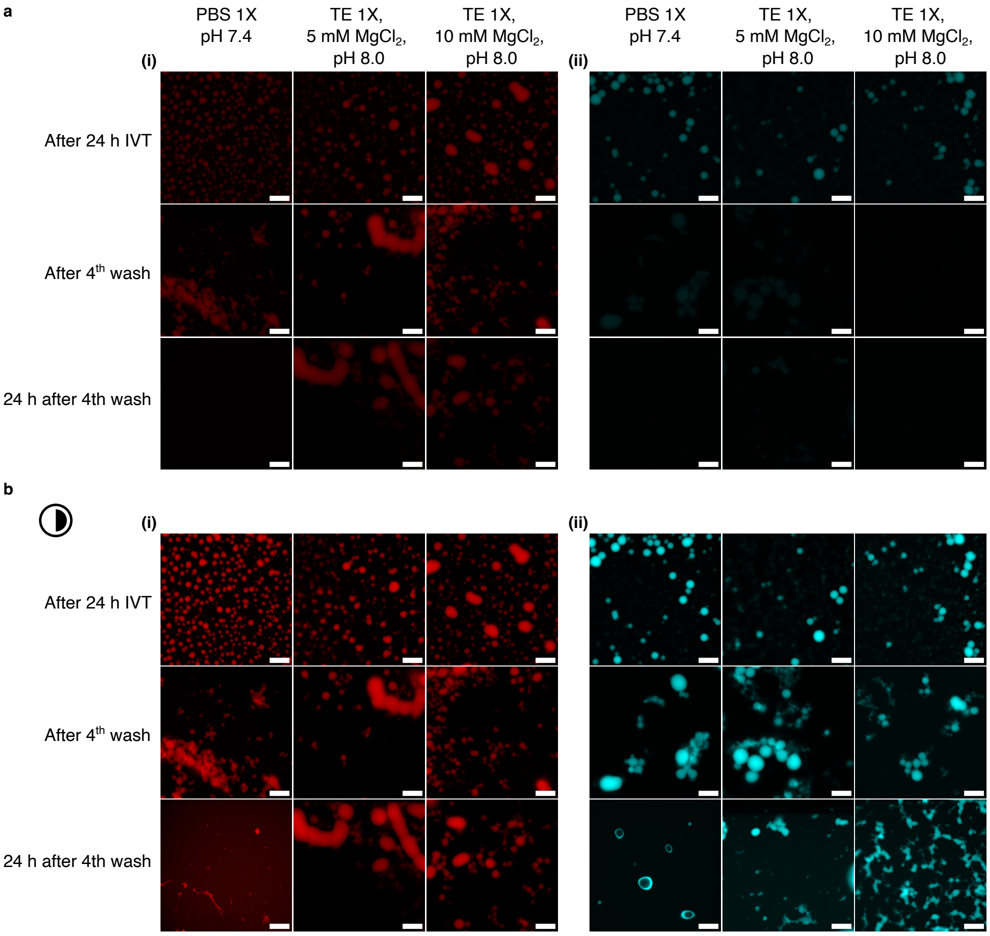


**Fig. S11 Stability of RNA condensates in different ionic conditions.** **(a)** Epifluorescence micrographs (reported as pristine, single equatorial planes) of (i) A and (ii) B RNA condensates were acquired after 24 h from start of bulk In Vitro Transcription (IVT) (top row: 'After 24 h IVT'). Samples underwent buffer exchange with either Phosphate-Buffered Saline (PBS) 1× pH 7.4 (left), Tris-EDTA (TE) 1× supplied with 5 mM $MgCl_2$ pH 8.0 (middle) or TE 1× supplied with 10 mM $MgCl_2$ pH 8.0 (right) through four washing cycles (see SI Methods, section 1.3.3) and imaged after the final wash (middle row: 'After $4^{th}$ wash'). Finally, condensate stability was inspected 24 h after the final wash (bottom row: '24 h after $4^{th}$ wash'). Notably, for these experiments condensates were formed in microwell plates rather than glass capillaries to enable buffer exchange (SI Methods, section 1.3.3), resulting in different condensate size and number after IVT. **(b)** Same as (a) after contrast enhancement, as highlighted by the half-shaded circle (0.2% pixels saturated). Scale bars are 100 $\mu$m.

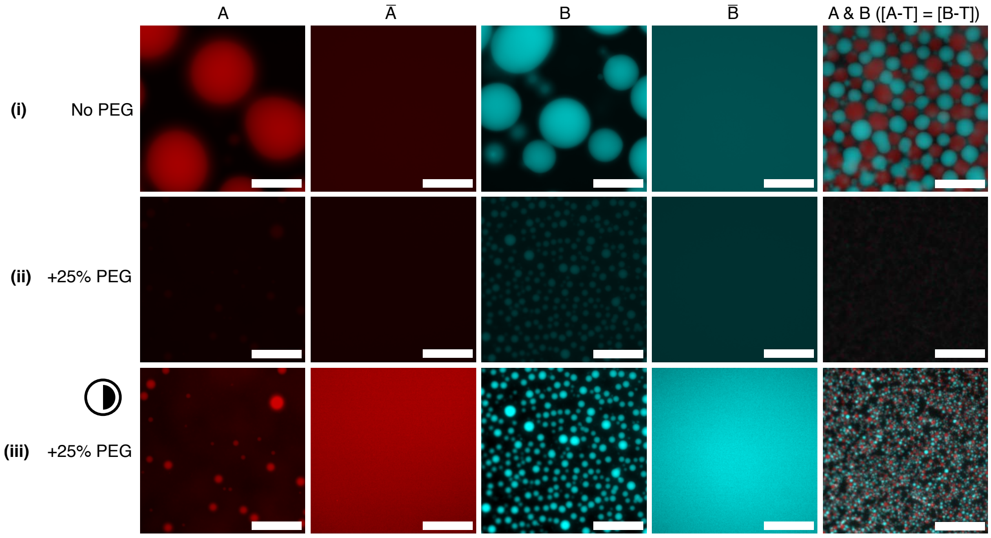


**Fig. S12 Influence of molecular crowding on bulk RNA condensate formation.** (i) In the absence of PEG, we observe condensate formation in sticky RNA motifs A and B, which remain distinct and demixed when co-transcribed (A & B). Non-sticky motifs only produce diffused fluorescence $\bar{\text{A}}$ and $\bar{\text{B}}$. (ii) In the presence of 25% volume/volume PEG 200, sticky nanostars still form condensates, albeit smaller ones, indicating that PEG 200 suppresses transcription rates [20]. In A & B mixtures, PEG induces adhesion of small A-type and B-type condensates and the formation of a percolating network. Non-sticky nanostructures remain soluble in the presence of PEG, but a decrease in fluorescence intensity compared to panel (i) confirms a lower transcription efficiency. (iii) Same micrographs presented in (ii) reported after contrast enhancement for ease of visualisation (0.20% pixels saturated). Micrographs in (i-ii) are pristine. All micrographs have been binned (4 × 4) down to 511 px × 512 px, and then cropped to a quarter of the original size. Samples were prepared as discussed in SI Methods, section 1.3.4. Scale bars are 100 $\mu$m.

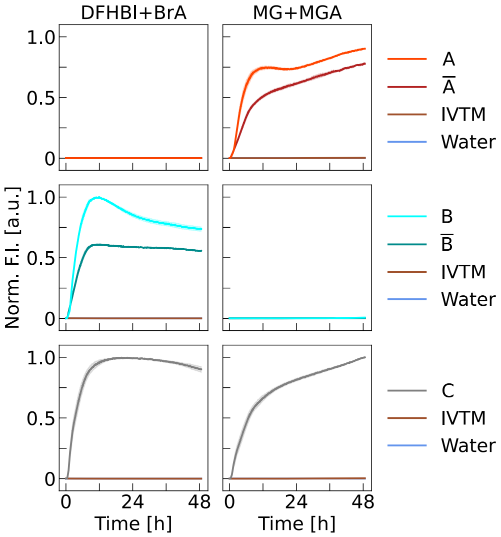


**Fig. S13 Fluorescence kinetics monitoring *in vitro* transcription and co-transcriptional folding of RNA nanostars.** Normalised fluorescence intensity (Norm. F.I.) increases with time, as RNA nanostars comprising functional fluorogenic aptamers (MGA in A and $\overline{\text{A}}$ – top, BrA in B and $\overline{\text{B}}$ – middle, both MGA and BrA in C – bottom) are synthesised. No increase is observed for controls lacking DNA templates (dyes in Water or IVTM - *In Vitro* Transcription Mixture, all panels, see SI Methods, section 1.5). Notably, sticky designs (A, B) exhibit a characteristic peak followed by a decrease in intensity, absent in non-sticky designs ($\overline{\text{A}}$, $\overline{\text{B}}$). This feature is likely due to fluorescent material being formed in the bulk (increase), condensating and sinking to the bottom of the wells, below the plane of focus (decrease). The different long-term behaviour among DFHBI+BrA (plateauing before 12 h) and MG+MGA (still increasing at 48 h) curves is likely related to differences in binding affinity and/or photophysics of the two dye-aptamer pairs, as it can be observed in designs with both one and two aptamers (C). Left/right column depicts normalised fluorescence intensity in the DFHBI+BrA/MG+MGA channel, respectively. Data are presented as normalised (see SI Methods, section 1.5) mean (solid lines) ± standard deviation (shaded regions) from triplicates.

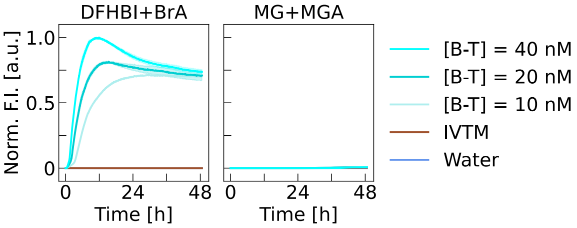


**Fig. S14 Effect of template concentration on the kinetics of RNA nanostar transcription in bulk.** Left/right column depicts normalised fluorescence intensity in the DFHBI+BrA/MG+MGA channel, respectively. Increasing the concentration (10, 20, 40 nM) of the DNA template coding for RNA nanostar B leads to faster fluorescence intensity increase in the DFHBI + BrA channel, related to the initial transcription rates, followed by an earlier and more prominent characteristic peak-decrease feature due to the concentration required for condensate formation being reached sooner. Data are presented as normalised (see SI Methods, section 1.5) mean (solid lines) ± standard deviation (shaded regions) from triplicates.

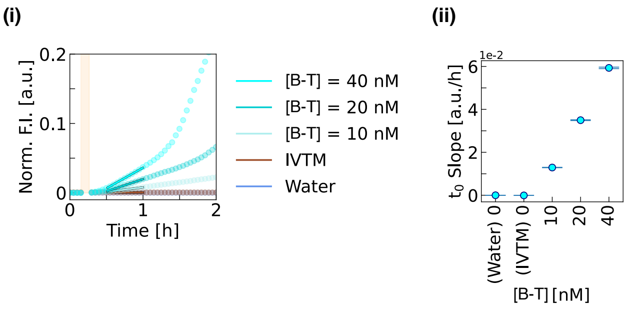


**Fig. S15 The initial increase rate in fluorescence intensity, reflecting the initial transcription rate of RNA nanostars (B) in bulk, increases monotonically with DNA template concentration ([B-T]).** (i) Zoomed inset of the kinetic profiles in Fig. S14 (DFHBI + Broccoli Aptamer channel only), with the injection window for DNA template coding for RNA nanostar B (in corresponding samples, no injection in negative controls, Water and IVTM for *In Vitro* Transcription Mixture, see SI Methods, section 1.5) shaded in light orange and timepoints used for linear fitting highlighted by surrounding boxes. Data are presented as normalised (see SI Methods, section 1.5) mean (circular markers) ± standard deviation (shaded regions) from triplicates. (ii) Initial fluorescence intensity increase rate evaluated by linear fitting of 30 timepoints (from normalised profiles in (i)) soon after the injection window. Data are presented as fitted slopes (markers) ± standard deviation from covariance matrix of the linear fit (error bars).

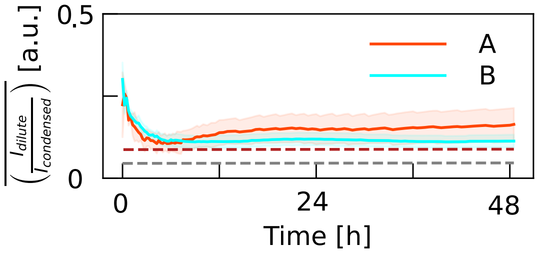


**Fig. S16 Time evolution of the ratio between the mean epifluorescence signal recorded in the bulk ($I_{\mathrm{dilute}}$) and within condensates ($I_{\mathrm{condensed}}$).** Data, extracted as outlined in the SI Methods section 1.4.2, are presented as sample mean (solid line) ± standard deviation (shaded region). The initial drop in the curves may be indicative of a transient in which the rate of RNA transcription exceeds that of condensate growth through monomer addition. However, the initial transient may be influenced by imaging or segmentation artefacts linked to the finite vertical resolution of the epifluorescence microscope or the possible presence of sub-diffraction condensates. Dashed horizontal lines indicate $I_{\mathrm{dilute}}/I_{\mathrm{condensed}}$ as determined from confocal micrographs collected at the end of the epifluorescence timelapse (Fig. 1b and Fig. S5). The difference between values extracted with epifluorescence and confocal is likely due to the greater focal depth of the former.

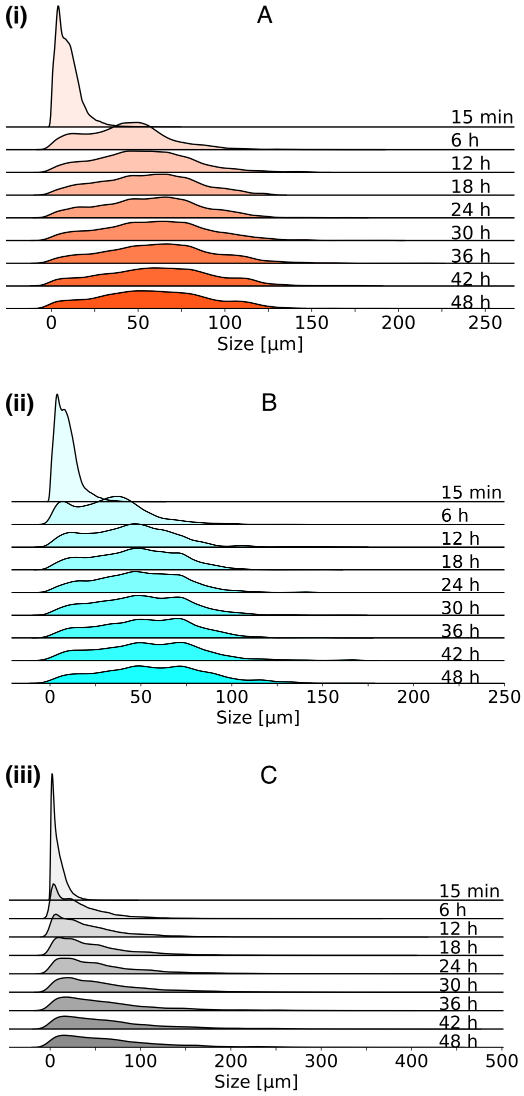


**Fig. S17 Ridge plot displaying the time evolution of the size of RNA condensates.** Size of condensates from RNA nanostars A (i), B (ii) and C (iii) in bulk is examined *via* the corresponding Chord-Length Distribution (CLD) at representative timepoints. The mean of these CLDs, $\mu_{\mathrm{CLD}}$, is used in Fig. 1d(i) as a proxy of the average size of the condensates. It can be observed that A and B form spherical condensates with a well defined CLD, mostly contained in the range 0-150 $\mu$m. Conversely, C forms extended mesh-like assemblies, and the corresponding CLD is very spread-out, surpassing 250 $\mu$m (range of x axis is dictated by the most spread out timepoint of each sample CLD).

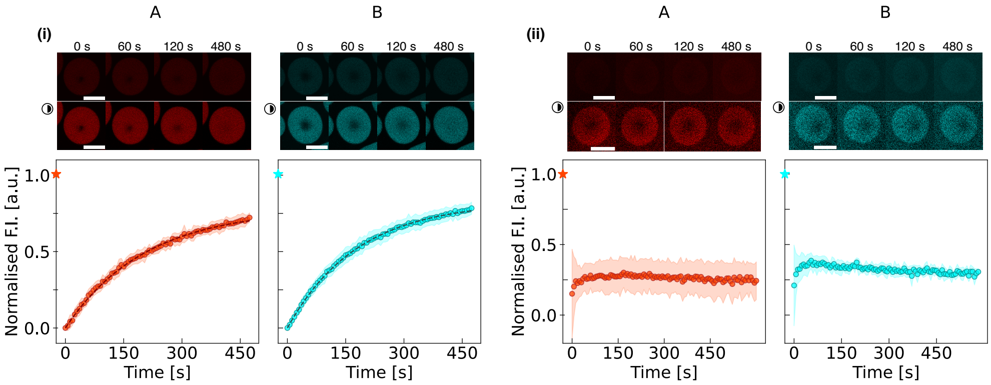


**Fig. S18 Fluorescence Recovery After Photobleaching (FRAP) of RNA condensates.** (i) FRAP recovery curves for A- and B-type condensates obtained by bleaching the embedded FLAPs: MGA for A-type condensates and BrA for B-type condensates. Exponential recovery is observed for both, with timescales $\tau_f = 204 \pm 5\,\mathrm{s}$ for A and $\tau_f = 180 \pm 3\,\mathrm{s}$ for B. The fitting function used is $A\,[1 - \exp(-t/\tau_f)]$. These recovery timescales are fast compared to observed droplet coalescence timescales (Fig. 1e and f), and recovery may be caused by exchange of bleached MG/DFHBI molecules with the bulk, reported to occur over comparable timescales for DFHBI variants DFHBI-1T and BI [21]. Curves are normalised between 0 and 1, using the first post-bleach intensity datum and the pre-bleach intensity (indicated by a star) as references. Data are shown as mean ± standard deviation of recovery curves collected on 25 (A) and 27 (B) condensates in 3 fields-of-view *per* sample. Top: representative confocal snapshots of the condensates (native and contrast-enhanced). (ii) Recovery curves obtained by bleaching fluorescein dyes covalently linked to the RNA nanostars, embedded by including fluorescein-labelled UTP in the transcription mixture (see SI Methods, section 1.3.5). No recovery is observed over the experimental timescales, confirming dye exchange as the likely cause of the recovery seen in panel (i). The slight downward trend is ascribed to bleaching during the recovery phase, while the initial quick recovery is ascribed to unbound fluorescein-UTP. Data are normalised using the pre-bleach intensity only (star symbol), and shown as mean ± standard deviation of recovery curves collected on 7 condensates in one field-of-view. Top: pristine and contrast-enhanced representative micrographs. See SI Methods, section 1.4.4 for experimental and image segmentation details.

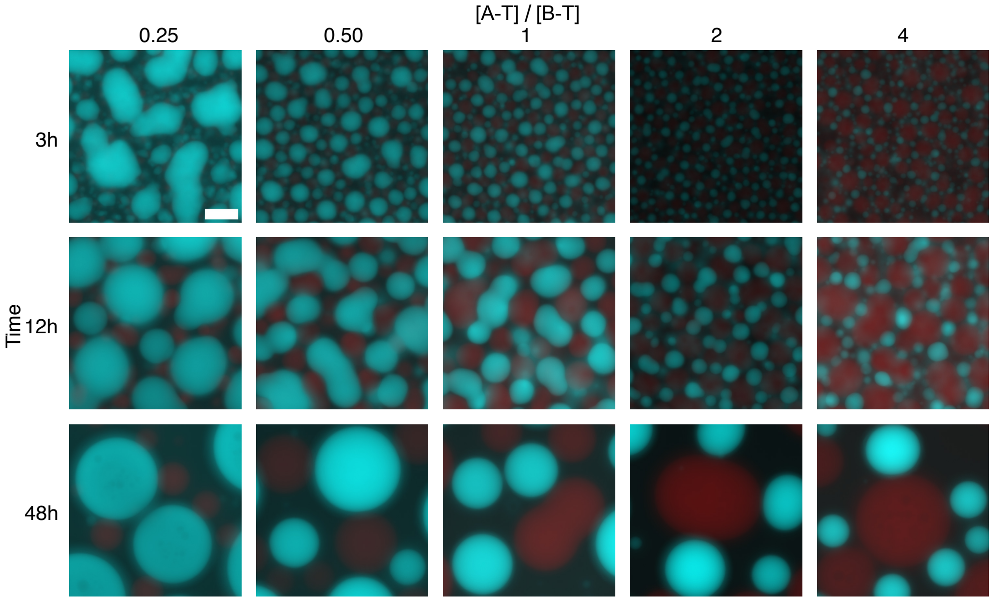


**Fig. S19 Binary A/B systems form orthogonal RNA condensates in bulk, with size tunable *via* the DNA template concentration ratios.** Pristine epifluorescence micrographs of representative timepoints (3, 12, 48 h, along vertical axis) depicting the bulk self-assembly of binary systems with different [A-T]/[B-T] composition ratios (from 0.25 to 4, horizontal axis). Timestamps are reported with respect to the start of timelapse imaging (see SI Methods, section 1.4.1 and Table S5). These micrographs correspond to those in Fig. 2a (contrast-enhanced). All micrographs depict the same FOV area and, as such, share the same scale bar (top left), which is 50 $\mu$m. Representative contrast-enhanced videos are provided in video S3.

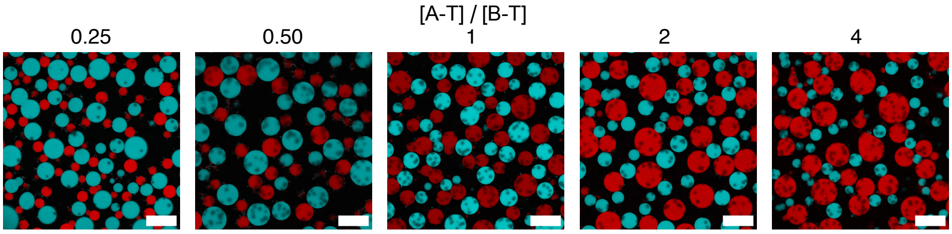


**Fig. S20 Confocal micrographs depicting orthogonal condensate formation in A/B systems with varying DNA template concentration ratios.** Samples ([A-T]/[B-T] composition ratios from 0.25 to 4, horizontal axis) were imaged after more than 48 h. The black dots are likely inorganic pyrophosphate precipitates or non-specific aggregates that are often observed in *in vitro* transcription mixtures after tens of hours. Micrographs are pristine composites of MG and DFHBI channels, and have been binned down to the desired size (1024 px × 1024 px). Scale bars are 100 $\mu$m.

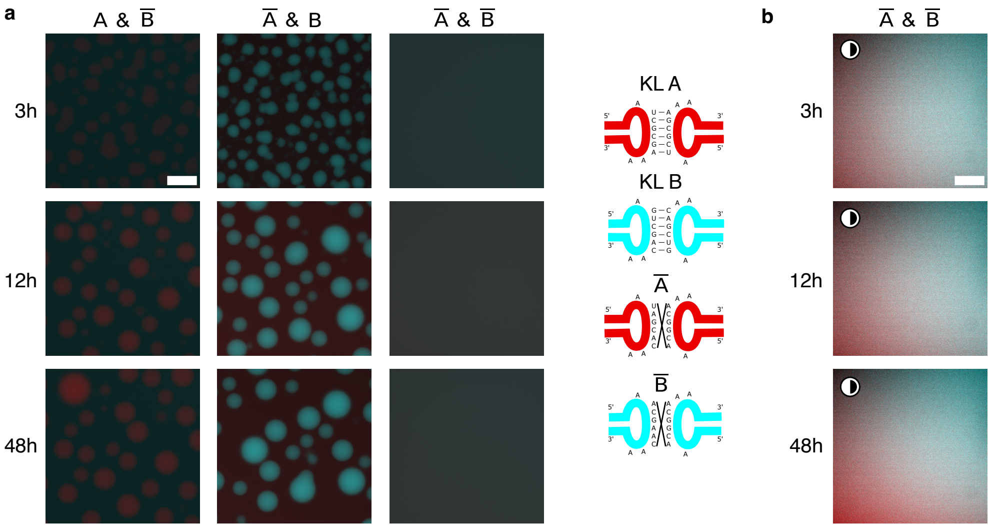


**Fig. S21 KL programmability enables control over condensate formation, or lack thereof, in binary systems with non-sticky nanostars.** (a) Pristine epifluorescence micrographs of representative timepoints (3, 12, 48 h, along vertical axis) depicting the bulk self-assembly of control binary mixtures (1:1 ratio) of A and B where either or both are incapable of condensation (indicated with a bar, i.e. $\overline{\mathrm{A}}$, $\overline{\mathrm{B}}$) due to scrambling of palindromic KL domains into non-binding sequences, as highlighted in sketches (left) of original KL A/B and their scrambled counterparts. Micrographs in the left (A & $\overline{\mathrm{B}}$) and middle ($\overline{\mathrm{A}}$ & B) columns correspond to those in Fig. 2b (contrast-enhanced). (b) Contrast-enhanced (as highlighted by the half-shaded circle) micrographs from the right-most column of A ($\overline{\mathrm{A}}$ & $\overline{\mathrm{B}}$). Micrographs in (a) all show the same FOV area as the top left micrograph and, as such, share the same scale bar. Same applies to (b). All timestamps are reported with respect to the start of timelapse imaging (see SI Methods, section 1.4.1 and Table S5). Scale bars are 50 $\mu$m. Representative contrast-enhanced videos are provided in video S4.

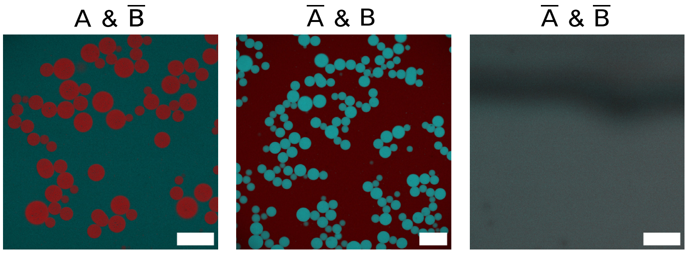


**Fig. S22 Confocal micrographs depicting control over condensate formation *via* KL programmability in binary A/B systems with non-sticky nanostars.** Samples with either or both non-sticky nanostar variants (indicated with a bar, *i.e.* A $\rightarrow$ $\overline{\mathrm{A}}$) were imaged after more than 48 h. Micrographs are pristine composites of MG and DFHBI channels, and have been binned down to the desired size (1024 px $\times$ 1024 px). Scale bars are 100 $\mu$m.

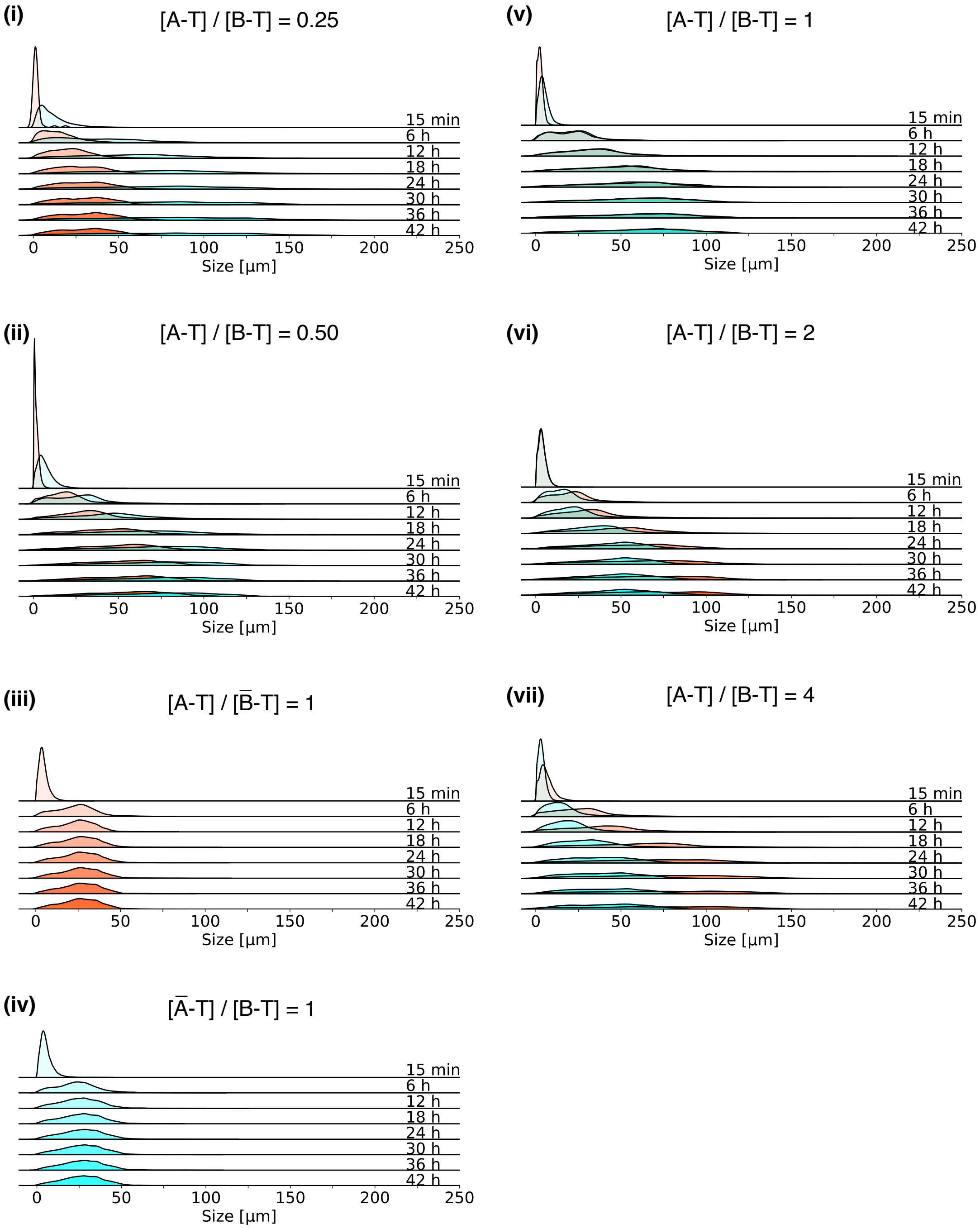


**Fig. S23 Ridge plot displaying the time evolution of the size of RNA condensates in binary mixtures in bulk.** Size of condensates formed in binary systems with at least one sticky nanostar species is investigated *via* Chord-Length Distribution (CLD) at representative timepoints. The mean of these CLDs, $\mu_{\mathrm{CLD}}$, is used in Fig. 2c as proxy of the average size of the condensates. DNA template concentration ratios are reported on top.

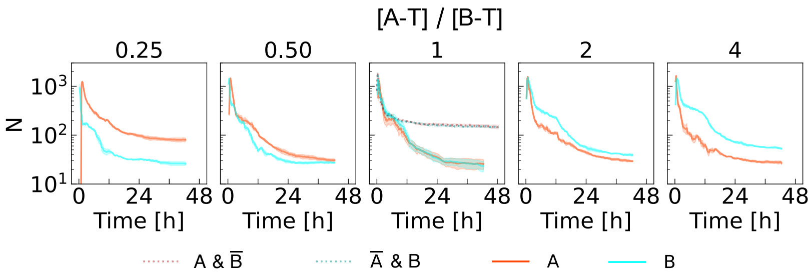


**Fig. S24 Template-concentration ratio and nanostar stickiness influence condensate coarsening kinetics.** Time evolution of the number ($N$) of A (red) and B (cyan) condensates in a microscopy FOV (see SI Methods, sections 1.4.2) for varying DNA-template concentration ratios ([A-T]/[B-T]). Higher relative template concentrations promote faster coarsening (smaller $N$). Two-step coarsening kinetics is noted and ascribed to transient jamming (see main text discussion), similar to the trends shown in Fig. 2c for condensate size. For [A-T] = [B-T], the condensate-forming A & B system is compared with systems where one nanostar type is rendered non-sticky ($\overline{\text{A}}$ & B and A & $\overline{\text{B}}$, dotted lines). For $\overline{\text{A}}$ and A & $\overline{\text{B}}$, $N$ plateaus at higher values compared to A & B, suggesting that, at long times, cooperativity between orthogonal condensates facilitates coalescence.

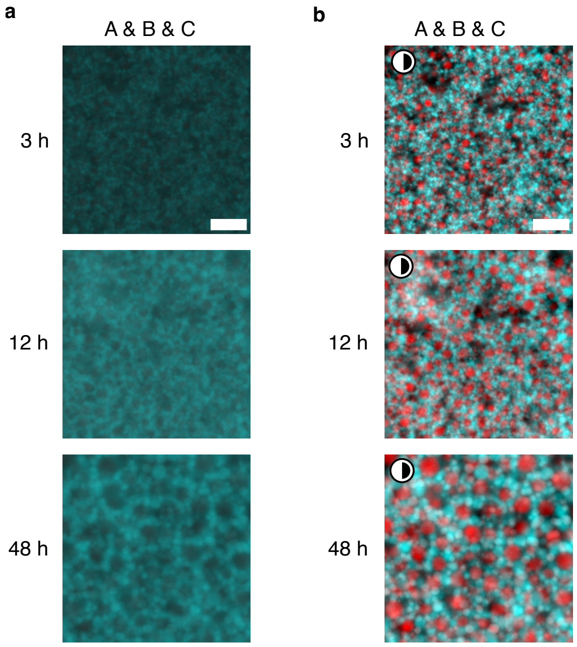


**Fig. S25 Bulk self-assembly of condensates in the ternary system.** (a) Pristine and (b) contrast-enhanced (as highlighted by the half-shaded circle) epifluorescence micrographs of representative timepoints (3, 12 and 48 h, along vertical axis) depicting the bulk self-assembly of condensates in the stoichiometric ternary system (A-T:B-T:C-T = 1:1:1). Due to the higher stiffness of construct C and the simultaneous presence of multiple orthogonal RNA nanostars, relaxation times are much longer and, even after 48 h, the sample appears as a network comprising a much larger number of smaller condensates than what we observe in single or binary component systems. Micrographs in (a) all show the same FOV area as the first micrograph of the series (3 h) and, as such, share the same scale bar. Same applies to (b). All timestamps are reported with respect to the start of timelapse imaging (see SI Methods, section 1.4.1 and Table S5). Scale bars are 50 $\mu$m. Representative contrast-enhanced videos are provided in video S5.

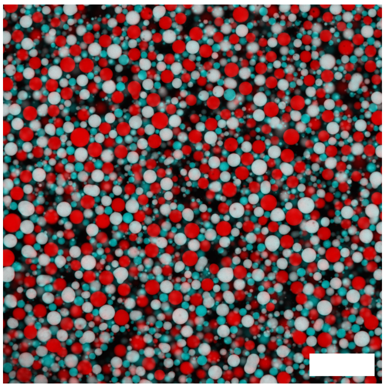

**Fig. S26 Confocal micrographs depicting formation of three distinct kinds of orthogonal RNA condensates in ternary systems.** Samples were imaged after more than 48 h, allowing for further relaxation of the condensates. Three different kinds of condensates (red - A nanostars -, cyan - B nanostars - or white - C nanostars -, *i.e.* both red and cyan) can now be spotted much more clearly than in Fig. S25. Micrographs are pristine composites of MG and DFHBI channels, and have been binned down to the desired size (1024 px × 1024 px). Scale bars are 100 $\mu$m.

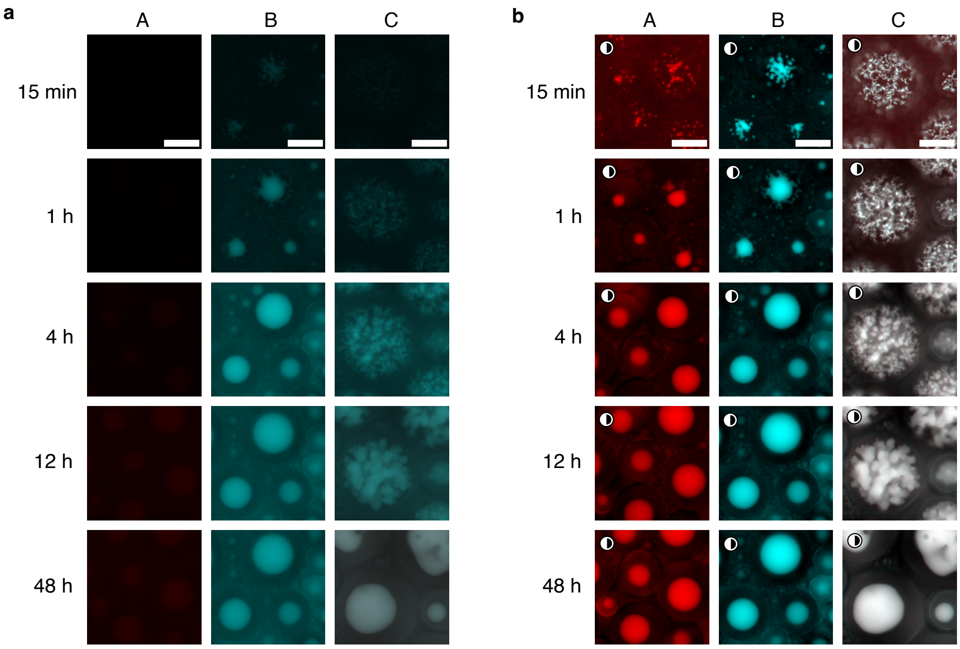


**Fig. S27 Self-assembly of RNA organelles within synthetic cells.** (a) Pristine epifluorescence micrographs of representative timepoints (15 min, 1 h, 4 h, 12 h and 48 h, along vertical axis) depicting the self-assembly within synthetic cells (water-in-oil droplets) of organelles from constructs A, B and C, respectively. Micrographs for timepoints at 15 min and 3 h (or 4 h for C) correspond to those in Fig. 3b. (b) Micrographs from panel (a) after contrast-enhancement (as highlighted by the half-shaded circle). C samples were imaged in a separate run which started 15 min earlier with respect to template mixing and encapsulation compared to A and B. For this reason, the second available timepoint was used to represent 15 min for A and B, while the first one (acquired 15 min after the start of the run) was used for C for ease of comparison (timestep in the first 10 h is equal to 15 min). See SI Methods section 1.4.1 and Table S6 for detailed information on the time interval between template mixing and start of timelapse imaging.. All micrographs for each sample (column) in A and B show the same FOV area, thus sharing scale bar with the first micrograph in the series. Scale bars are 50 $\mu$m. Representative contrast-enhanced videos are provided in video S6 (top).

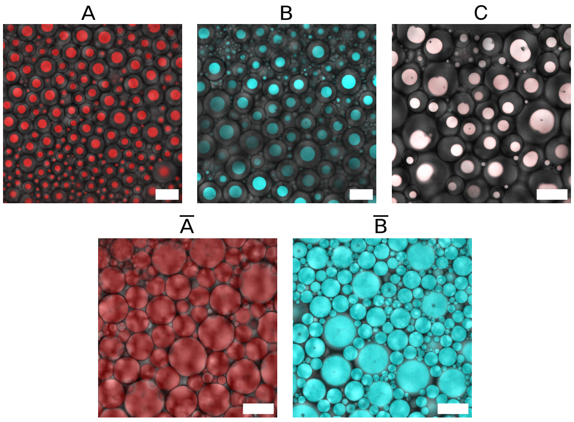


**Fig. S28 Confocal micrographs depicting organelle formation, or lack thereof, in samples of co-transcriptionally folding sticky and non-sticky RNA nanostars within synthetic cell mimics.** Samples were imaged after more than 48 h. Micrographs are pristine composites of bright-field, MG and DFHBI channels, and have been binned down to the desired size (1024 px $\times$ 1024 px). Scale bars are 100 $\mu$m.

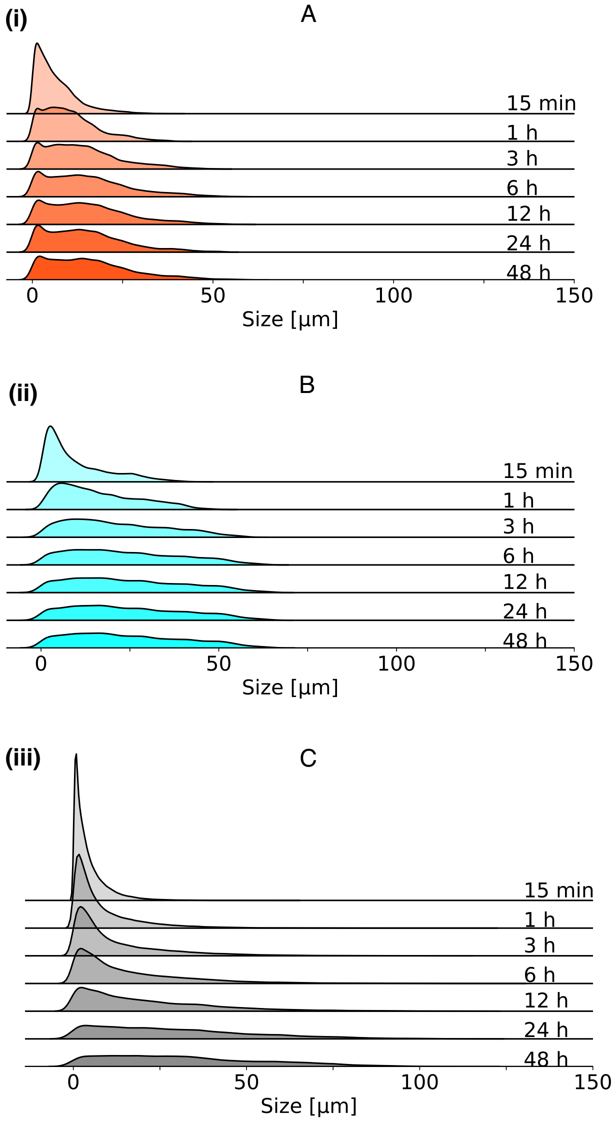


**Fig. S29 Ridge plot displaying the time evolution of the size of RNA organelles in synthetic cells.** Size of organelles formed by RNA nanostars A (i), B (ii) and C (iii) in synthetic cells (water-in-oil droplets) is examined *via* the corresponding Chord-Length Distribution (CLD) at representative timepoints. The mean of these CLDs, $\mu_{\mathrm{CLD}}$, is used in Fig. 3c as a proxy of the average size of the condensates. It can be observed how A and B readily reach a steady-state size distribution (6 h timepoint onwards), while C relaxes more slowly. In all samples, CLDs are more spread out compared to the bulk case (*e.g.* looking at the ratio between the mean and the standard deviation) due to a wide range of droplet sizes in the imaged FOVs.

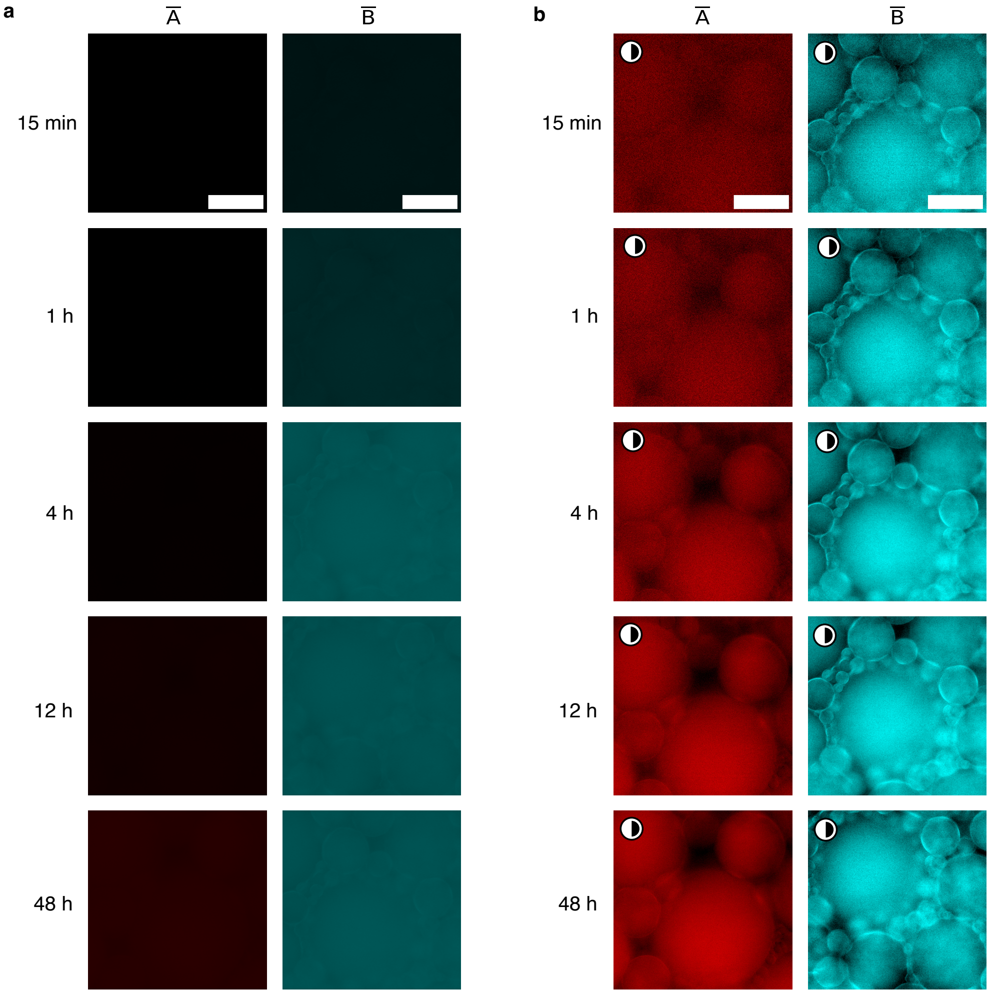


**Fig. S30 Non-sticky nanostars do not yield organelles in synthetic cells.** (a) Pristine epifluorescence micrographs of representative timepoints (15 min, 1 h, 4 h, 12 h and 48 h, along vertical axis) depicting the self-assembly within synthetic cells (water-in-oil droplets) of condensates (or, in this case, lack thereof) from non-sticky nanostar variants (scrambled KL domains). (b) Micrographs from panel (a) after contrast-enhancement (as highlighted by the half-shaded circle). All micrographs for each sample (column) show the same FOV area and, as such, share the same scale bar as the first micrograph in the series in both (a) and (b). All timestamps are reported with respect to the start of timelapse imaging (see SI Methods, section 1.4.1 and Table S6). Scale bars are 50 $\mu$m. Representative contrast-enhanced videos are provided in video S6 (bottom).

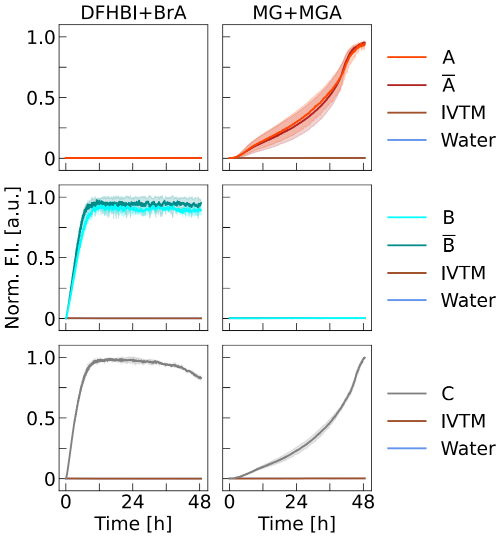


**Fig. S31 Fluorescence kinetics monitoring transcription and folding of RNA nanostars within synthetic cells.** Left/right column depicts normalised fluorescence intensity in the DFHBI+BrA/MG+MGA channel, respectively. No increase is observed for negative controls lacking DNA templates (Water and IVTM - *In Vitro* Transcription Mixture, see SI Methods, section 1.5). Different from bulk measurements (Fig. S13), in synthetic cells we observe no difference between samples with and without palindromic KLs (A, B vs $\overline{\text{A}}$, $\overline{\text{B}}$). This supports the hypothesis ascribing the KL-related peak-decrease feature to condensates forming and sinking in the bulk volume, which does not take place within confined emulsion droplets. Signals from nanostars bearing MGA (A, $\overline{\text{A}}$, C) display a delayed multi-stage increase, which does not correspond to a delay in condensate growth (Fig. 3c). We argue that the trend emerges from the affinity of MG for hydrophobic phases, previously observed in Ref. [12] and supported by the calculated n-octanol/water partitioning (logP) in Ref. [9]. Therefore, MG initially partitions in the oil phase and then slowly accumulates within the condensates by binding to the MGA motifs. Data are presented as normalised (see Methods, section 1.5) mean (solid lines) ± standard deviation (shaded regions) from triplicates.

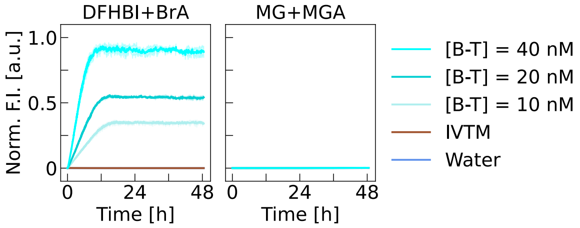


**Fig. S32 Effect of template concentration on the kinetics of RNA nanostar transcription in synthetic cells.** Left/right column depicts normalised fluorescence intensity in the DFHBI+BrA/MG+MGA channel, respectively. Increasing the concentration of the DNA template coding for RNA nanostar B ([B-T] = 10, 20 or 40 nM) leads to faster fluorescence intensity increase in the DFHBI+BrA channel, related to the initial transcription rates, as well as earlier plateauing. Higher DNA template concentrations also lead to higher plateaus, indicating a larger yield of RNA transcripts. Data are presented as normalised (see SI Methods, section 1.5) mean (solid lines) ± standard deviation (shaded regions) from triplicates.

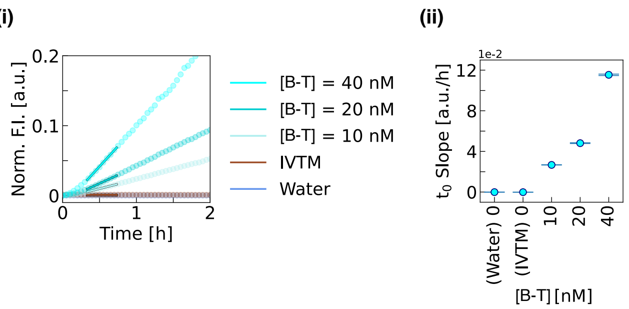


**Fig. S33 The initial increase rate in fluorescence intensity, reflecting the initial transcription rate of RNA nanostars (B) in synthetic cells, increases monotonically with DNA template concentration ([B-T]).** (i) Zoomed inset of the kinetic profiles in Fig. S32 (DFHBI+BrA channel only), with timepoints used for linear fitting highlighted by surrounding boxes. Data are presented as normalised (see SI Methods, section 1.5) mean (circular markers) ± standard deviation (shaded regions) from triplicates. (ii) Initial fluorescence intensity increase rate evaluated by linear fitting of 30 timepoints (from normalised profiles in (i)) soon after the start of the measurement run. Such values are approximately twice as large as those in bulk in Fig. S15. As DNA templates are injected prior to emulsification, *i.e.* before starting the measurement run, the fitted time window might not be equivalent to the one used for bulk measurements. This might in turn lead to the observed value discrepancy. Data are presented as fitted slopes (markers) ± standard deviation from covariance matrix of the linear fit (error bars).

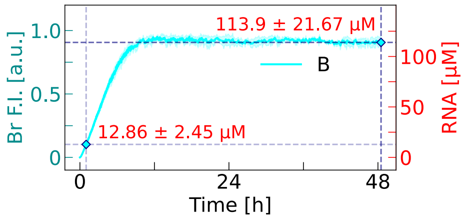


**Fig. S34 Estimating RNA nanostar synthesis yield within synthetic cells through calibration of fluorescence intensity kinetics.** Normalised fluorescence intensity in the DFHBI channel (Br F.I., left y-axis) *vs* time for RNA nanostar B within synthetic cells, as shown in Fig. S31. Fluorescence intensity kinetics for RNA nanostar B were calibrated with respect to the RNA concentration in micromolar units (RNA [$\mu$M], right y-axis) by estimating the average endpoint nanostar concentration within synthetic cells as outlined in SI Methods, section 1.5. Two markers indicate the RNA concentration within synthetic cells at 1 h after the start of the run (approximately 1 h and 30 min after mixing the DNA templates, *i.e.* start of transcription, see SI Methods section 1.4.1 and Table S6) and at 48.5 h (endpoint). The first timepoint was chosen as a proxy for the 15 min timepoint in the corresponding epifluorescence timelapses of co-transcriptional self-assembly within synthetic cells, where the run started 1 h and 17 min after injection of the DNA templates in the transcription mixture (Fig. 3b and Table S6). Evidence of aggregate formation at this time-point, where we estimate the RNA concentration to be $12.86 \pm 2.45\,\mu$M (equivalent to a mass RNA concentration of $1.07 \pm 0.20$ g L$^{-1}$), is consistent with previous experimental reports for DNA nanostars [22, 23].

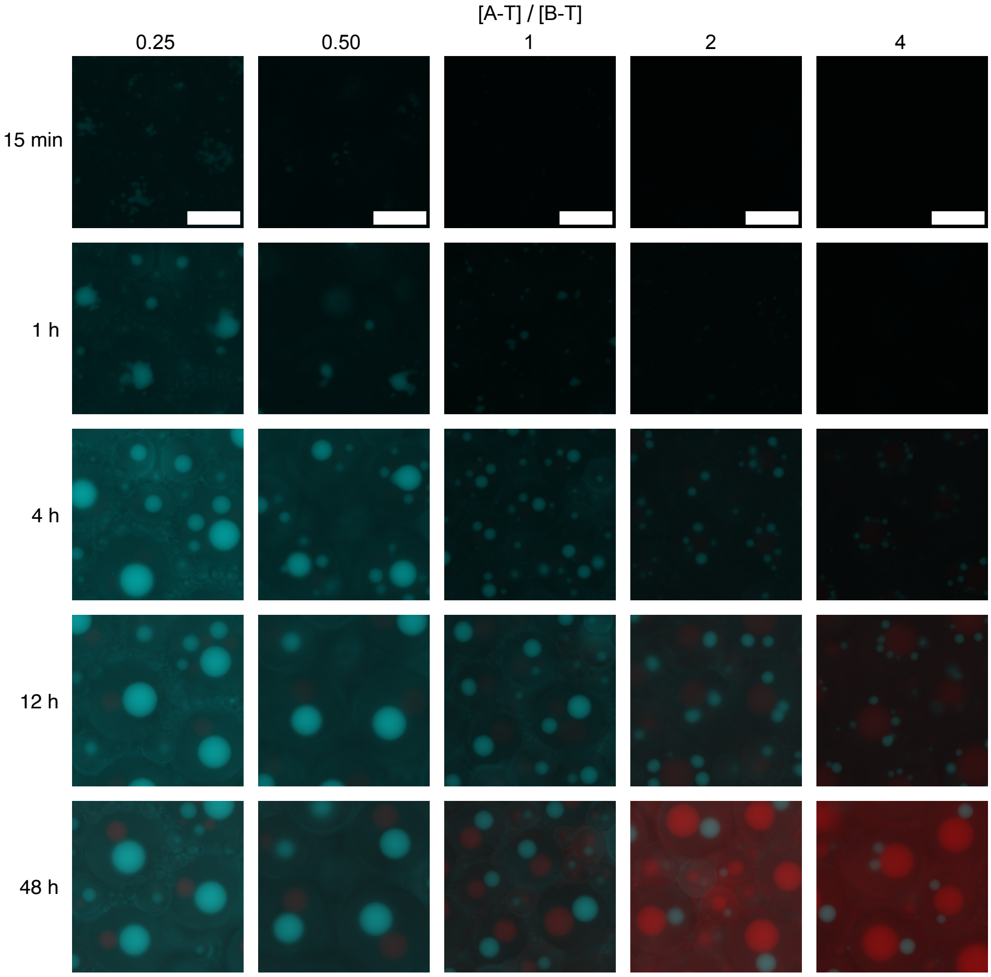


**Fig. S35 Binary A/B systems form orthogonal RNA organelles with tunable relative size within synthetic cells.** Pristine epifluorescence micrographs of representative timepoints (15 min, 1 h, 4 h, 12 h and 48 h, along vertical axis) depicting the self-assembly within synthetic cell mimics (water-in-oil droplets) of binary condensate systems with different [A-T]/[B-T] composition ratios (from 0.25 to 4, along the horizontal axis). All micrographs for each sample (column) show the same FOV area and, as such, share the same scale bar as the first micrograph in the series. Timestamps are reported with respect to the start of timelapse imaging (see SI Methods, section 1.4.1 and Table S6). Scale bars are 50 $\mu$m. Representative contrast-enhanced videos are provided in video S7.

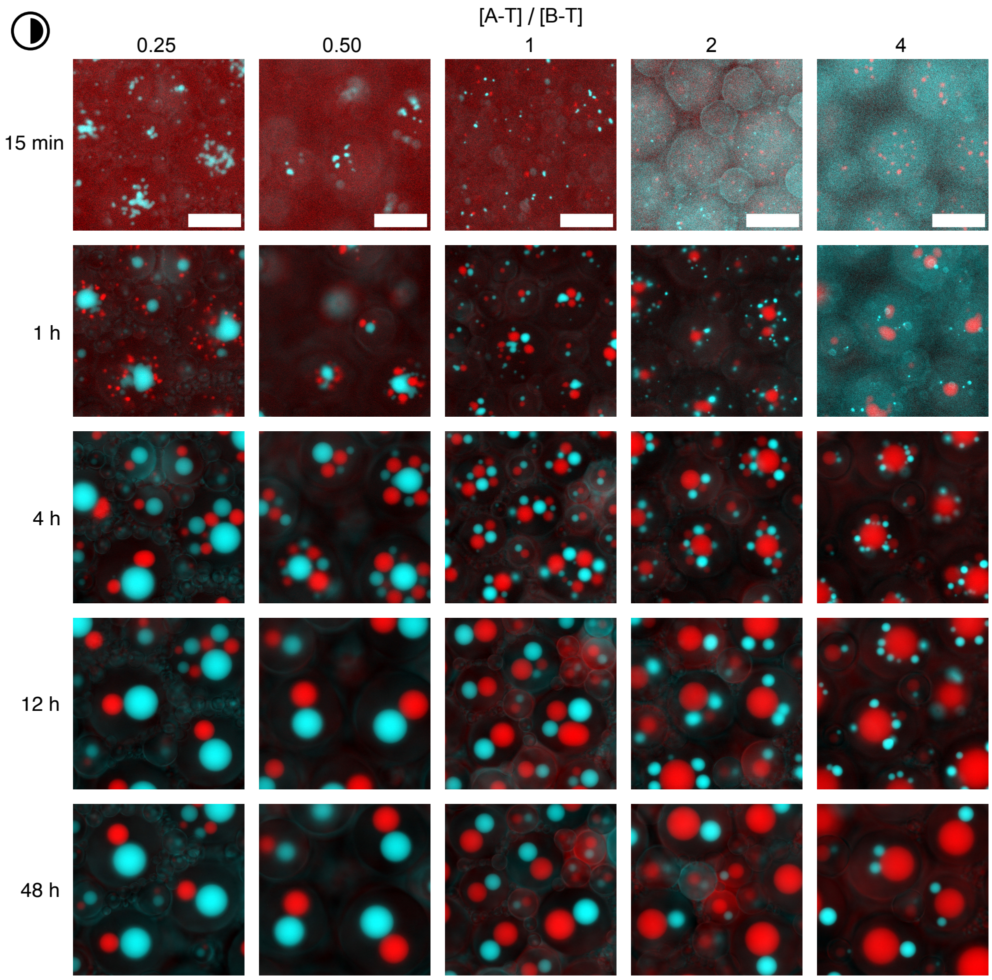


**Fig. S36 Binary A/B systems form orthogonal RNA organelles with tunable relative size within synthetic cells (processed).** Contrast-enhanced (as indicated by the half-shaded circle) epifluorescence micrographs corresponding to pristine ones in Fig. S35. All micrographs for each sample (column) show the same FOV area and, as such, share the same scale bar as the first micrograph. Timestamps are reported with respect to the start of timelapse imaging (see SI Methods, section 1.4.1 and Table S6). Scale bars are 50 $\mu$m. Representative contrast-enhanced videos are provided in video S7.

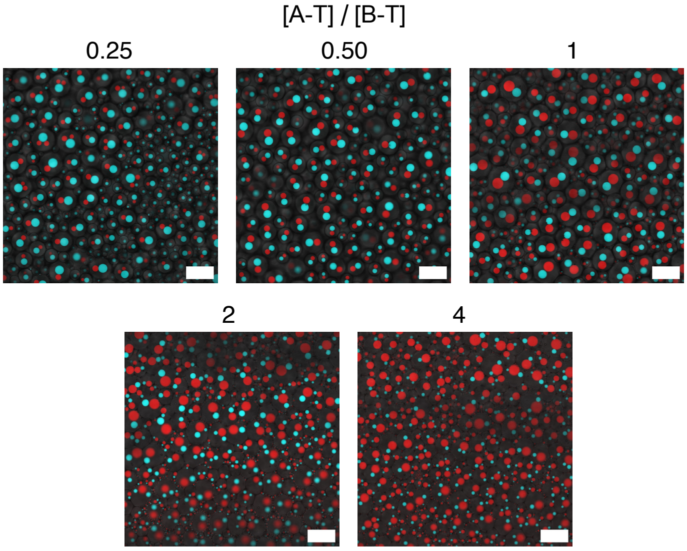


**Fig. S37 Confocal micrographs depicting the orthogonal RNA organelles with tunable relative size formed by binary A/B systems in synthetic cells.** Samples ([A-T]/[B-T] composition ratios from 0.25 to 4, horizontal axis) were imaged after more than 48 h. Micrographs are pristine composites of bright-field, MG and DFHBI channels, and have been binned down to the desired size (1024 px × 1024 px). Scale bars are 100 $\mu$m.

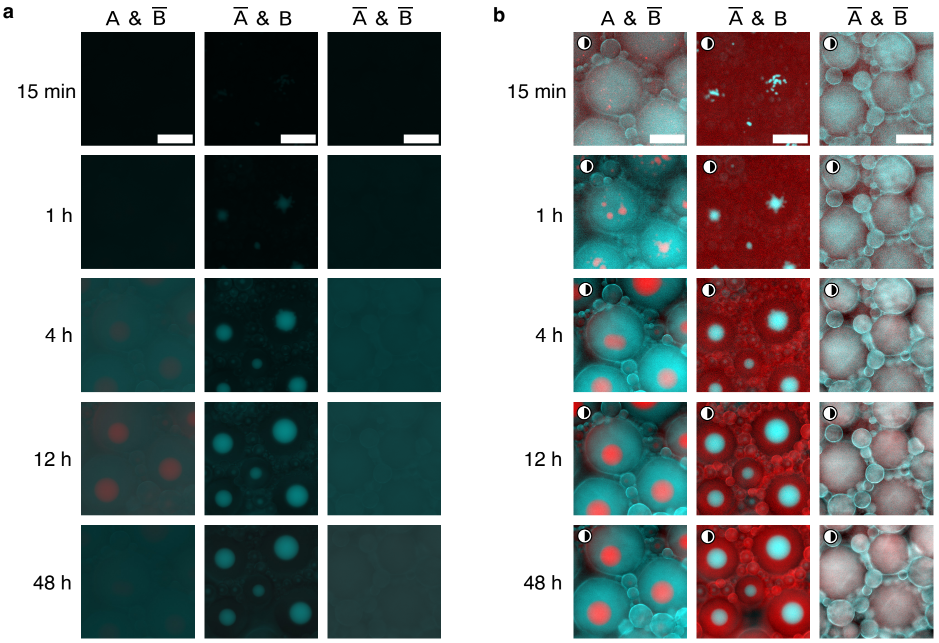


**Fig. S38 KL programmability enables control over organelle formation, or lack thereof, in binary systems with non-sticky nanostars within synthetic cells.** (a) Pristine epifluorescence micrographs of representative timepoints (15 min, 1 h, 4 h, 12 h and 48 h, along vertical axis) depicting the self-assembly within synthetic cells (water-in-oil droplets) of control binary mixtures (1:1 ratio) of A and B where either or both are non-sticky (scrambled KL domains). (b) Micrographs from (a) after contrast-enhancement (as highlighted by the half-shaded circle). Micrographs in (a) and (b) all show the same FOV area as the top left micrograph and, as such, share the same scale bar. All timestamps are reported with respect to the start of timelapse imaging (see SI Methods, section 1.4.1 and Table S6). Scale bars are 50 $\mu$m. Representative contrast-enhanced videos are provided in video S8.

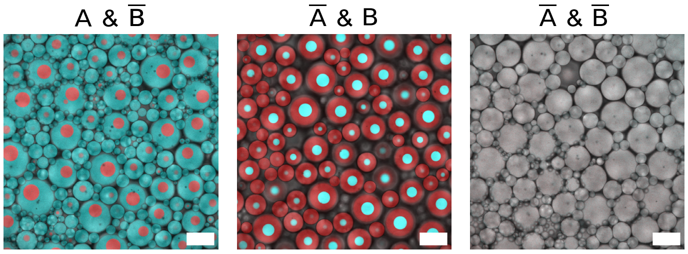


**Fig. S39 Confocal micrographs depicting control over condensate formation *via* KL programmability in binary A/B systems with non-sticky nanostars transcribed within synthetic cells.** Pristine confocal micrographs (composite, including bright-field) representing condensates (or lack thereof) formed in synthetic cells (water-in-oil droplets) by binary control mixtures of A and B where either or both are non-sticky (scrambled KL domains, indicated with a bar, *i.e.* $\overline{\text{A}}$, $\overline{\text{B}}$). Samples were imaged after more than 48 h. Micrographs have been binned down to the desired size (1024 px × 1024 px). Scale bars are 100 $\mu$m.

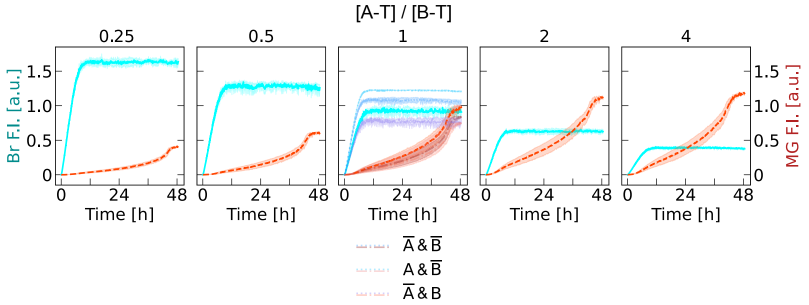


**Fig. S40 Fluorescence kinetics monitoring *in vitro* transcription and co-transcriptional folding of orthogonal A and B RNA nanostars in binary systems with varying DNA template concentration ratio within synthetic cells (water-in-oil droplets).** Samples were incubated at 30 °C. Left/right y-axis depicts normalised fluorescence intensity in the DFHBI+BrA (Br F.I., cyan label) / in the MG+MGA (MG F.I., red label) channel in solid/dashed line curves, respectively (see SI Methods, section 1.5). There is a clear trend in both channels: when the corresponding DNA template concentration increases (left to right for A, right to left for B), the fluorescence signal increases faster, plateaus earlier and reaches a higher relative value when normalised with respect to the stoichiometric condition (middle panel, [A-T]:[B-T] = 1:1). Controls with either or both non-sticky components (in 1:1 ratio) are included in the middle panel and do not show clear differences compared to the case of both sticky components, further proving that KL interactions do not impede any of the aptamers from correctly binding the respective fluorogenic dyes. Data are presented as normalised (see SI Methods, section 1.5) mean (solid/dashed lines) ± standard deviation (shaded regions) from triplicates.

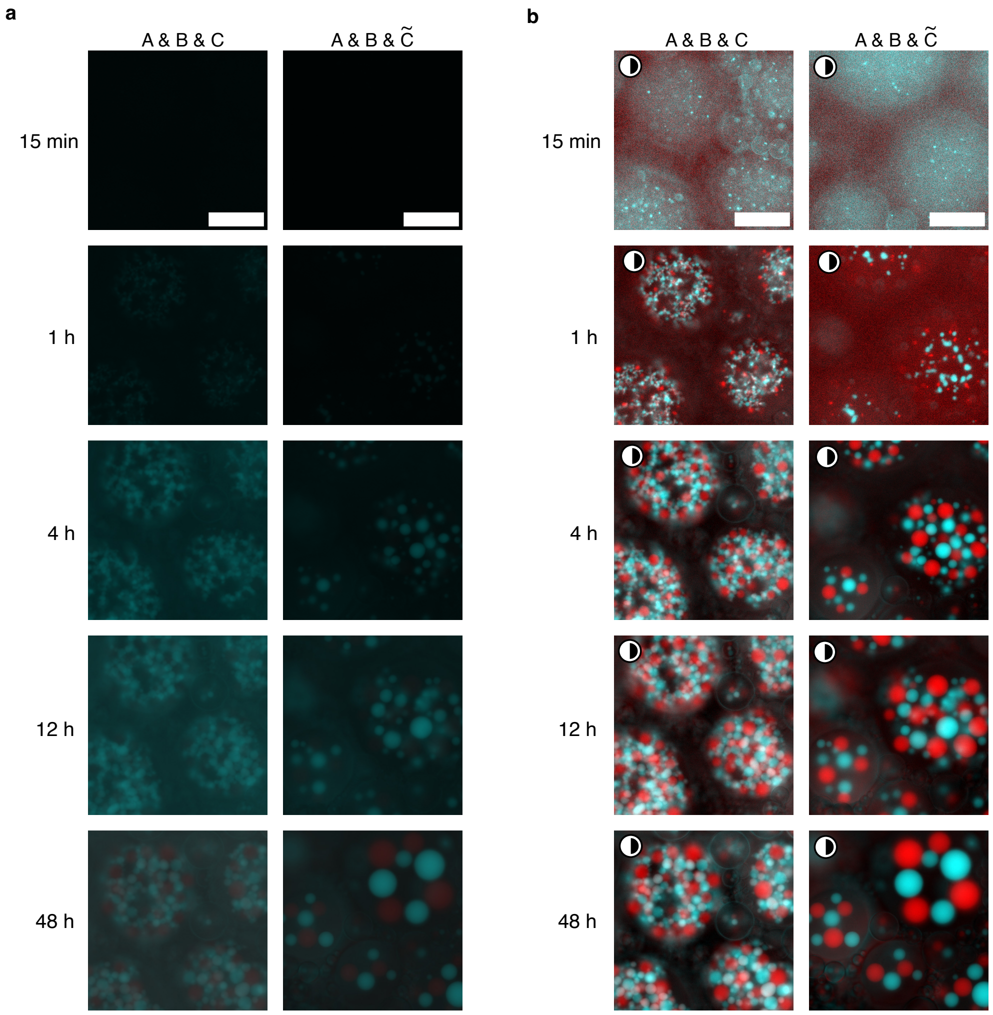


**Fig. S41 Self-assembly of organelles in ternary systems within synthetic cells.** (a) Pristine epifluorescence micrographs of representative timepoints (15 min, 1 h, 4 h, 12 h and 48 h, along vertical axis) depicting the self-assembly within synthetic cells (water-in-oil droplets) of ternary condensate systems ([A-T]:[B-T]:[C-T]([$\tilde{C}$-T]) = 1:1:1(1)). $\tilde{C}$ indicates a nanostar with the same KL as C but lacking any fluorogenic aptamer: as such, it is not visible on fluorescent channels, but its presence can be clearly spotted thanks to gaps between orthogonal red (A) and cyan (B) condensates. The stiffness of construct C leads to much longer relaxation times and a much larger number of smaller condensates within each droplet at the 48 h timepoint. (b) Micrographs from (a) after contrast-enhancement (as highlighted by the half-shaded circle). Micrographs in (a) and (b) all show the same FOV area as the top left micrograph and, as such, share the same scale bar. All timestamps are reported with respect to the start of timelapse imaging (see SI Methods, section 1.4.1 and Table S6). Scale bars are 50 $\mu$m. Representative contrast-enhanced videos are provided in video S9.

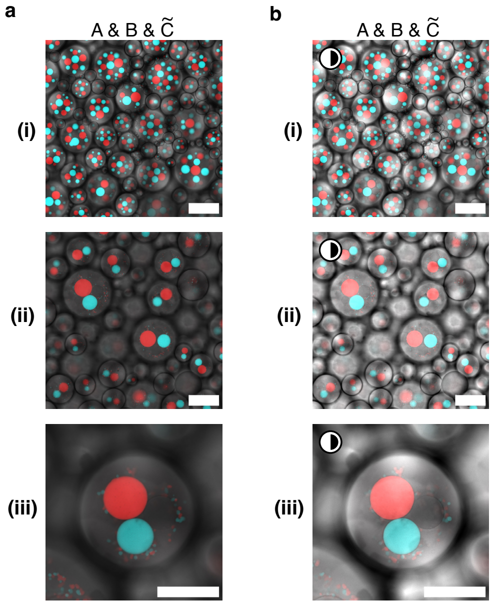


**Fig. S42 Confocal micrographs depicting formation of three distinct kinds of orthogonal RNA organelles in stoichiometric ternary systems within synthetic cells.** (a) Pristine and (b) contrast-enhanced (bright-field only, half-shaded circle) confocal micrographs (composite of bright-field/MG/DFHBI channels) representing condensates formed in bulk stoichiometric ternary systems with [A-T]:[B-T]:[C̃-T] = 1:1:1, where C̃ shares the same KL as C but lacks any fluorogenic aptamer. Micrographs in (b) were contrast-enhanced to better highlight the presence of this third non fluorescent phase. (i) Samples were imaged after more than 48 h. Every droplet contains three different kinds of condensates, one from each orthogonal construct, but their number widely varies. (ii) Samples were imaged after more than a week, allowing for even further coalescence and relaxation, as demonstrated by the reduced number of organelles within each droplet. (iii) Zoomed insets from (ii) depicting examples of synthetic cells (water-in-oil droplets) containing exactly one condensate of each kind. Micrographs in (i), (ii) have been binned down to the desired size (1024 px × 1024 px). Scale bar is 100 $\mu$m in (i)-(ii) and 50 $\mu$m in (iii).

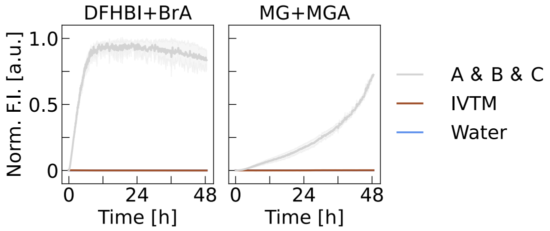


**Fig. S43 Fluorescence kinetics monitoring *in vitro* transcription and co-transcriptional folding of orthogonal A, B and C RNA nanostars ternary systems within synthetic cells.** Samples were incubated at 30 °C. Left/right column depicts normalised fluorescence intensity in the DFHBI+BrA/MG+MGA channel, respectively. Negative controls include dyes in water (Water) and dyes in *In Vitro* Transcription mixture (IVTM) with no added DNA template. Data are presented as normalised (see Methods) mean (solid/dashed lines) ± standard deviation (shaded regions) from triplicates.

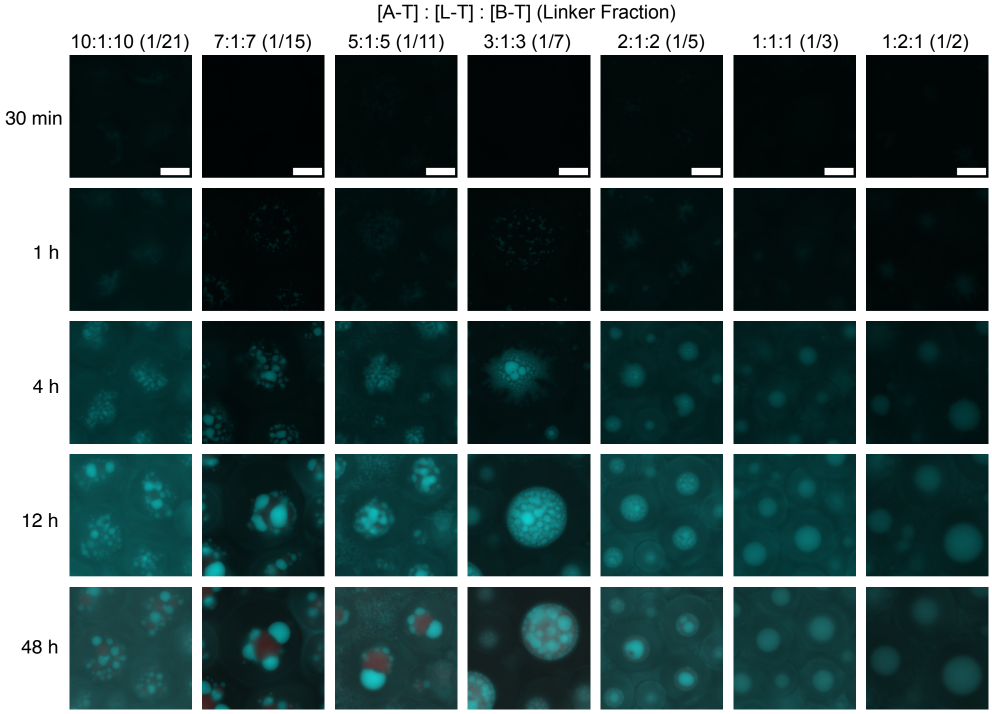


**Fig. S44 Self-assembly of single and multi-phase organelles in programmably mixed A/L/B systems within synthetic cells.** Pristine epifluorescence micrographs depicting representative timepoints (30 min, 1 h, 4 h, 12 h and 48 h, vertical axis) during the self-assembly of samples with varying DNA template proportions ([A-T]:[L-T]:[B-T] or Linker Fraction, horizontal axis). Condensates appear to stick to one another in networks at low L-T proportions (10:1:10, 7:1:7), progressively merging into single multiphasic condensates at intermediate ones, and finally completely mixing at high linker proportions (1:1:1 or 1:2:1). Samples corresponding to DNA template proportions 1:7:1 and 1:3:1 were imaged in a separate run where template mixing and encapsulation was carried out 30 min earlier compared to remaining samples. As such, the first available timepoint was used to represent 30 min for all remaining samples, while the third timepoint (acquired 30 min later) was used for 1:7:1 and 1:3:1 for ease of comparison (timestep in first 10 h = 15 min). See SI Methods section 1.4.1 and Table S6 for detailed information on the time interval between template mixing and start of timelapse imaging. All micrographs for each sample (column) show the same FOV area, thus sharing scale bars. Scale bars are 50 $\mu$m. Representative contrast-enhanced videos are provided in video S10.

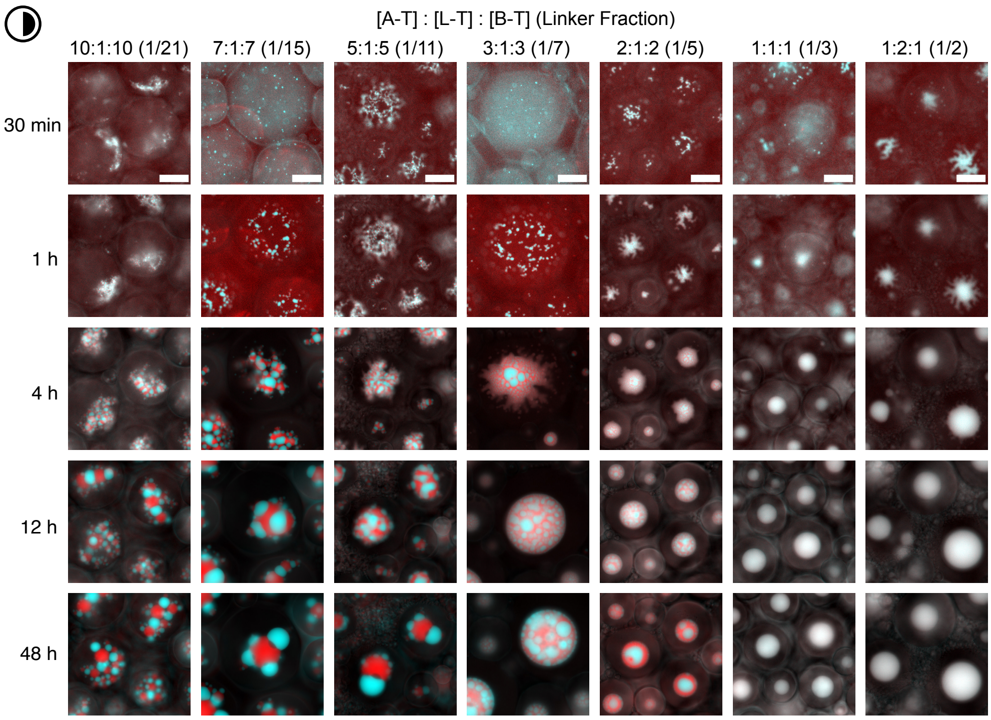


**Fig. S45 Self-assembly of single and multi-phase organelles in programmably mixed A/L/B systems within synthetic cells (processed).** Contrast-enhanced (as indicated by the half-shaded circle) epifluorescence micrographs corresponding to pristine ones in Fig. S44. Samples corresponding to DNA template proportions 1:7:1 and 1:3:1 were imaged in a separate run where template mixing and encapsulation was carried out 30 min earlier compared to remaining samples. As such, the first available timepoint was used to represent 30 min for all remaining samples, while the third timepoint (acquired 30 min later) was used for 1:7:1 and 1:3:1 for ease of comparison (timestep in first 10 h = 15 min). See SI Methods section 1.4.1 and Table S6 for detailed information on the time interval between template mixing and start of timelapse imaging. All micrographs for each sample (column) show the same FOV area, thus sharing scale bars. Scale bars are 50 $\mu$m. Representative contrast-enhanced videos are provided in video S10.

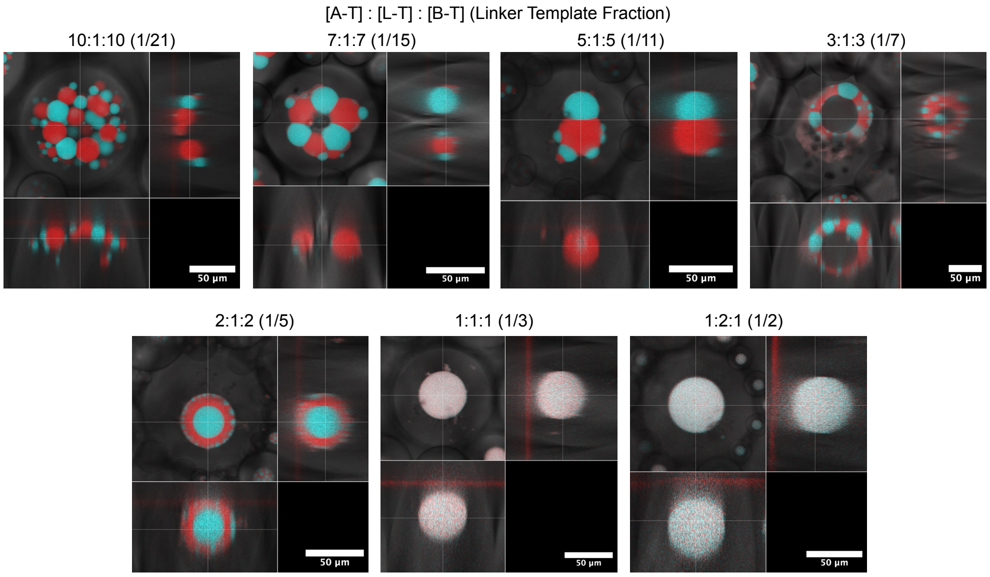


**Fig. S46 Orthogonal cross-sections depicting the 3D shape of multi- and single-phase RNA organelles in A/L/B systems within synthetic cells.** Orthogonal cross-section slices, obtained from confocal z-stacks, are reported in the following order: main panel - XY, bottom - XZ, right - YZ. Samples were imaged after more than 48 h. Micrographs are pristine composites of bright-field, MG and DFHBI channels. Organelles are located at the bottom of the enclosing synthetic cell due to sedimentation. Z-spacing was calibrated as outlined in the SI Methods, section 1.4.3. 3D renderings and clippings showcasing the internal structure of RNA organelles formed in A:L:B systems with Linker Template Fractions = 1/11, 1/7 and 1/5 are provided in video S11. A larger FOV confocal z-stack for hollow, capsule-like organelles formed in A/L/B systems with Linker Template Fraction = 1/7 is provided in video S12.

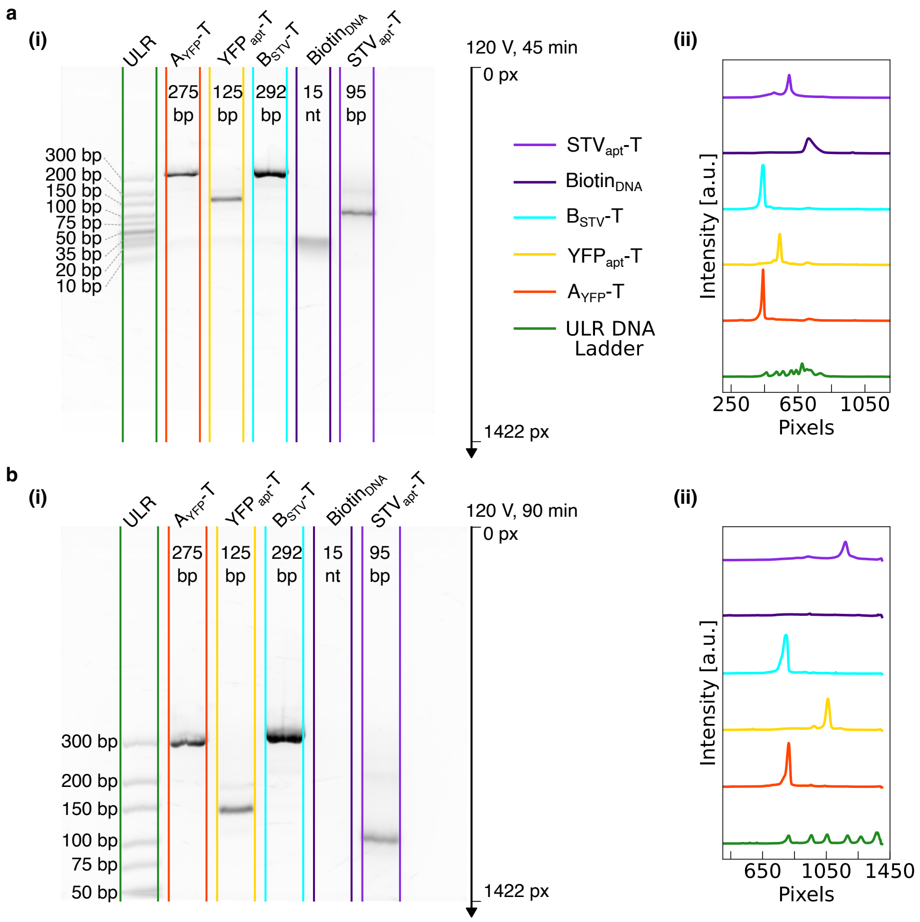


**Fig. S47 Size of PCR amplified DNA templates for protein capturing systems investigated *via* agarose gel electrophoresis** (a) (i) Agarose gel representing the electrophoretic mobilities of PCR amplified DNA templates $A_{YFP}$-T, $YFP_{apt}$-T, and $B_{STV}$-T, as well as non amplified single-stranded oligonucleotide $Biotin_{DNA}$ and double-stranded template $STV_{apt}$-T after 45 min (see SI Methods, section 1.2.3). Ultra-Low-Range (ULR) DNA ladder is used as reference in the first lane, coloured in green. (ii) Lane intensity profiles computed from gel in (a)-(i) (see SI Methods, section 1.2.3). Visual inspection of the gel in (a)-(i) and the lane profiling in (a)-(ii) confirms the expected sizes of the DNA templates coding for the designed constructs. (b)(i) Same gel in (a)-(i) after additional 45 min of electrophoresis. (ii) Lane intensity profiles computed from gel in (b)-(i). Visual inspection of the gel in (b)-(i) and the lane profiling in (b)-(ii) allows to better appreciate the migration of $A_{YFP}$-T and $B_{STV}$-T, confirming the respective bands indeed lie slightly below and around 300 bp, as expected from the respective sizes. $Biotin_{DNA}$ is no longer detectable due to having overrun the available gel length. In both (a)-(i) and (b)-(i), the length in base pairs for ladder markings is reported to the side of the corresponding bands, while the expected length of DNA templates and constructs is reported on top of the corresponding lane.

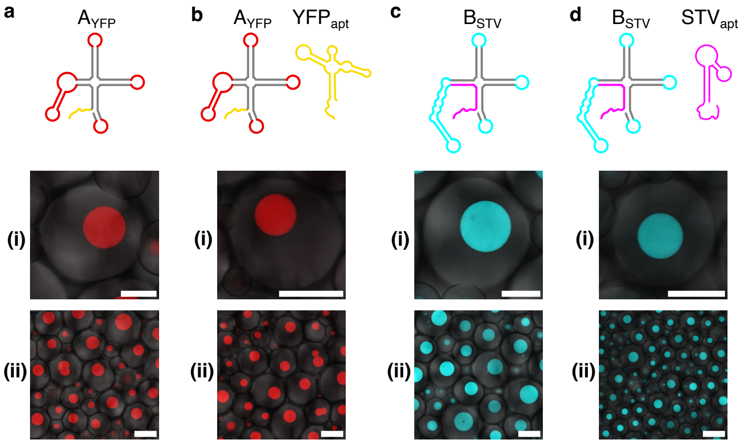


**Fig. S48 Modified protein capturing RNA nanostars form synthetic organelles within droplet-based synthetic cells in the absence of target proteins.** (a) In the absence of $\mathrm{YFP_{apt}}$, $\mathrm{A_{YFP}}$ forms single MG-rich (red) organelles within emulsion droplets, as demonstrated by (i) zoomed-in and (ii) large field-of-view confocal micrographs. (b) Same as (a) for simultaneous transcription of $\mathrm{A_{YFP}}$ and $\mathrm{YFP_{apt}}$. (c) In the absence of $\mathrm{STV_{apt}}$ $\mathrm{B_{STV}}$ forms single DFHBI-rich (cyan) organelles within emulsion droplets, as demonstrated by (i) zoomed-in and (ii) large field-of-view confocal micrographs. (b) Same as (a) for simultaneous transcription of $\mathrm{A_{YFP}}$ and $\mathrm{YFP_{apt}}$. All samples were imaged after 48 h from the start of transcription. All confocal micrographs are pristine and uncropped. Scale bars in (a)-(d) (i) are 50 $\mu$m. Scale bars in (a)-(d) (ii) are 100 $\mu$m. For experimental conditions, see SI Methods, section 1.3.6.

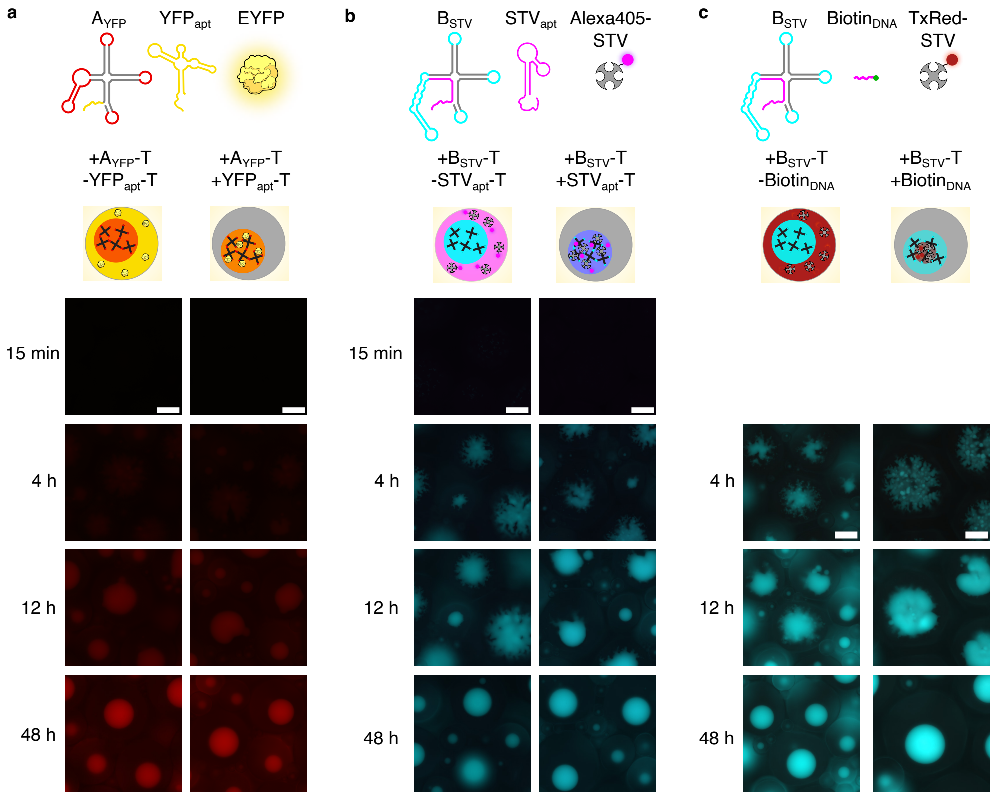


**Fig. S49 Modified RNA nanostars self-assemble into organelles capable of capturing target proteins.** Pristine epifluorescence micrographs depicting representative timepoints (15 min, 4 h, 12 h, 48 h, along vertical axis) during the self-assembly of protein capturing RNA organelles within droplet-based synthetic cells. (a) $A_{YFP}$ (red) nanostars fail to capture EYFP (yellow) in the absence of $YFP_{apt}$ (-$YFP_{apt}$-T, left), while succeed in doing so when simultaneously transcribed with it (+$YFP_{apt}$-T, right). (b) Same as (a) for $B_{STV}$ (cyan) nanostars failing to capture/capturing Alexa405-STV (magenta) in the presence/absence of the DNA template coding for the corresponding aptamer, $STV_{apt}$ ($\mp STV_{apt}$ - T, left/right). (c) Same as (b) for $B_{STV}$ (cyan) nanostars capturing TexasRed-STV (red) *via* short biotinylated DNA oligonucleotide $Biotin_{DNA}$. The timepoint at 15 min is lacking due to software-related issues in the corresponding run resulting in usable data from 3.5 h onwards only. For experimental conditions, see SI Methods, section 1.3.6. Timestamps are reported with respect to the start of timelapse imaging (see SI Methods, section 1.4.1 and Table S6). All micrographs in the same column show the same FOV area and, as such, share the same scale bar. Scale bars are 50 $\mu$m. Representative contrast-enhanced videos are provided in video S13.

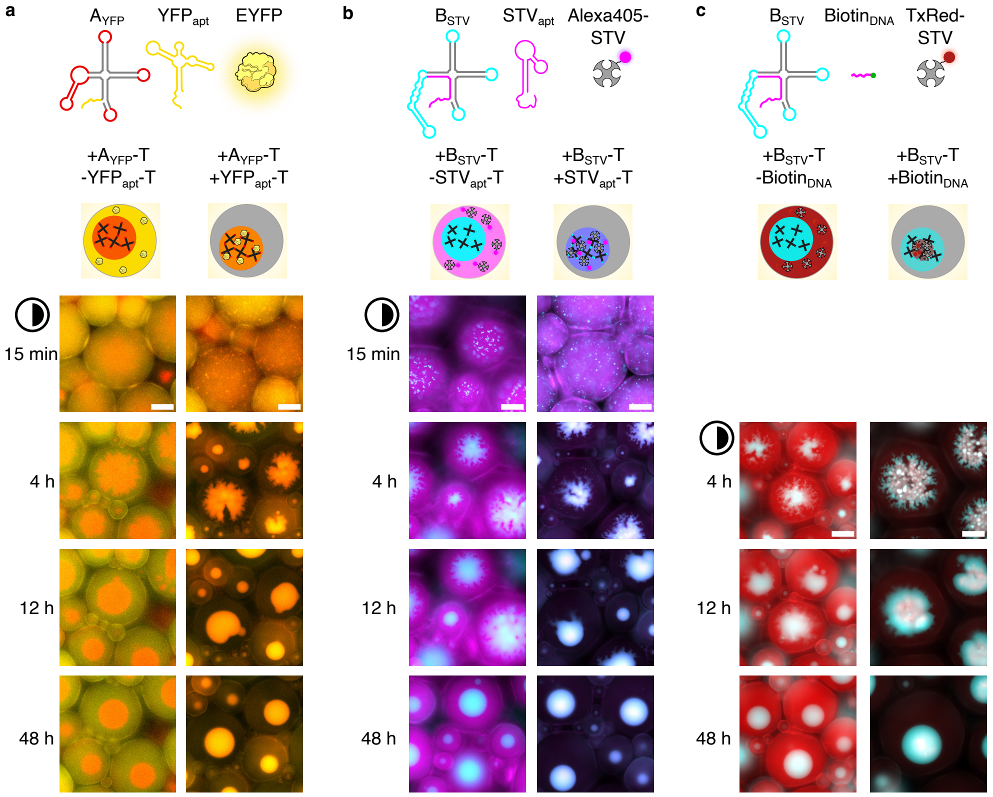


**Fig. S50 Modified RNA nanostars self-assemble into organelles capable of capturing target proteins.** Contrast-enhanced (as indicated by the half-shaded circle) epifluorescence micrographs corresponding to pristine ones in Fig. S49. Timestamps are reported with respect to the start of timelapse imaging (see SI Methods, section 1.4.1 and Table S6). All micrographs in the same column show the same FOV area and, as such, share the same scale bar. Scale bars are 50 $\mu$m. Representative contrast-enhanced videos are provided in video S13.

### 4 Supplementary Videos

**Video S1** - self-assembly of single RNA species systems in bulk
**Video S2** - melting of condensates from single RNA species in bulk
**Video S3** - self-assembly of binary RNA systems with varying DNA template ratio in bulk
**Video S4** - self-assembly of binary RNA systems with non-sticky species in bulk
**Video S5** - self-assembly of ternary RNA system in bulk
**Video S6** - self-assembly of single RNA species systems in synthetic cells
**Video S7** - self-assembly of binary RNA systems with varying DNA template ratio in synthetic cells
**Video S8** - self-assembly of binary RNA systems with non-sticky species in synthetic cells
**Video S9** - self-assembly of ternary RNA systems in synthetic cells
**Video S10** - self-assembly of RNA mixtures with programmable linker-induced mixing in synthetic cells
**Video S11** - 3D rendering and clipping showcasing the internal structure of condensates formed in A:L:B systems with Linker Template Fractions = 1/11, 1/7 and 1/5
**Video S12** - confocal Z-stack showcasing the internal structure of condensates formed in A:L:B systems with Linker Template Fraction = 1/7
**Video S13** - self-assembly of protein capturing RNA organelles
